# 2D HSQC-derived “dark forest” image with enhanced local resolution via first derivative processing–logarithmic cosine transformation (FDP–LCT): Demonstration on per-*O*-ethylated kappa- and iota-carrageenans

**DOI:** 10.1101/2025.11.21.689839

**Authors:** Xiaohui Xing, Jeffrey P. Tingley, Barinder Bajwa, Vincent Weiler, Tony Montina, Steve W. Cui, D. Wade Abbott

## Abstract

Solution-state two-dimensional (2D) ^1^H–^13^C HSQC NMR is a powerful tool for polysaccharide structure elucidation but often suffers from limited sensitivity and broad peaks due to the low natural abundance of ^13^C and poor digital resolution of the indirect dimension, respectively, as well as the typically low concentration and high viscosity of polysaccharide solutions. It is therefore pivotal to improve the resolution of 2D ^1^H–^13^C HSQC spectra for accurate peak picking and assignment, particularly in the indirect ^13^C dimension. In this study, we developed an algorithm that combines first derivative processing with a novel logarithmic cosine transformation (FDP–LCT) to convert 2D ^1^H–^13^C HSQC spectra into local-resolution-enhanced images resembling a dark forest of straight, densely standing trees. These images revealed sharpened spectral features and enabled extraction of precise ^1^H and ^13^C chemical shifts, as demonstrated using per-*O*-ethylated kappa-and iota-carrageenans, two sulfated galactans differing only by a single substitution at the *O*-2 position of anhydrogalactose. In conclusion, this approach provides an effective post-acquisition strategy for enhancing digital resolution in 2D HSQC spectra and improving the structural analysis of closely related complex polysaccharides.

**Graphic abstract:** 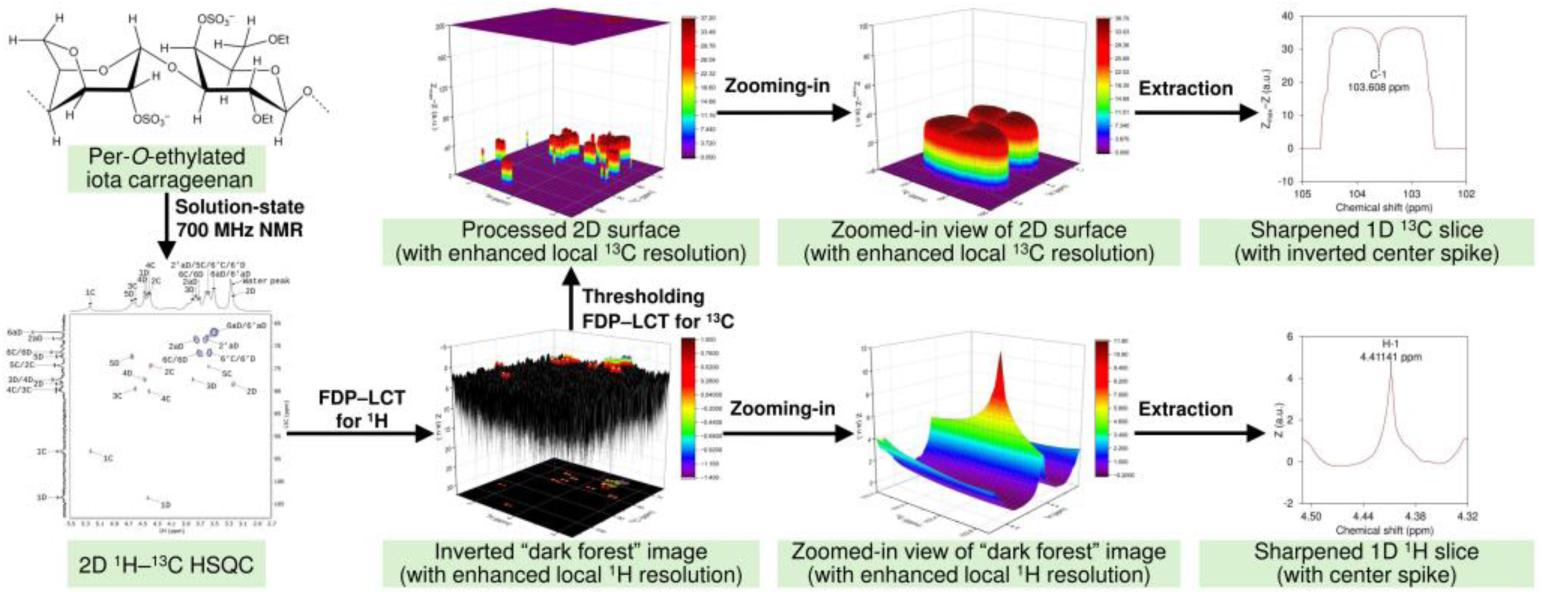

**Highlights:**

- An algorithm combining first derivative processing and logarithmic cosine transformation was developed for 2D HSQC.
- The algorithm was used to boost the local resolution of 2D HSQC spectra of the per-*O*-ethylated carrageenans.
- Structures were distinguished by complete interpretation of 2D NMR spectra.
- A post-processing workflow was developed to facilitate chemical shift extraction from sharpened HSQC.

## 1. Introduction

Solution-state two-dimensional (2D) NMR spectroscopy has become indispensable for the structural elucidation of complex carbohydrates, as it alleviates the severe resonance overlap that limits one-dimensional (1D) approaches and provides a more reliable framework for signal assignment [1–4]. Among these, 2D ^1^H–^13^C heteronuclear single quantum coherence (HSQC) spectroscopy is particularly valuable because it directly correlates protons with their covalently bonded carbons, yielding clearer spectra with reduced overlap [5, 6]. This property makes 2D ^1^H–^13^C HSQC well suited for purified oligo- and polysaccharide fractions, and for more complex, unfractionated samples (*e.g.*, whole cell wall) where spectral complexity is extreme [6–15]. Nevertheless, sensitivity limitations arising from low ^13^C natural abundance, the typically low sample concentrations employed to reduce viscosity, and viscosity-induced line broadening often diminish signal-to-noise and complicate peak deconvolution [16, 17]. In the indirect ^13^C dimension of 2D HSQC, peak broadening is especially pronounced in polysaccharide analysis, limiting spectral resolution [3, 18]. Improvements in pulse sequences and acquisition efficiency have advanced 2D HSQC by enhancing resolution, preserving spectral quality, and enabling efficient analysis of complex systems [19–22]. Software-based post-processing has also emerged as a complementary strategy, where advanced algorithms applied to processed 2D HSQC spectra disentangle overlapping signals, sharpen cross-peaks, and improve the extraction of chemical shifts and other spectral features [23–26].

Carrageenans are a group of polysaccharides from red algae, consisting of alternating β-(1→4)-linked D-galactopyranose and α-(1→3)-linked residues that are either D-galactopyranose or 3,6-anhydro-D-galactopyranose, with this variation together with the degree and position of sulfation determining the structural types, among which kappa- and iota-carrageenans are two of the common forms [27, 28]. Carrageenans are widely used in the food, pharmaceutical, and biotechnological industries as functional additives and materials owing to their viscosity-enhancing properties, ion-sensitive gelation, and synergistic interactions with proteins and other polysaccharides [29–33]. Kappa-carrageenan consists of repeating β-D-galactopyranose-4-sulfate and 3,6-anhydro-α-D-galactopyranose, whereas iota-carrageenan carries an additional sulfate group at the *O*-2 position of the 3,6-anhydro-α-D-galactopyranose residue [28, 33, 34], as shown in Scheme 1. Despite differing by only this single sulfate substitution, kappa- and iota-carrageenans exhibit markedly different properties: kappa-carrageenan has lower water solubility than iota-carrageenan, requiring heating to dissolve in water, and forms rigid gels upon cooling or in the presence of K^+^, and they also differ in synergistic interactions with proteins (*e.g.*, casein), leading to distinct effects on gel strength, elasticity, and microstructure [29, 31, 32, 35–37]. However, the strong tendency of carrageenans, especially kappa-carrageenan, to interact with cations and proteins makes unfractionated and crude samples prone to complex formation, thereby impairing their solubility and rehydration properties in aqueous systems. This limitation renders repeated freeze-drying in D2O particularly challenging for solution-state 1D and 2D NMR investigations of carrageenans in crude water extracts and unfractionated cell walls (*e.g.*, alcohol-insoluble residues) of red seaweeds.

Alkylation strategies such as per-*O*-methylation and per-*O*-ethylation have been widely used for GC–MS-based glycosidic linkage analysis of purified polysaccharide fractions [38–54] and unfractionated mixtures such as whole cell wall samples from higher plants [55, 56], seaweeds [57, 58], and fungi [59]. High-resolution 2D NMR, particularly 2D ^1^H–^13^C HSQC, of per-*O*-alkylated samples dissolved in organic solvents has proven valuable for characterizing water-insoluble unfractionated polysaccharides, as demonstrated in higher plant cell walls [10], yet this approach has not been applied to seaweed samples. For analysis of sulfated galactans, per-*O*-methylation– or per-*O*-ethylation–GC–MS analysis has been widely applied to resolve linkage patterns as well as the degree and position of sulfation when combined with desulfation analysis; notably, per-*O*-ethylation provides the additional advantage of determining the degree and position of natural *O*-methylation [40, 41, 43–45, 52–54]. However, to date, no study has reported the application of per-*O*-ethylation–NMR analysis to carrageenans.

In this study, we developed an algorithm combining first derivative processing with logarithmic cosine transformation (FDP–LCT) to enhance the local resolution of 2D ^1^H–^13^C HSQC spectra, sharpening ^1^H and ^13^C peaks and thereby facilitating chemical shift extraction. We also demonstrate that per-*O*-ethylation–NMR analysis yielded high-quality 2D ^1^H–^13^C HSQC and other 2D NMR spectra, enabling complete discrimination and assignment of signals from per-*O*-ethylated kappa- and iota-carrageenans, whose structures differ by only a single *O*-2 substitution on anhydrogalactose (Scheme 1).

**Scheme 1.**
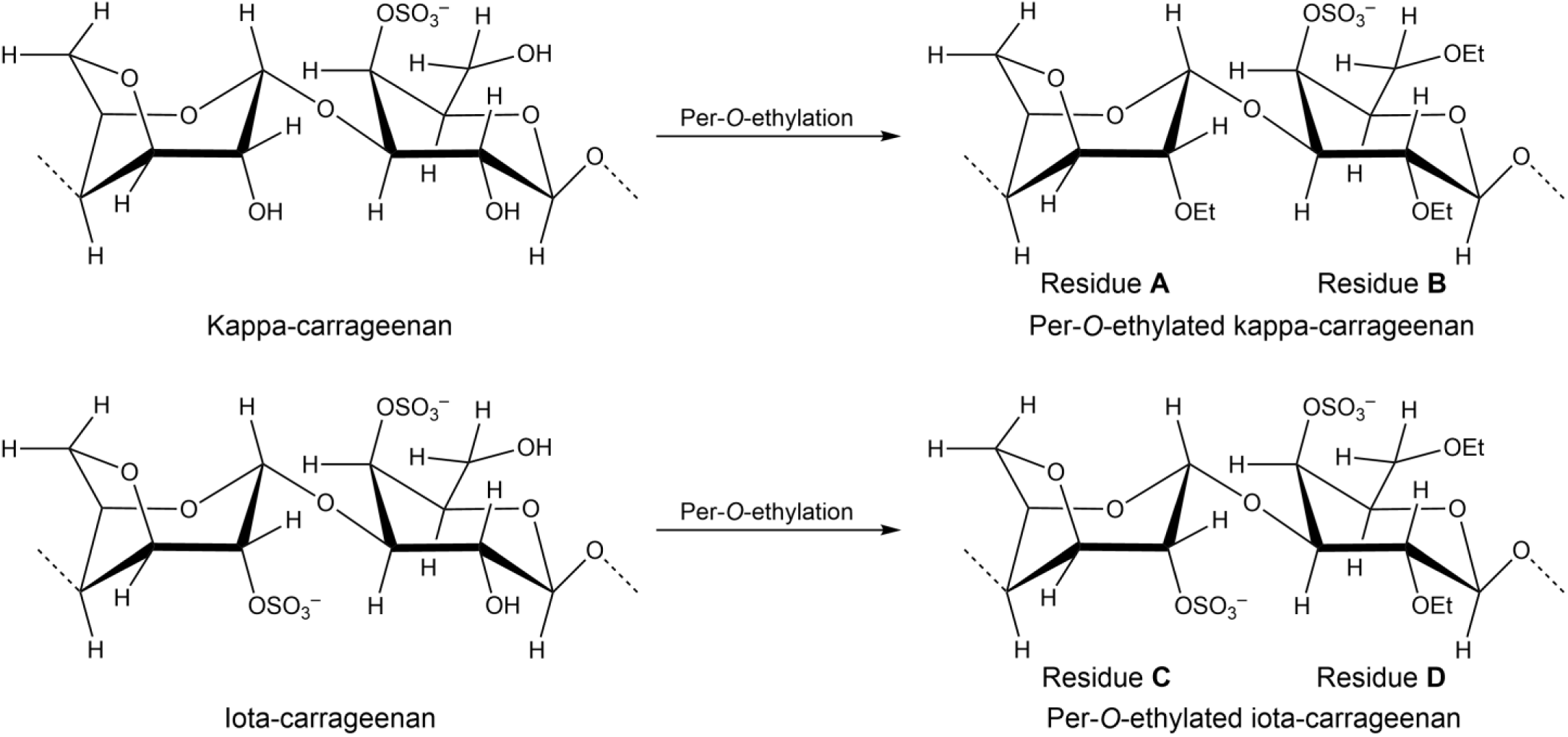
Structures of the disaccharide repeating units of kappa- and iota-carrageenans and their per-*O*-ethylated derivatives. Residues A and B represent 4-linked 2-*O*-ethyl-3,6-anhydro-α-D-galactopyranose and 3-linked 2,6-di-*O*-ethyl-4-*O*-sulfo-β-D-galactopyranose in per-*O*-ethylated kappa-carrageenan, respectively. Residues C and D represent 4-linked 2-*O*-sulfo-3,6-anhydro-α-D-galactopyranose and 3-linked 2,6-di-*O*-ethyl-4-*O*-sulfo-β-D-galactopyranose in per-*O*-ethylated iota-carrageenan, respectively. In all NMR spectra presented in this study, residues A–D are used to designate the monosaccharides within the corresponding disaccharide structures of the per-*O*-ethylated carrageenans.

## 2. Theory

Derivative values (*e.g.*, those from first derivative spectroscopy) can be converted into corresponding 𝑍 values defined by the equation below:

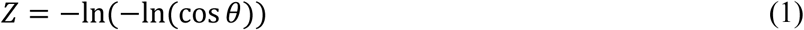

where 𝜃 is defined as the tangential angle in the range of -π/2 to π/2, and 𝜃 can be expressed in tangent function form as:

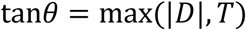

or in inverse tangent function form as:

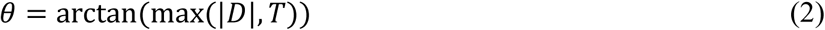

In the above equations, 𝐷 represents derivative value, and |𝐷| denotes the absolute value of 𝐷. 𝑇 is a selected threshold, an extremely small positive value, close to zero but not equal to zero. The purpose of introducing 𝑇 is to avoid tan𝜃 = 0 and cos𝜃 = 1, which would render 𝑍 in Equation (1) undefined (+∞), and to provide 𝑍 with a defined possible maximum when tan𝜃 = 𝑇.

By combining Equations (1) and (2), the following is obtained:

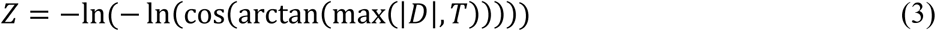

Solving the trigonometric functions in Equation (3) results in:

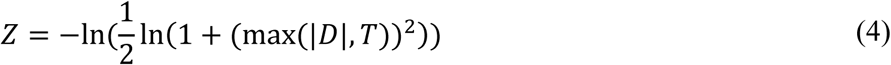

A 𝑇 value of 1.1 × 10^-8^ is recommended to prevent small derivative values from being rounded to zero during logarithmic calculations by common software (*e.g.*, Microsoft Excel). This 𝑇 value allows for a possible maximum 𝑍 value of 36.7368005696771. Substituting this 𝑇 value into Equation (4) gives the practical equation:

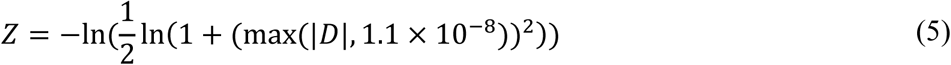

As a theoretical example, when Equation (5) is applied to the first derivative (d𝑦⁄d𝑥) values (Scheme 2B) of a 1D Gaussian peak (Scheme 2A) in the 𝑥𝑦 coordinate system, the resulting plot (Scheme 2C) shows a sharpened central spike at the peak center, corresponding to the maximum point of the original Gaussian peak (Scheme 2A) and to the zero-crossing of its first derivative (Scheme 2B), together with symmetric valleys corresponding to the inflection points of the original curve (Scheme 2A) and the extrema of its derivative (Scheme 2B). In a second theoretical example, Equation (5) was applied to the first partial-derivative (∂𝑧⁄∂𝑥) values (Scheme 3B) of a 2D Gaussian peak (Scheme 3A) in the 𝑥𝑦𝑧 coordinate system, yielding an inverted surface (Scheme 3C) with a wedge-like central notch of enhanced resolution, whose projection along the *y*-axis (Scheme 3D) shows a sharpened central spike with symmetric valleys, while its projection along the *x*-axis (Scheme 3E) lacks these spike–valley features.

**Scheme 2.**
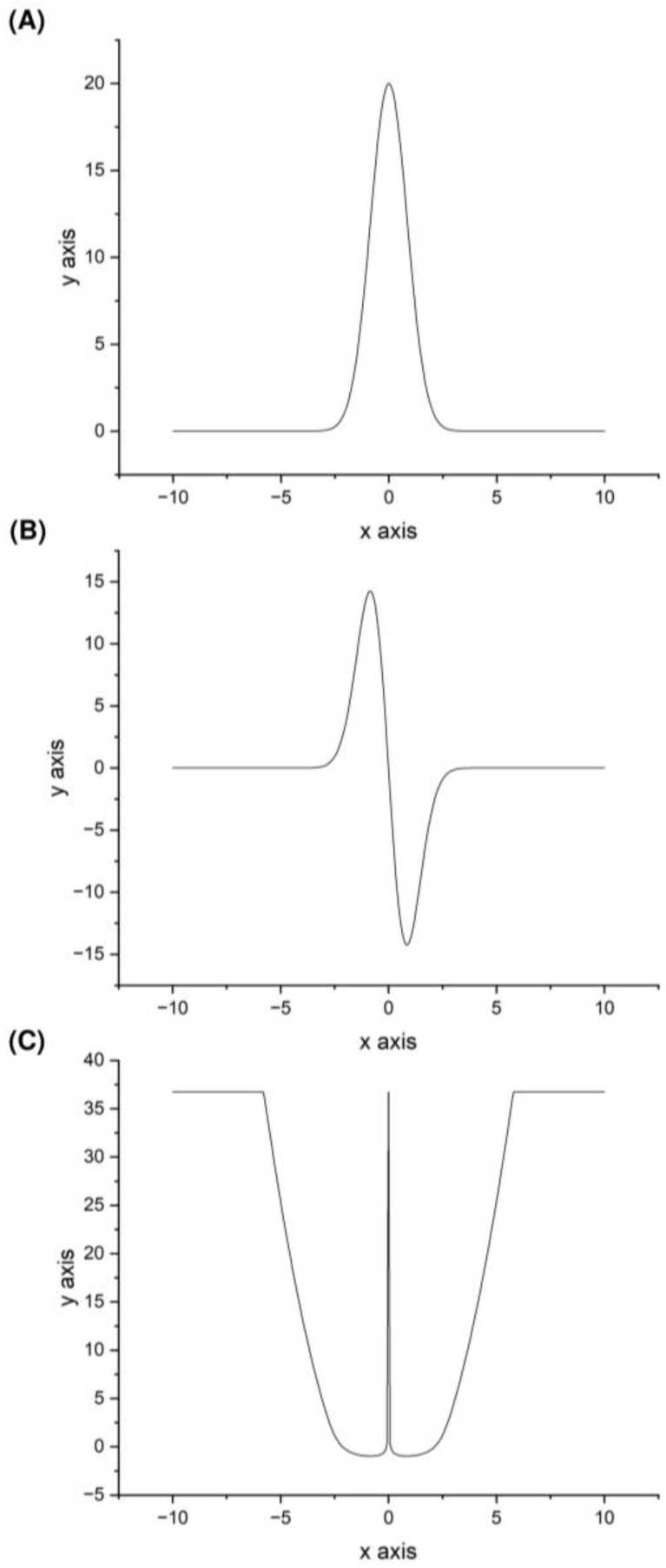
(A) 1D Gaussian peak, 𝑦 = 20e^−𝑥2ln2^, with a peak height of 20 and a full width at half maximum (FWHM) of 2; (B) the first derivative (d𝑦⁄d𝑥) of the Gaussian function, showing the characteristic antisymmetric slope profile with a zero-crossing point corresponding to the peak center and positive and negative extrema corresponding to the inflection points of the original curve (A); and (C) the logarithmic cosine transform of the first derivative, revealing enhanced local resolution of the central spike originating from the peak center of the original plot (A), along with symmetric valley features corresponding to the inflection points of plot (A) and the extrema of plot (B).

**Scheme 3.**
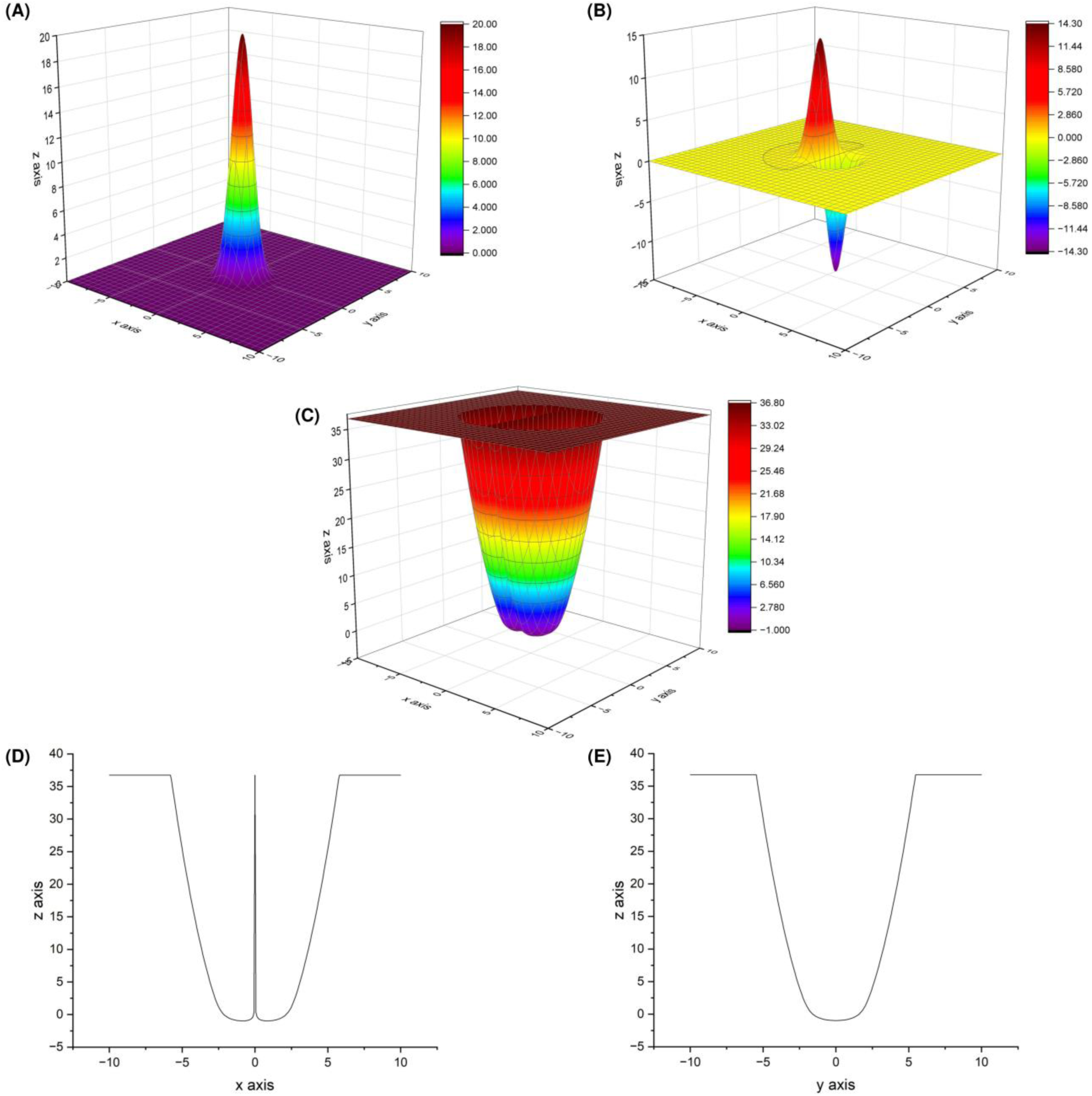
(A) 2D Gaussian peak, 𝑧 = 20e^−(𝑥2+𝑦2)ln2^, with a peak height of 20 and an FWHM of 2 along both the *x*- and *y*-axes; (B) the first partial-derivative (∂𝑧⁄∂𝑥) of the original function, showing the characteristic antisymmetric ridge–valley profile along the *x*-axis; (C) the logarithmic cosine transform of the first partial-derivative values, illustrating an inverted peak with a resolution-boosted wedge-like central notch; (D) projection of the panel (C) surface along the *y*-axis, showing the central spike and symmetric valleys; and (E) projection of the panel (C) surface along the *x*-axis, lacking the central spike and valley features.

## 3. Materials and methods

### 3.1. Materials and reagents

In the present study, three carrageenan samples were utilized: two commercial standards and one in-house prepared fraction. The commercial iota-carrageenan (catalogue No. YC30038) and kappa-carrageenan (catalogue No. YC30039) were procured from Biosynth Carbosynth USA (San Diego, CA, USA). The carrageenan fraction (designated F60) was prepared in-house from *Mazzaella japonica*, a red seaweed belonging to Gigartinaceae, a family known to produce kappa- and iota-carrageenans [57, 60–62], by hot water extraction of the alcohol-insoluble residue, followed by purification with thermostable α-amylase treatment and gradient ethanol precipitation. A comprehensive, detailed description of the *Mazzaella japonica* sample collection, the methodologies for extraction, purification, and compositional analysis of F60, and the chemicals, reagents, and consumables employed in this study is provided in the Supplementary Materials and Methods.

### 3.2. Per-O-ethylation of carrageenans

Carrageenan sample (∼10 mg) was magnetically stirred overnight at 70 °C in 10 mL of deionized water, followed by 24 h of dialysis with molecular weight cut-off (MWCO) of 6,000–8,000 Da against 4 L of 0.1 M triethylammonium chloride, then another 24 h of dialysis against 4 L of deionized water, and freeze-dried [57]. The dry sample was suspended in 2 mL of dimethyl sulfoxide (DMSO) by magnetic stirring overnight at room temperature. After adding 200 mg of sodium hydroxide powder and magnetic stirring for 15 min, 1.2 mL of ethyl iodide was added, the tube resealed, and magnetically stirred for 24 h at room temperature in the dark [15]. The product was partitioned between 3 mL of dichloromethane and 3 mL of deionized water three times. The lower phase was evaporated to dryness under nitrogen. The pooled upper phases were neutralized with acetic acid, evaporated under nitrogen to reduce the volume by half, and then combined with the dried lower phase. The mixture was dialyzed (MWCO 6,000–8,000 Da) overnight in running water, then for 24 h in 4 L of deionized water, and then freeze-dried [57, 63]. The dry sample was redissolved in 2 mL of DMSO, followed by the same ethylation reaction and cleanup through partitioning, dialysis, and freeze-drying as described above. The ethylation process was then repeated once. For each of the kappa-carrageenan, iota-carrageenan, and F60 fraction isolated from *Mazzaella japonica*, three independent experiments were performed, and the resulting per-*O*-ethylated products for each sample were pooled for NMR analysis.

### 3.3. Solution-state 1D and 2D NMR analysis of per-O-ethylated carrageenans

The pooled per-*O*-ethylated sample was freeze-dried twice with 1 mL of D2O (99.9% atom D, Product No. 151882, Sigma-Aldrich) and once with a mixture of 1 mL of D2O and 50 μL of D2O containing 0.75% (w/w) sodium 3-trimethylsilyl-2,2,3,3-tetradeuteriopropionate (TSP-d4, 99.9% atom D, Product No. 293040, Sigma-Aldrich). The dry sample was dissolved in 1 mL of DMSO-d6 (99.96% atom D, Product No. 156914, Sigma-Aldrich), followed by transferring 0.68 mL of the solution to a 5-mm NMR tube. All 1D and 2D NMR experiments were performed at 323 K on a Bruker Avance III HD 700 MHz spectrometer (Bruker BioSpin GmbH, Rheinstetten, Germany), operating at 700.44 and 176.14 MHz for ^1^H and ^13^C NMR analyses, respectively. Measurements were carried out using a triple-resonance TBO-Z probe with the outer coil tuned to ^1^H and the inner coil to ^13^C. ^1^H and ^13^C chemical shifts were internally referenced to TSP-d4 at 0 ppm. 1D ^1^H, 1D ^13^C, and 2D ^1^H–^13^C HSQC spectra were collected using standard Bruker pulse sequences zg30, zgpg30, and hsqcedetgpsp.3, respectively. In addition, ^1^H–^1^H COSY (cosygpmfqf), ^1^H–^1^H TOCSY (dipsi2gpphzs), and ^1^H–^13^C HMBC (hmbcetgpl3nd) spectra were acquired to facilitate spectral interpretation. Data were processed and analyzed using Mnova software version 14.2.1-27684 (Mestrelab Research S.L., Santiago de Compostela, Spain). Comprehensive, detailed acquisition parameters for each sample and experiment are provided in the Supplementary Materials and Methods.

### 3.4. FDP–LCT processing of 2D ^1^H–^13^C HSQC spectra

The 2D matrix data in the CSV file exported from Mnova-processed 2D ^1^H–^13^C HSQC spectra of per-*O*-ethylated kappa- and iota-carrageenans were subjected to the following processing steps:

**Step 1.** A 1D slice along the ^1^H dimension for each ^13^C chemical shift value (F1 point) is subjected to FDP–LCT, yielding a 2D matrix of transformed values (denoted as 𝑍 values) that resembles a dark forest of straight, densely standing trees, which we refer to as a “dark forest image” in this study.

**Step 2.** Each 𝑍 value obtained in Step 1 is processed using the operation max(Z_thresh_ − 𝑍, 0), where Z_thresh_ is an empirically optimized threshold used for signal selection. This operation generates a 2D matrix with positive values corresponding to the retained (target) signals and zero values representing the filtered (screened-out) signals.

**Step 3.** Each 1D slice along the ^13^C dimension of the matrix obtained in Step 2 is subjected to FDP–LCT.

**Step 4.** The resulting new 𝑍 values are then processed using the operation Z_max_ − 𝑍, where Z_max_ is a constant set as 36.7368005696771, as detailed in Section 2 (Theory). This transformation keeps the processed target signals positive while maintaining the background at zero.

**Step 5.** Each contour signal is zoomed into, and the corresponding ^13^C projection is extracted from the selected region of the processed 2D matrix obtained in Step 4. To minimize artifacts arising from peak overlap at the contour boundaries, the projection is calculated as the average of the longest continuous columns of nonzero values located near the contour center along the ^13^C dimension. For sugar-ring signals, the central five of these columns are used, whereas for the sharper methyl and methylene signals of the ethyl substituent, the central three are used. From each resulting 1D ^13^C projection, the corresponding ^13^C chemical shift of the cross-peak is determined from the sharp inverse spike (arising from local resolution enhancement) near the peak midpoint.

**Step 6.** Once the ^13^C chemical shift of each cross-peak is determined in Step 5, the corresponding region in the dark forest image generated in Step 1 is zoomed into, and the 1D ^1^H slice is extracted by selecting the rows of 𝑍 values corresponding to that specific ^13^C chemical shift. From this slice, the sharp spike (arising from local resolution enhancement at the original peak center) enables accurate determination of the ^1^H chemical shift of the cross-peak.

In the above Steps 1 and 3, first-derivative analysis with built-in Savitzky–Golay smoothing of 1D ^1^H and ^13^C slices from the processed 2D HSQC data of per-*O*-ethylated kappa- and iota-carrageenans was performed in OriginPro 2023 (64-bit), version 10.0.0.154 (OriginLab Corp., Northampton, MA, USA), with a polynomial order of 2 optimized for smoothing. For the upfield region (0.20000–1.99700 ppm for ^1^H and 7.00000–27.0290 ppm for ^13^C), corresponding to cross-signals from protons and carbons in the methyl groups of ethyl substituents, smoothing windows of 3 points along the ^1^H dimension and 20 points along the ^13^C dimension were used for per-*O*-ethylated kappa-carrageenan, and 5 and 20 points for per-*O*-ethylated iota-carrageenan, respectively. For the relatively downfield region (2.20000–6.19811 ppm for ^1^H and 55.0000–114.996 ppm for ^13^C), which contains cross-signals from protons and carbons in sugar rings as well as from the methylene groups of ethyl substituents, smoothing was applied using windows of 30 points along the ^1^H dimension and 20 points along the ^13^C dimension for both per-*O*-ethylated carrageenan standards.

The LCT conversion of the first-derivative data in Steps 1 and 3, as well as the operations max(Z_thresh_ − 𝑍, 0) in Step 2 and Z_max_ − 𝑍 in Step 4 applied to the corresponding converted values, were carried out in Microsoft Excel for Microsoft 365 MSO (64-bit), version 2507 (Microsoft Corp., Redmond, WA, USA). All 3D colormap surface plots of the 2D matrices obtained in Steps 1–4 were generated in OriginPro. Generation of 1D projections and extraction of row and column slices from selected regions of the 2D matrices (Steps 5 and 6) were also performed in Microsoft Excel. All software operated under Microsoft Windows 11 Enterprise, version 10.0.22631 (Microsoft Corp., Redmond, WA, USA), on a Dell Precision 7750 mobile workstation (Dell Inc., Round Rock, TX, USA) equipped with an Intel Core i9-10885H CPU, an NVIDIA Quadro RTX 3000 GPU, 64 GB of RAM, and a 1 TB Micron 2300 NVMe SSD.

## 4. Results

Complete interpretation was achieved for the 2D ^1^H–^13^C HSQC spectra (Figs. 1 and 2), together with COSY, TOCSY, and HMBC spectra (Figs. S1–S7), of per-*O*-ethylated kappa-carrageenan and iota-carrageenan, with cross-peak assignments mutually validated and confirmed by full agreement among all 2D NMR datasets. In addition, the 1D ^1^H and ^13^C spectra were assigned with the aid of the 2D NMR correlations, and the corresponding 1D traces were stacked alongside the relevant 2D NMR spectra to facilitate direct comparison (Figs. 1, 2, and S1–S7). The HSQC datasets of per-*O*-ethylated kappa-carrageenan (Fig. 1) and iota-carrageenan (Fig. 2) were subsequently subjected to the procedure outlined in Section 3.4 (Steps 1–6). The figures generated at each step for selected regions of the 2D HSQC data of per-*O*-ethylated iota-carrageenan (Figs. 3–15) are presented in the main text as representative examples of the methodology, whereas the remaining figures for per-*O*-ethylated iota-carrageenan and all figures generated for per-*O*-ethylated kappa-carrageenan (Figs. S8–S25) are provided in the Supplementary Figures and Tables.

**Fig. 1.**
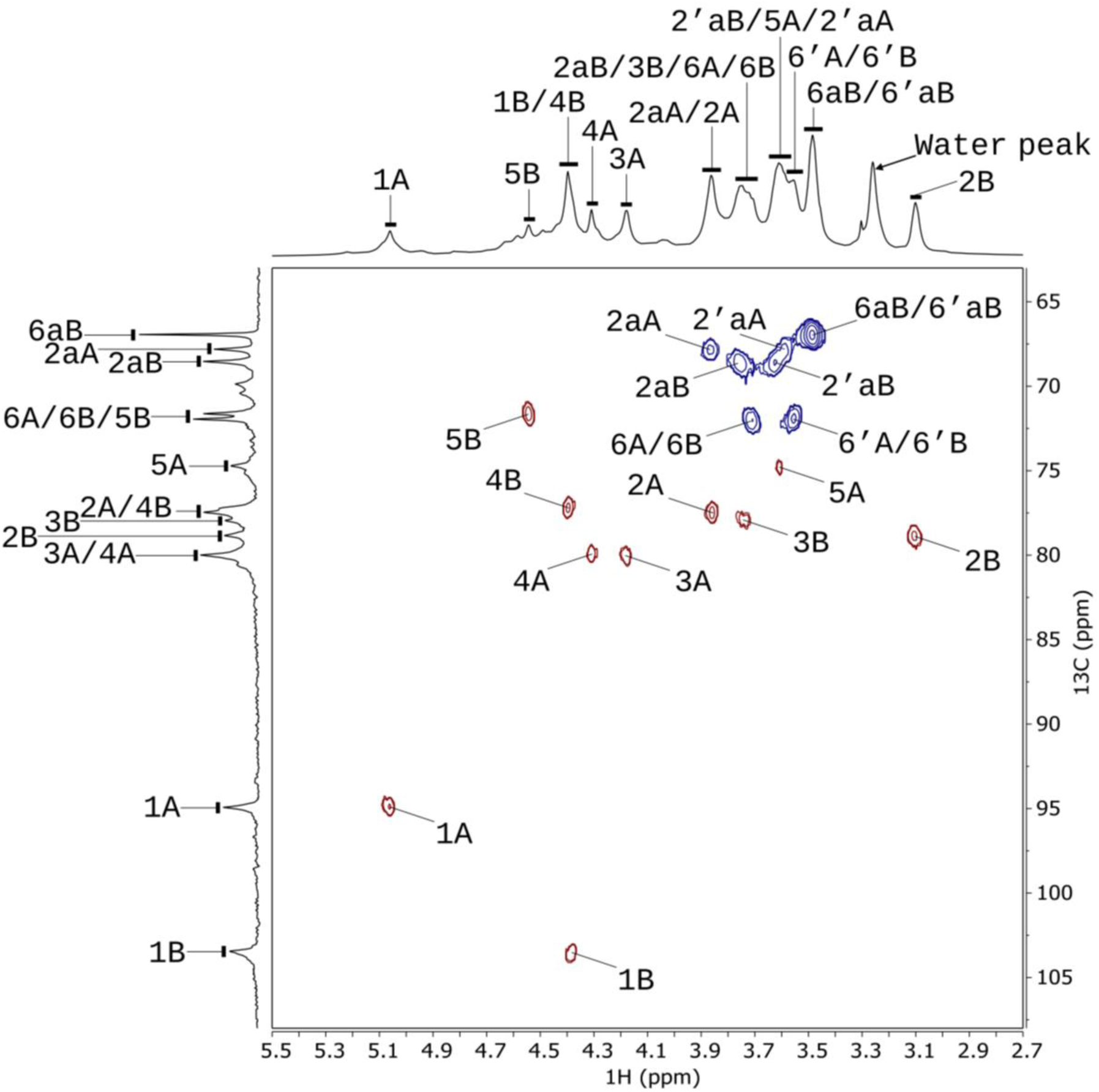
2D ^1^H–^13^C HSQC NMR spectrum (700 MHz, 323 K) of DMSO-d6 solution of per-*O*-ethylated commercial kappa-carrageenan. Residues A and B represent 4-linked 2-*O*-ethyl-3,6-anhydro-α-D-galactopyranose and 3-linked 2,6-di-*O*-ethyl-4-*O*-sulfo-β-D-galactopyranose in per-*O*-ethylated kappa-carrageenan, respectively. Cross peaks corresponding to correlations between directly bonded carbons and protons (one bond away) are marked. For example, the cross-peak “1A” refers to the correlation between C-1 and H-1 of residue A. The lowercase “a” indicates a signal from the methylene group of the ethyl group, and the number before it indicates the location of the *O*-ethyl group on the sugar ring. Signals separated by the solidus (/) symbol indicate overlapping. For example, “6A/6B” refers to the overlapping of signals 6A and 6B. ^1^H and ^13^C chemical shifts were internally referenced to TSP-d4 as 0 ppm.

**Fig. 2.**
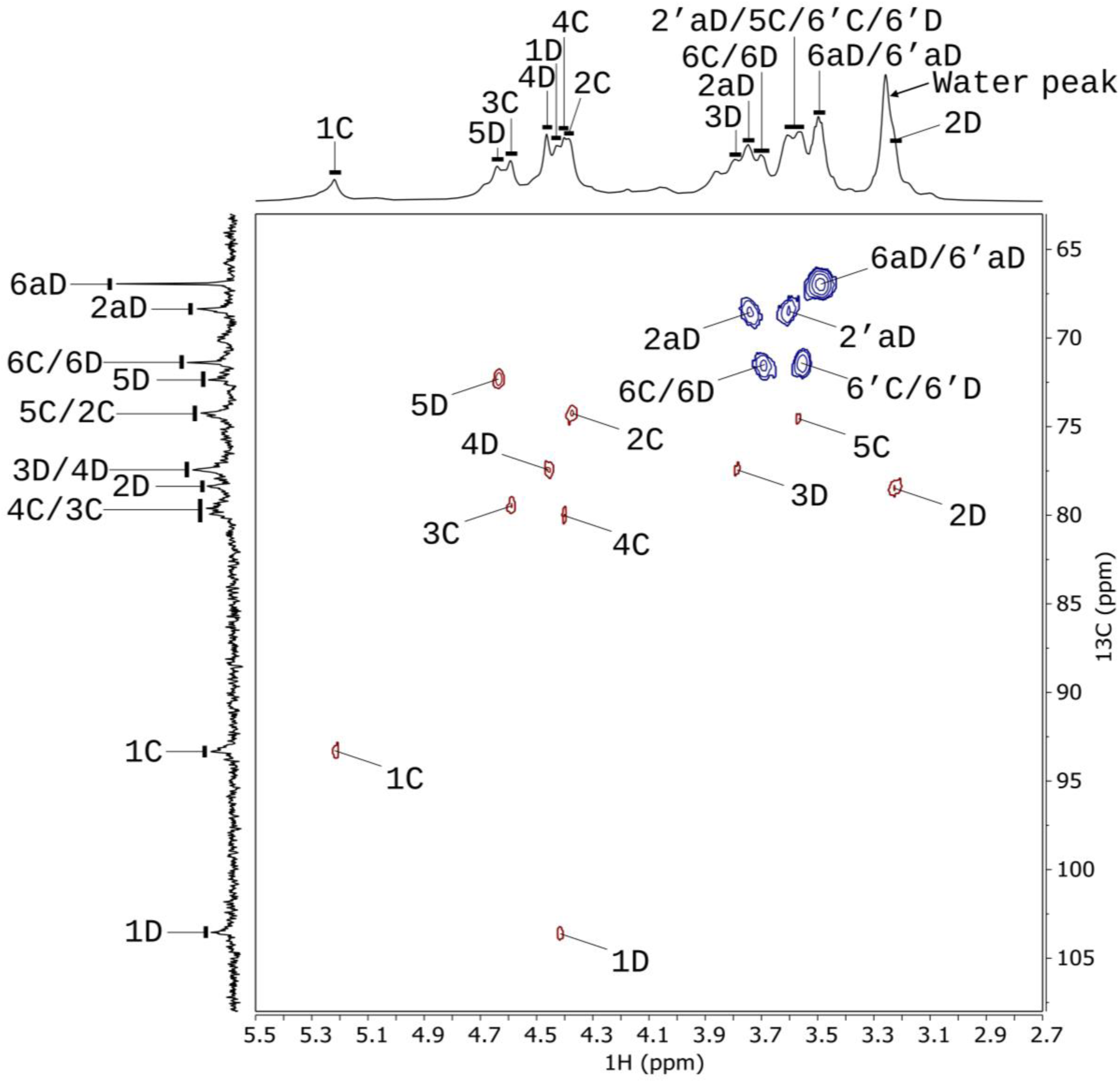
2D ^1^H–^13^C HSQC NMR spectrum (700 MHz, 323 K) of DMSO-d6 solution of per-*O*-ethylated commercial iota-carrageenan. Residues C and D represent 4-linked 2-*O*-sulfo-3,6-anhydro-α-D-galactopyranose and 3-linked 2,6-di-*O*-ethyl-4-*O*-sulfo-β-D-galactopyranose in per-*O*-ethylated iota-carrageenan, respectively. Cross peaks corresponding to correlations between directly bonded carbons and protons (one bond away) are marked. For example, the cross-peak “1C” refers to the correlation between C-1 and H-1 of residue C. The lowercase “a” indicates a signal from the methylene group of the ethyl group, and the number before it indicates the location of the *O*-ethyl group on the sugar ring. Signals separated by the solidus (/) symbol indicate overlapping. For example, “6C/6D” refers to the overlapping of signals 6C and 6D. ^1^H and ^13^C chemical shifts were internally referenced to TSP-d4 as 0 ppm.

**Fig. 3.**
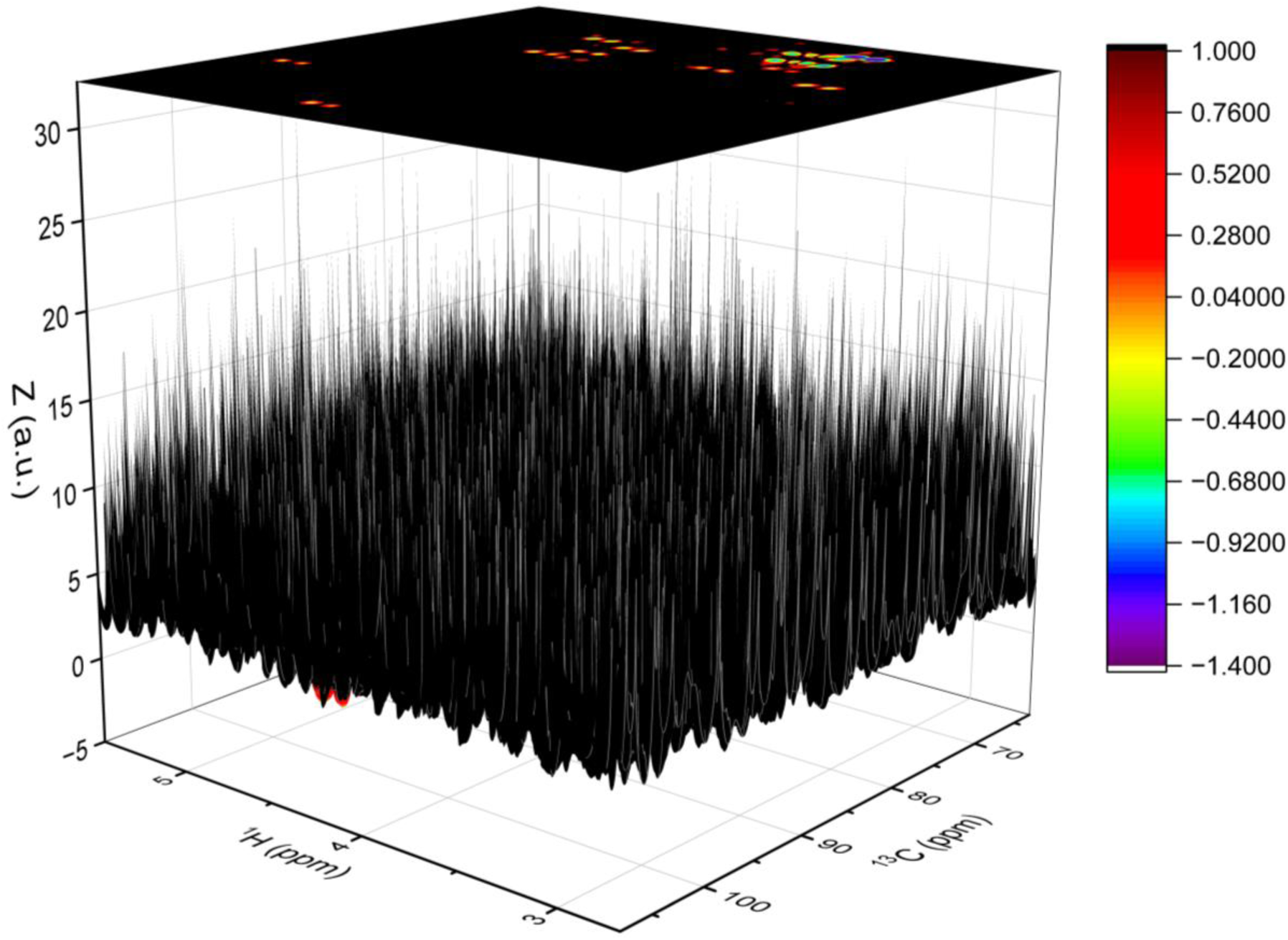
Side-view dark forest image (3D colormap surface plot) of 2D ^1^H–^13^C HSQC data of per-*O*-ethylated iota-carrageenan after FDP–LCT along the ^1^H dimension, with the surface generated from 𝑍 values corresponding to ^1^H and ^13^C chemical shifts, the color scale displayed on the right, and the corresponding 2D contour map projection shown above.

**Fig. 4.**
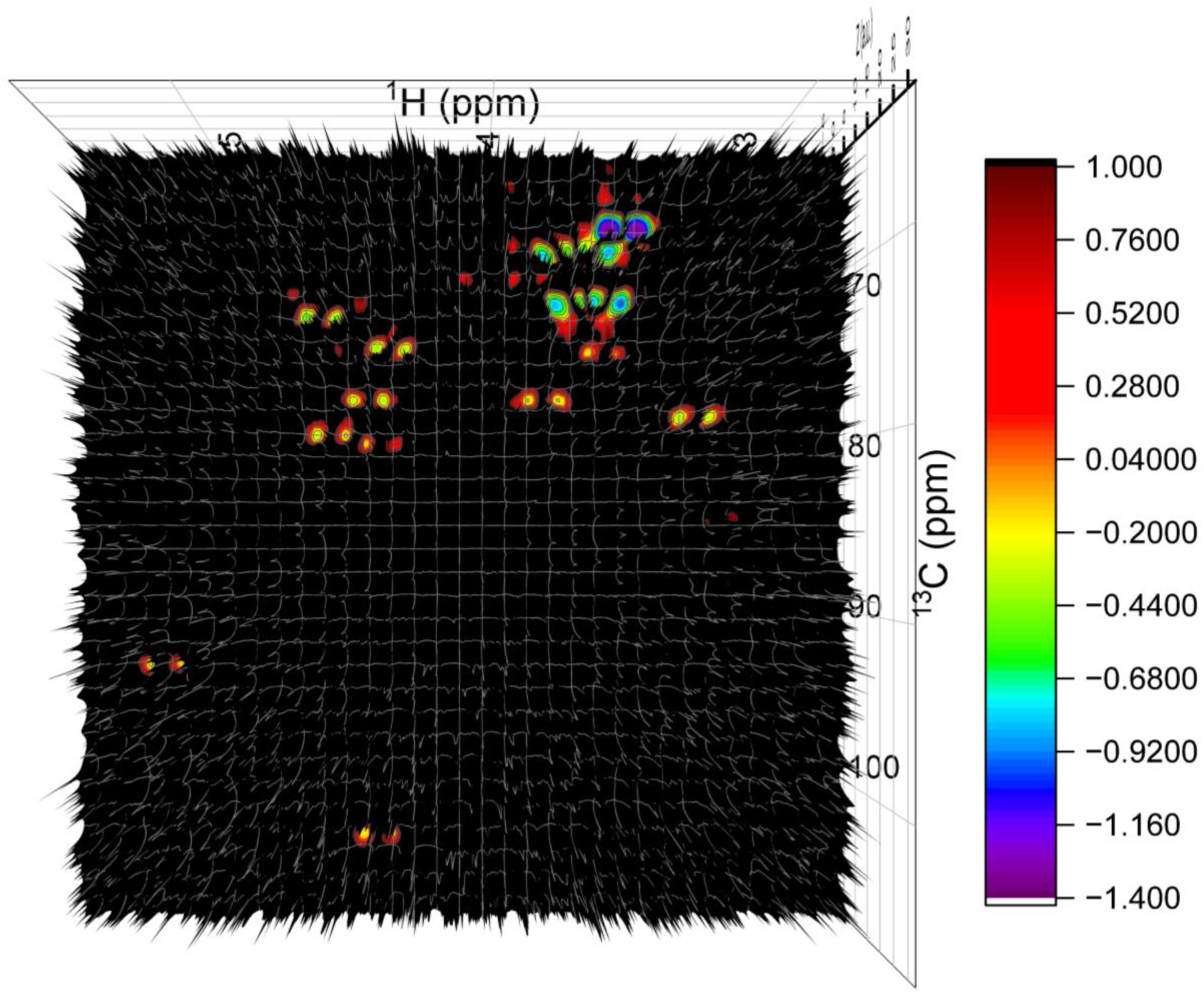
Top-view dark forest image (3D colormap surface plot) of 2D ^1^H–^13^C HSQC data of per-*O*-ethylated iota-carrageenan after FDP–LCT along the ^1^H dimension, with the surface generated from 𝑍 values corresponding to ^1^H and ^13^C chemical shifts and the color scale displayed on the right.

**Fig. 5.**
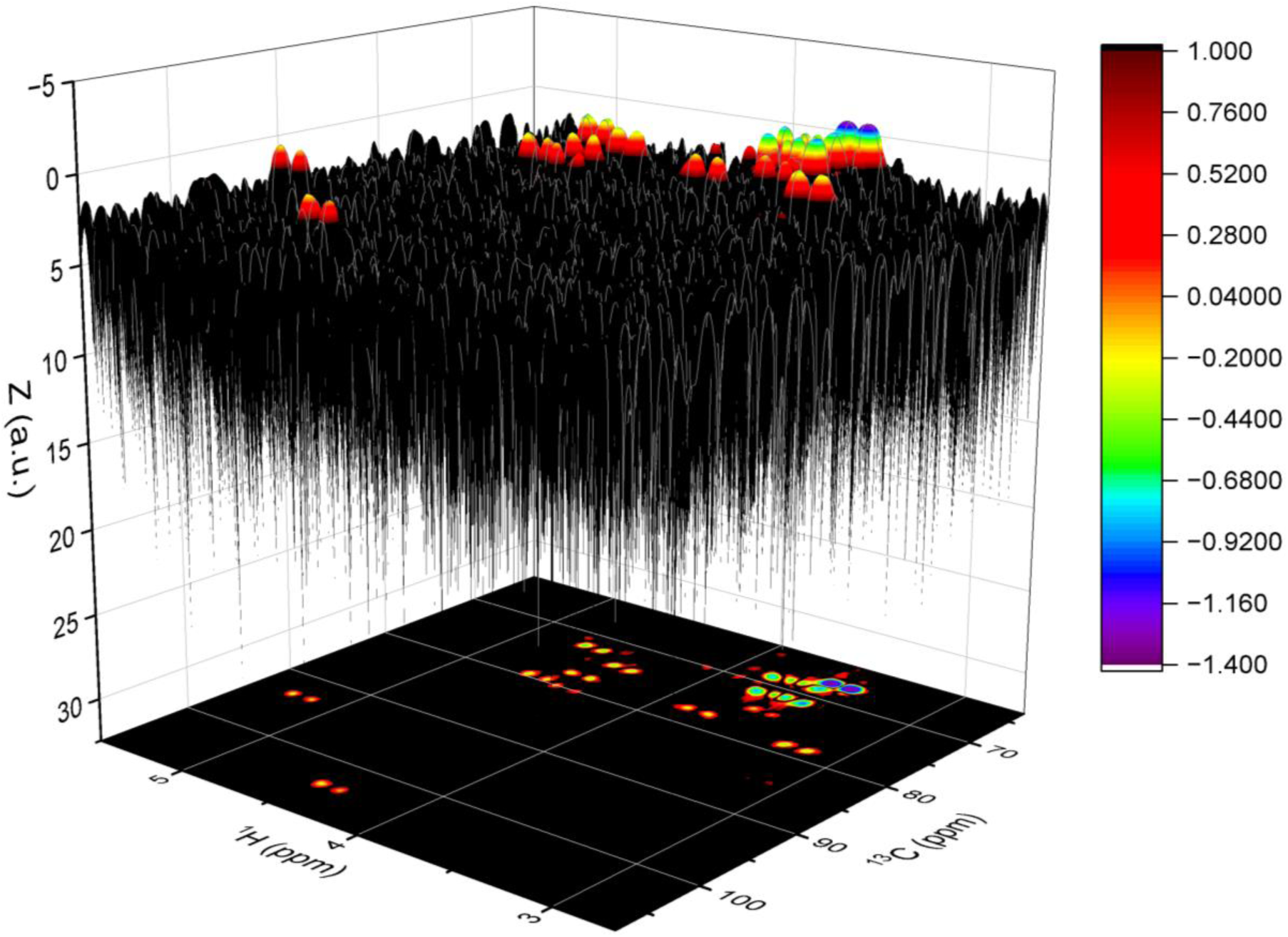
Inverted side-view dark forest image (3D colormap surface plot) of 2D ^1^H–^13^C HSQC data of per-*O*-ethylated iota-carrageenan after FDP–LCT along the ^1^H dimension. The surface is generated from 𝑍 values corresponding to ^1^H and ^13^C chemical shifts, with the color scale displayed on the right and the corresponding 2D contour map projection shown below.

**Fig. 6.**
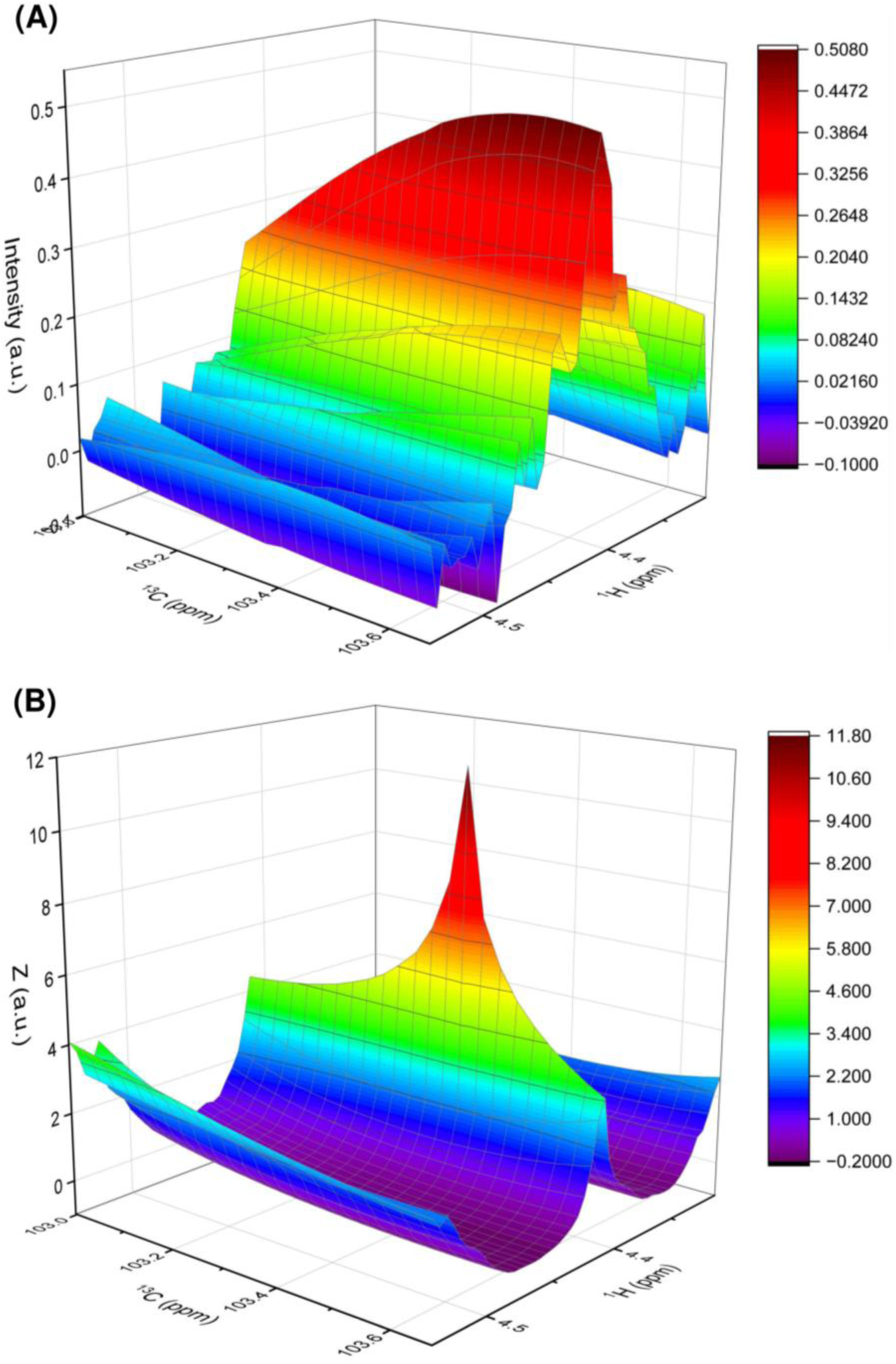
3D colormap surface plots of HSQC data in the region corresponding to the correlation between C-1 and H-1 of residue D in per-*O*-ethylated iota-carrageenan, showing (A) the data before and (B) the data after FDP–LCT along the ^1^H dimension. The surfaces are generated from the original intensities in (A) and from 𝑍 values in (B), corresponding to ^1^H and ^13^C chemical shifts, with the color scale displayed on the right.

**Fig. 7.**
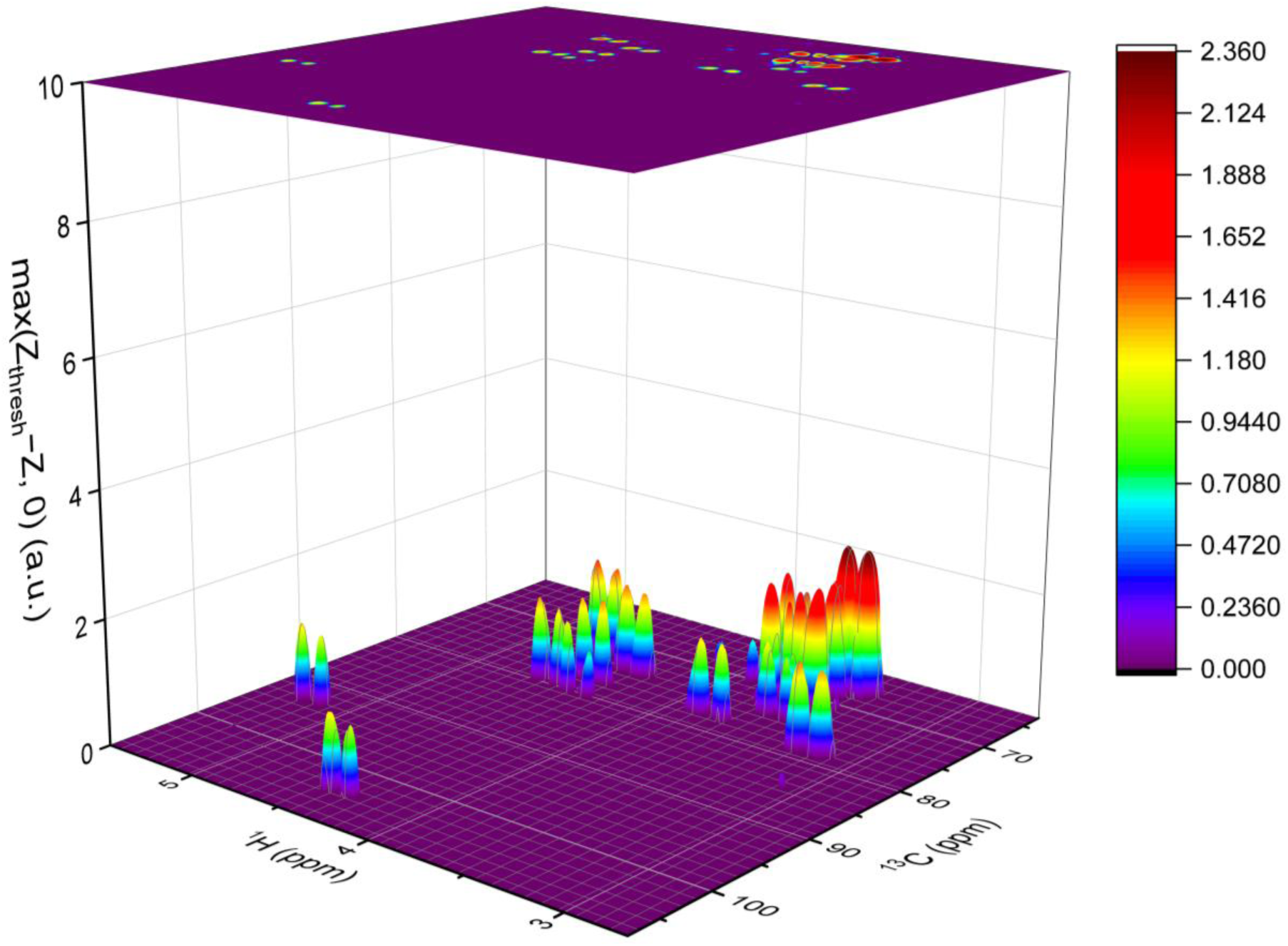
3D colormap surface plot of 2D ^1^H–^13^C HSQC data of per-*O*-ethylated iota-carrageenan after FDP–LCT along the ^1^H dimension, with the surface generated from max(Z_thresh_ − 𝑍, 0) corresponding to ^1^H and ^13^C chemical shifts, the color scale displayed on the right, and the corresponding 2D contour map projection shown above. Z_thresh_ is an empirically optimized threshold value, and a Z_thresh_ of 1.0 was used for plotting this figure.

**Fig. 8.**
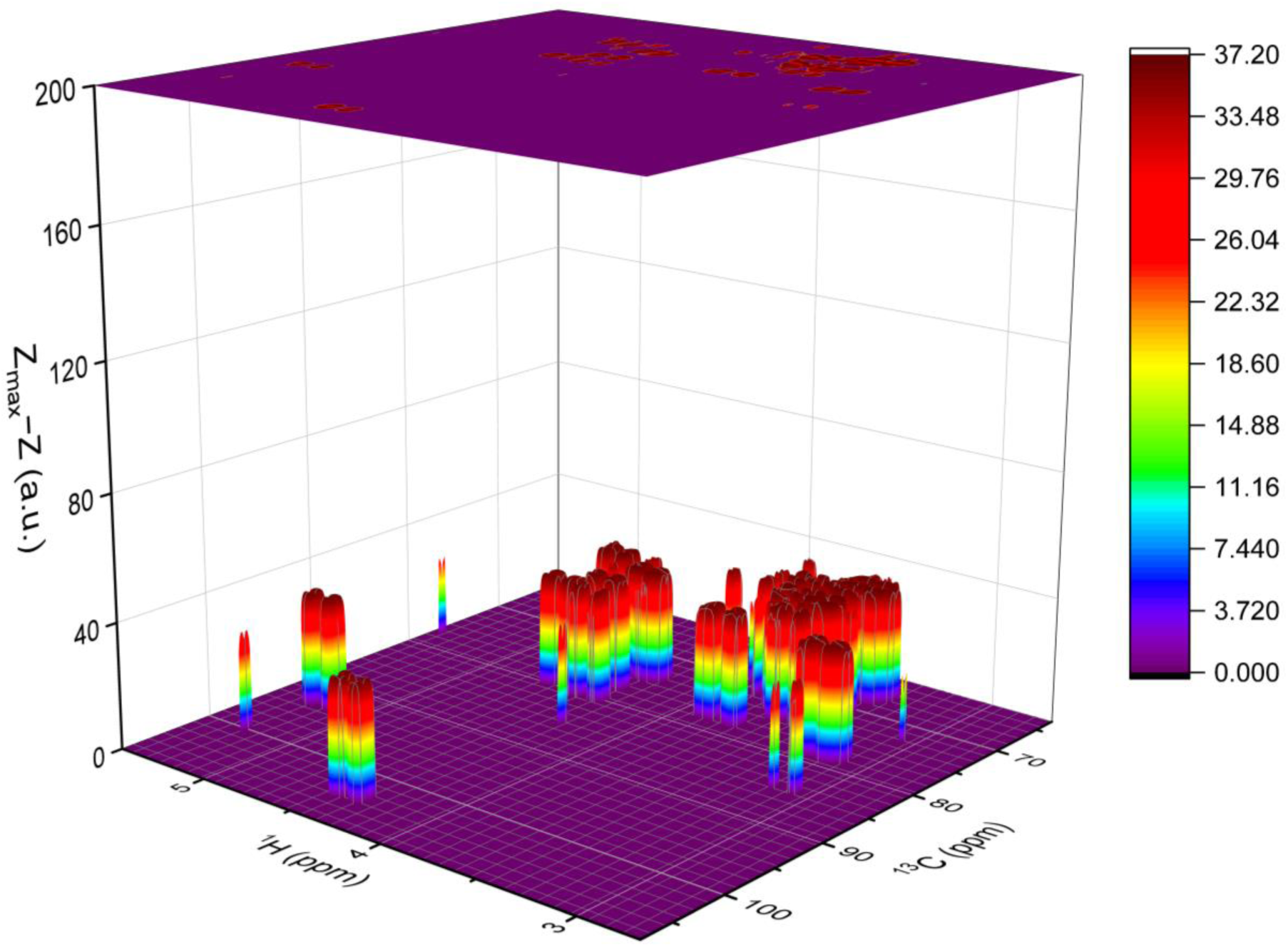
3D colormap surface plot of 2D ^1^H–^13^C HSQC data of per-*O*-ethylated iota-carrageenan after FDP–LCT applied along the ^13^C dimension to the processed data shown in Fig. 7. The surface is generated from Z_max_ − 𝑍, corresponding to ^1^H and ^13^C chemical shifts, with the color scale displayed on the right and the corresponding 2D contour map projection shown above. Z_max_is a constant set as 36.7368005696771, as detailed in Section 2 (Theory).

**Fig. 9.**
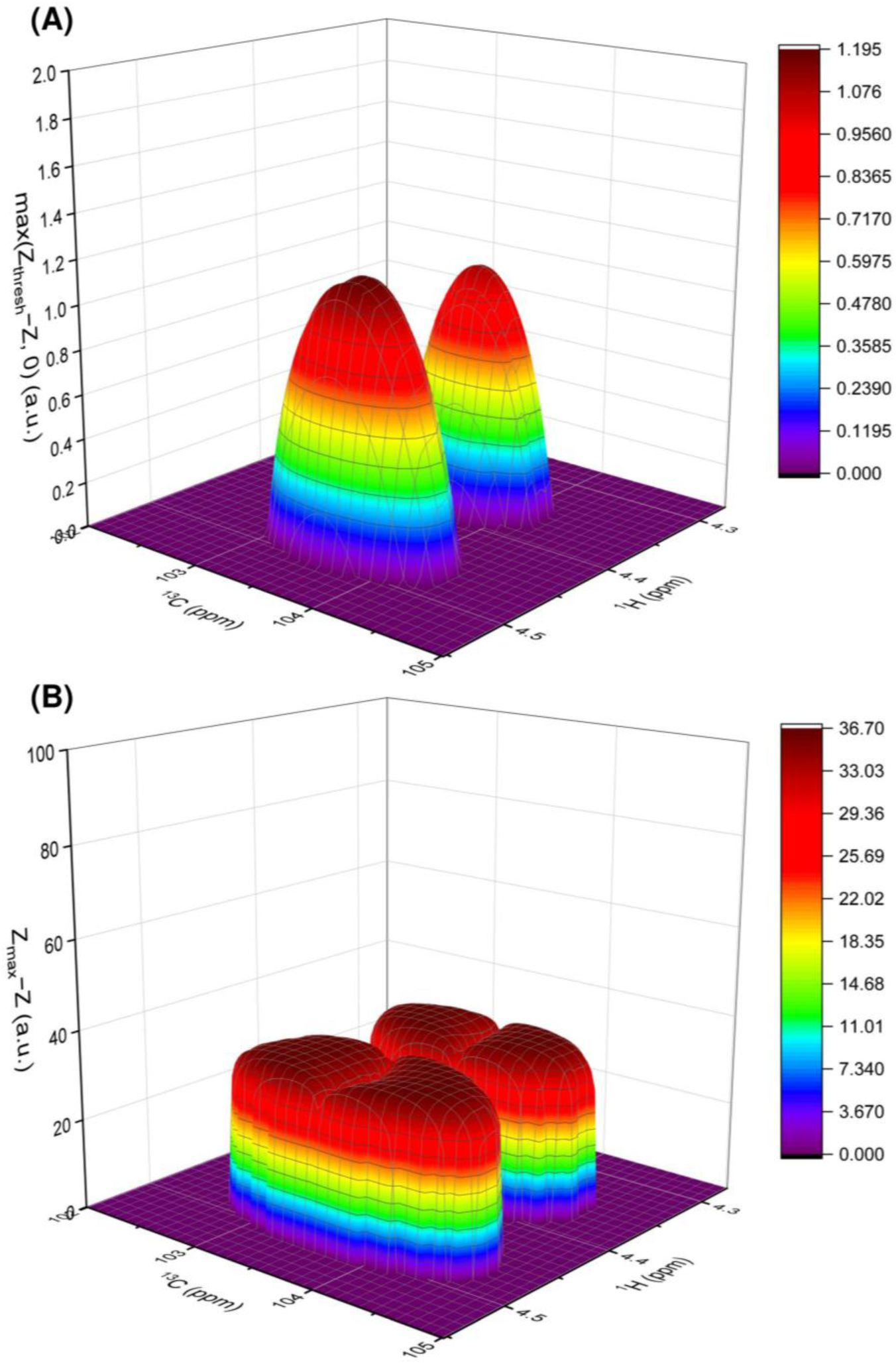
Regions of 3D colormap surface plots of processed ^1^H–^13^C HSQC data, zoomed in from (A) Fig. 7 and (B) Fig. 8, respectively, corresponding to the correlation between C-1 and H-1 of residue D in per-*O*-ethylated iota-carrageenan.

**Fig. 10.**
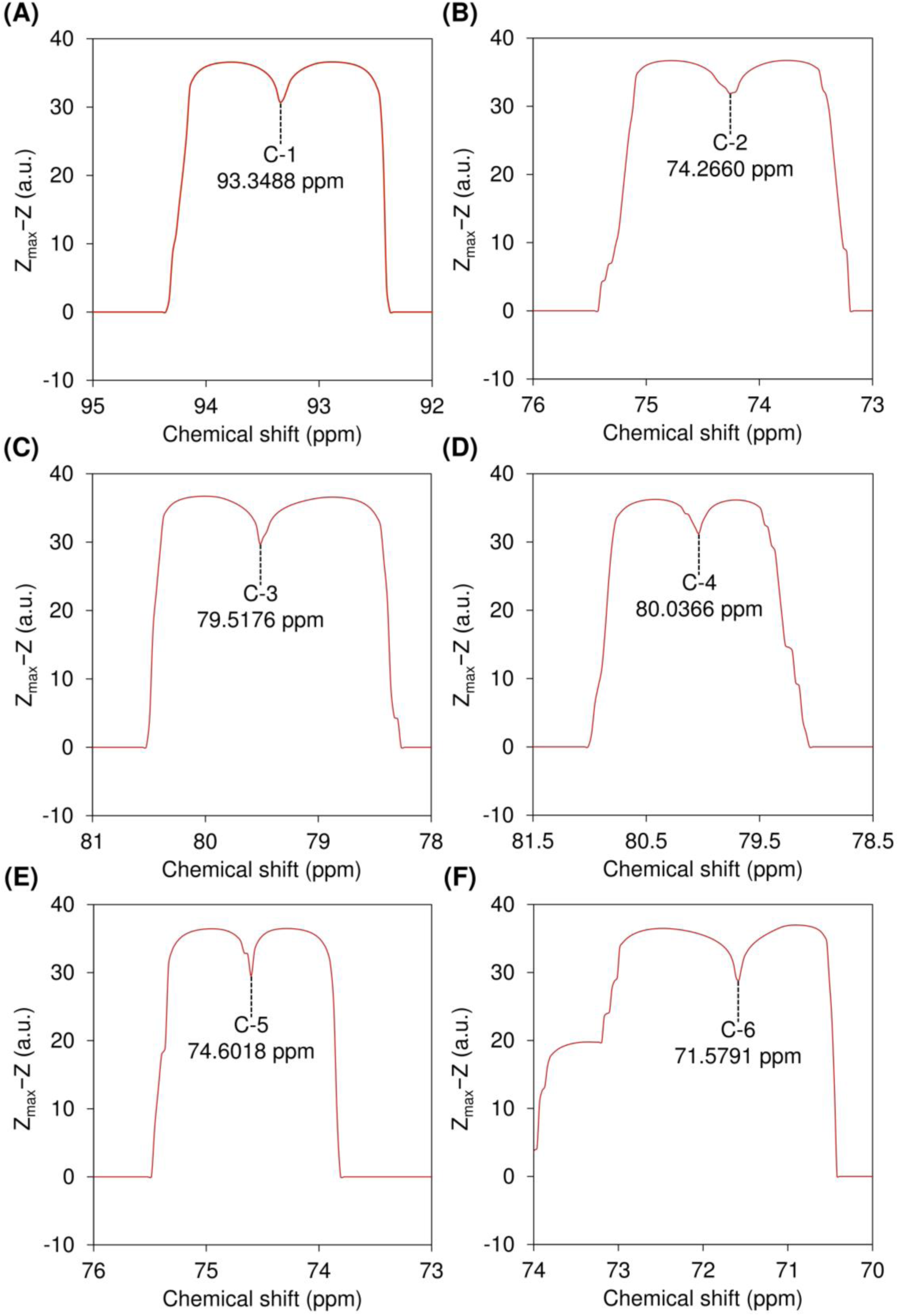
Local resolution-boosted spectra of sugar ring carbons from 4-linked 2-*O*-sulfo-3,6-anhydro-α-D-galactopyranose (residue C), extracted from 2D ^1^H–^13^C HSQC data of per-*O*-ethylated iota-carrageenan after FDP–LCT along the ^1^H dimension followed by the ^13^C dimension. Z_max_is a constant set as 36.7368005696771, as detailed in Section 2 (Theory).

**Fig. 11.**
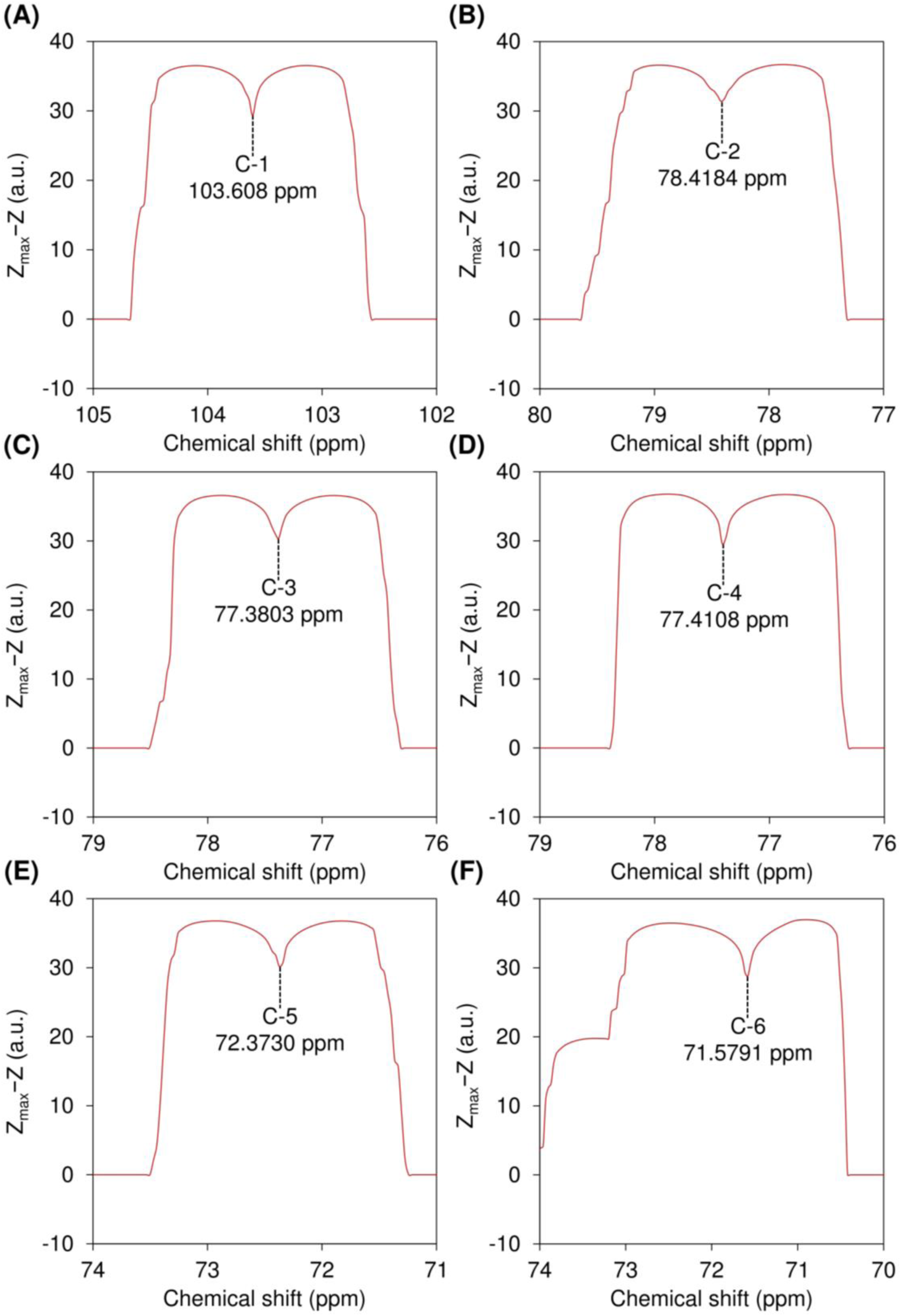
Local resolution-boosted spectra of sugar ring carbons from 3-linked 2,6-di-*O*-ethyl-4-*O*-sulfo-β-D-galactopyranose (residue D), extracted from 2D ^1^H–^13^C HSQC data of per-*O*-ethylated iota-carrageenan after FDP–LCT along the ^1^H dimension followed by the ^13^C dimension. Z_max_ is a constant set as 36.7368005696771, as detailed in Section 2 (Theory).

**Fig. 12.**
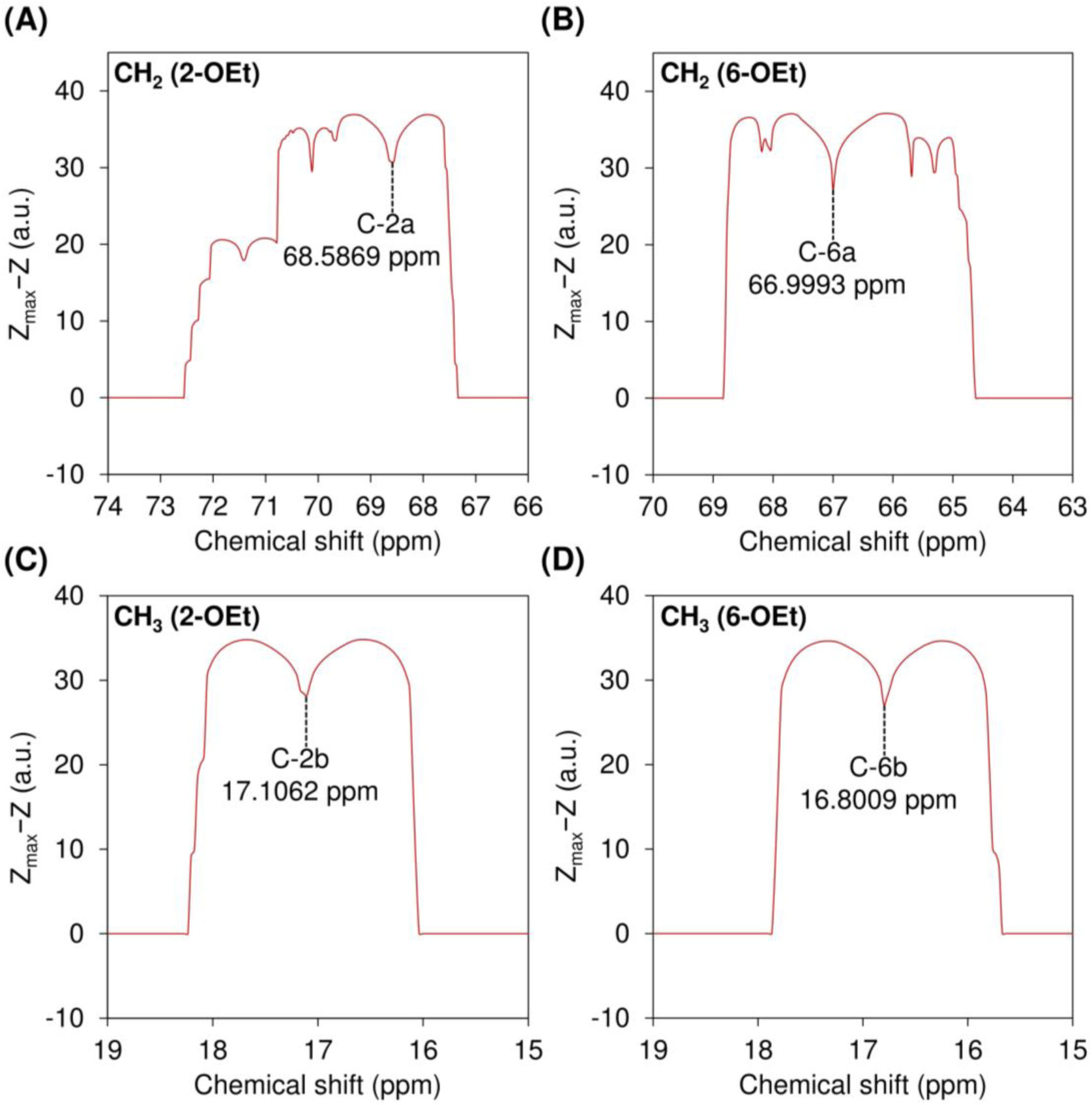
Local resolution-boosted spectra of carbons on the methylene (CH2) and methyl (CH3) groups of the ethyl substituents at the *O*-2 and *O*-6 positions (designated 2-OEt and 6-OEt, respectively) of 3-linked 2,6-di-*O*-ethyl-4-*O*-sulfo-β-D-galactopyranose (residue D), extracted from 2D ^1^H–^13^C HSQC data of per-*O*-ethylated iota-carrageenan after FDP–LCT along the ^1^H dimension followed by the ^13^C dimension. Z_max_ is a constant set as 36.7368005696771, as detailed in Section 2 (Theory).

**Fig. 13.**
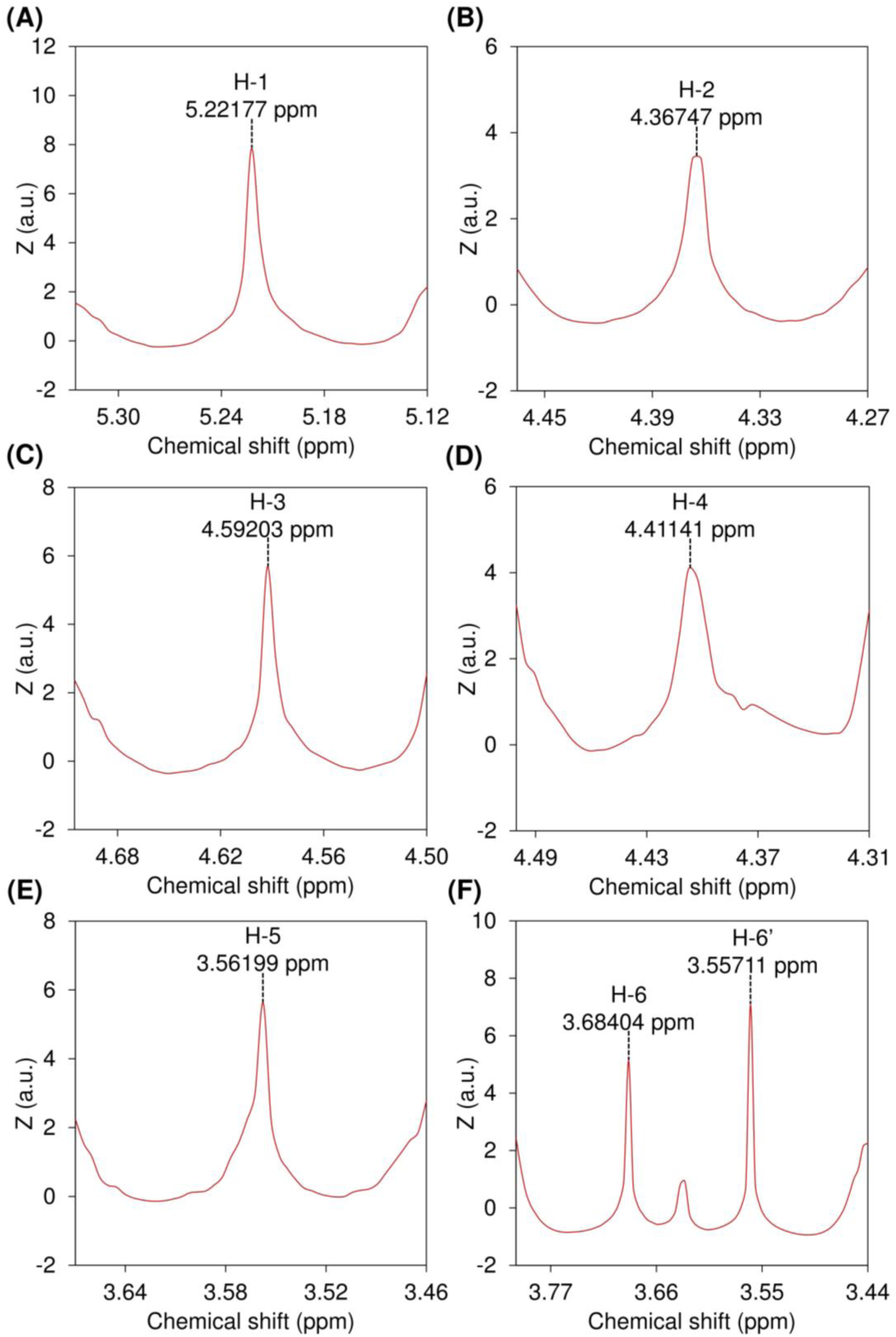
Local resolution-boosted spectra of sugar ring protons from 4-linked 2-*O*-sulfo-3,6-anhydro-α-D-galactopyranose (residue C), extracted from 2D ^1^H–^13^C HSQC data of per-*O*-ethylated iota-carrageenan after FDP–LCT along the ^1^H dimension.

**Fig. 14.**
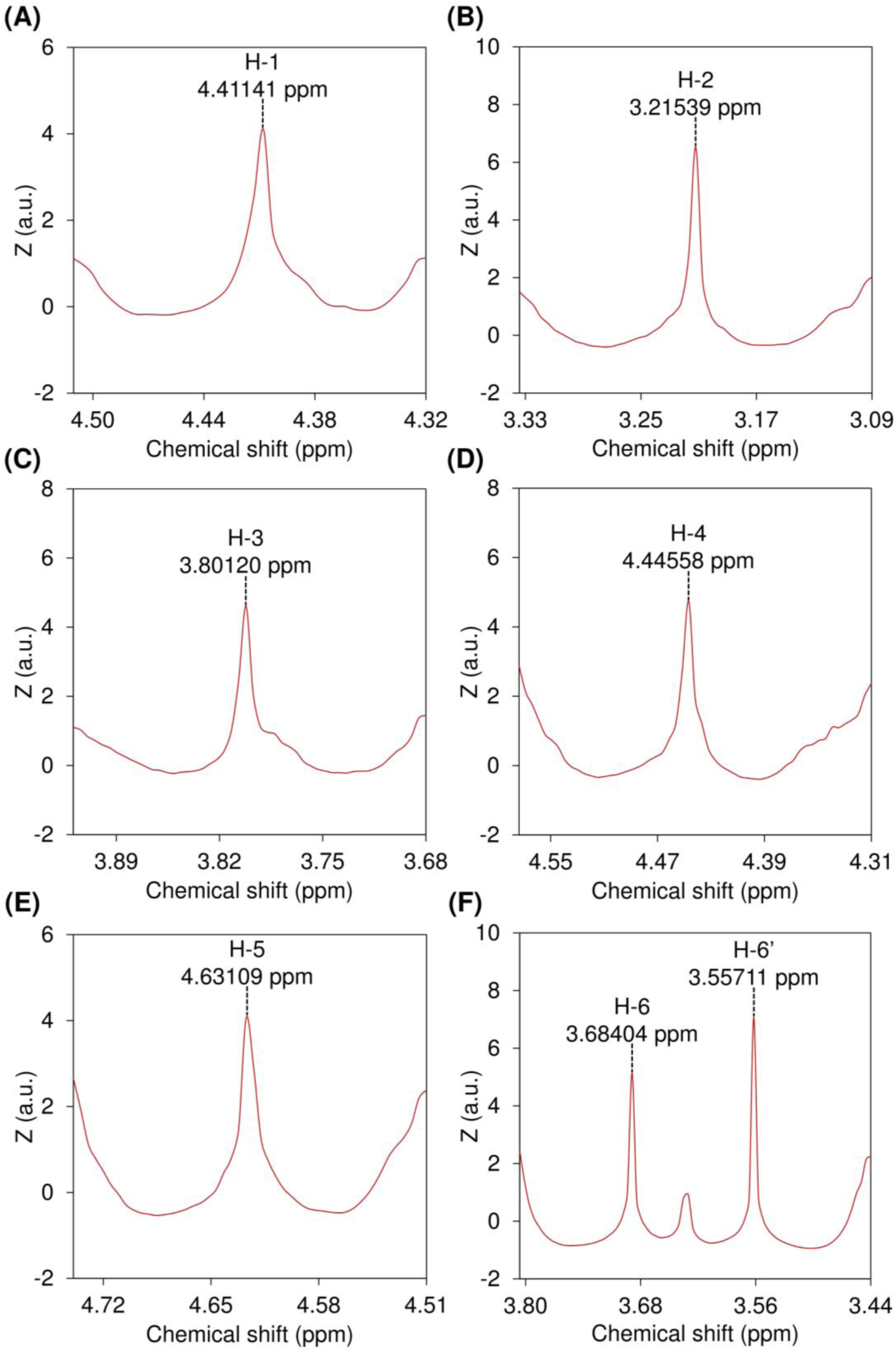
Local resolution-boosted spectra of sugar ring protons from 3-linked 2,6-di-*O*-ethyl-4-*O*-sulfo-β-D-galactopyranose (residue D), extracted from 2D ^1^H–^13^C HSQC data of per-*O*-ethylated iota-carrageenan after FDP–LCT along the ^1^H dimension.

**Fig. 15.**
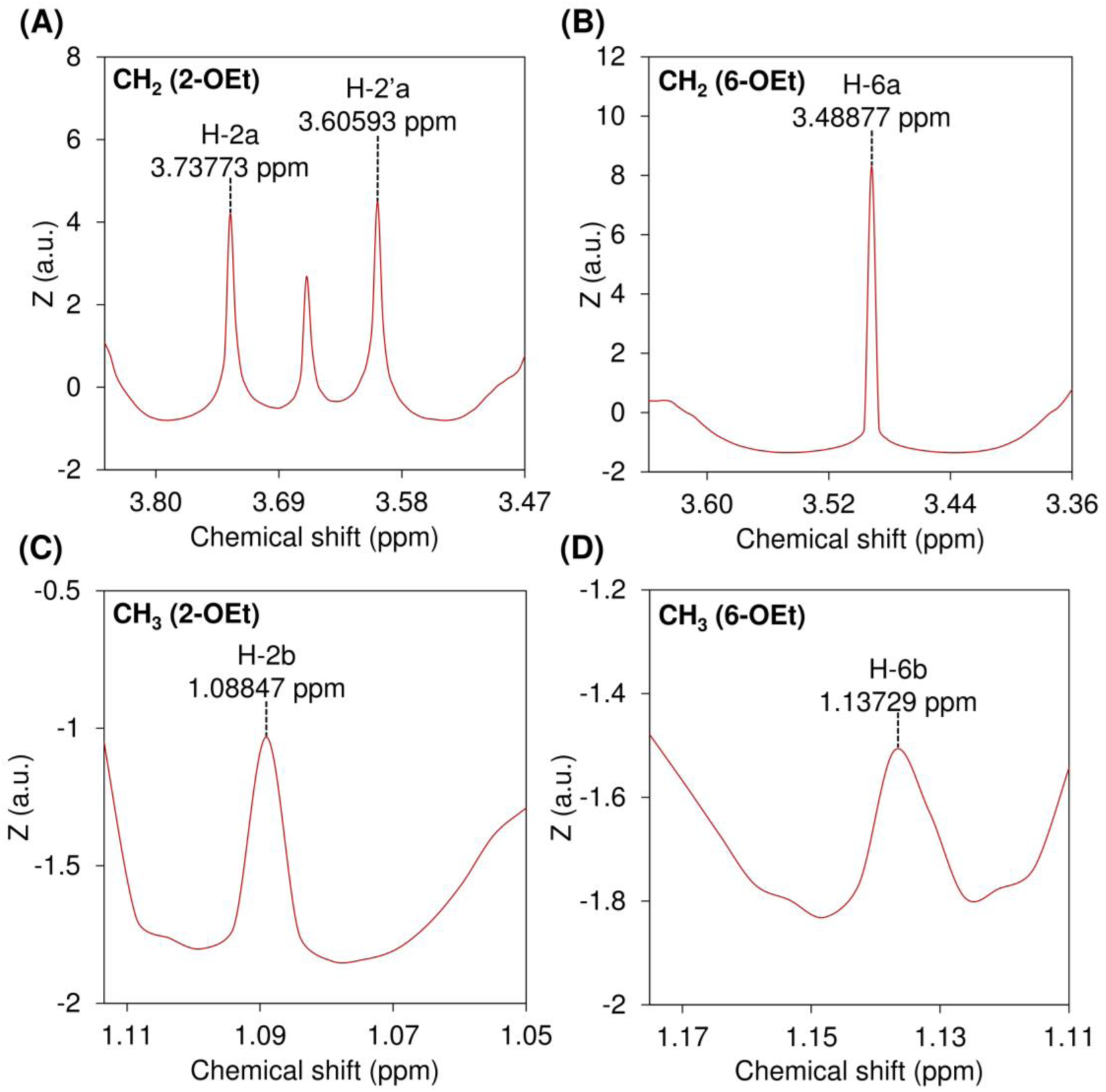
Local resolution-boosted spectra of protons on the methylene (CH2) and methyl (CH3) groups of the ethyl substituents at the *O*-2 and *O*-6 positions (designated 2-OEt and 6-OEt, respectively) of 3-linked 2,6-di-*O*-ethyl-4-*O*-sulfo-β-D-galactopyranose (residue D), extracted from 2D ^1^H–^13^C HSQC data of per-*O*-ethylated iota-carrageenan after FDP–LCT along the ^1^H dimension.

After the first step of processing, in which each 1D slice along the ^1^H dimension was subjected to FDP–LCT, the 2D HSQC data of per-*O*-ethylated iota-carrageenan yielded a 2D matrix densely populated with spikes (Fig. 3). These spikes arise from zero or near-zero derivative values corresponding to local extrema in the 2D HSQC spectra and represent resolution-boosted features that collectively create the characteristic “dark forest” appearance, with spikes mimicking “trees”. Importantly, the spikes originate not only from genuine one-bond ^1^H–^13^C cross-peaks of iota-carrageenan but also from local extrema of baseline undulations and small artifact peaks, meaning that both true signals and background fluctuations contribute to the dense landscape observed in the 3D surface contour plot (Fig. 3). This demonstrates that LCT conversion enhances local resolution while simultaneously reducing global clarity. However, in the top view (Fig. 4), the landscape does not appear as a uniformly dark forest; instead, the central spike from each prominent cross-peak is flanked by two symmetric valleys, highlighted in warm colors and standing in sharp contrast to the dense distribution of black spikes, which becomes even more pronounced in the inverted side-view of the dark forest image (Fig. 5). The color contrast reflects contour-level settings: regions above an empirically optimized 𝑍 threshold (Z_thresh_, set to 1 for the per-*O*-ethylated iota-carrageenan examples in Figs. 3–5) were rendered in black, whereas regions below this threshold were displayed in a gradient from warm to cold colors, with the corresponding scales shown in the gradient bars alongside the surface plots. The depth and broadness of these concave valleys arise from the steepness and large spatial extent of strong cross-peaks. By contrast, weak background fluctuations yield only shallow and narrow features: their small derivatives keep 𝑍 values relatively high, and their limited peak size restricts the spatial extent, preventing the development of broad, spike-free regions, as further supported by the absence of small warm-colored spots within the black regions surrounding the broad valleys in the 2D projection for the contour surface (Figs. 3 and 5). Examples of the locally resolution-enhanced features are also shown in Fig. 6, which presents a zoom-in view of the region showing the C-1/H-1 correlation of residue D before and after FDP–LCT along the ^1^H dimension.

In the second step of processing, the conversion operation max(Z_thresh_ − 𝑍, 0) transforms 𝑍 values smaller than the empirically optimized threshold Z_thresh_ into positive values Z_thresh_ − 𝑍 > 0 while converting 𝑍 values greater than or equal to Z_thresh_into zero, thereby generating a new surface plot in which positive regions correspond to the smooth valleys (concave regions) that were spike-free in the original dark forest image from Step 1, whereas spike-dense regions are reduced to zero (Fig. 7). This transformation offers two advantages. First, it converts the negative values of the smooth valleys into positive values (turning concave into convex), thereby preserving these features in a more interpretable form. Second, it eliminates the spike signals by reducing them to zero, simplifying the image. It deserves mentioning that the removal of spikes effectively eliminates features arising from noise, while for genuine cross-peaks, the spikes are suppressed but the corresponding smooth valleys are retained and converted to positive values, ensuring that the essential cross-peak features remain represented.

For the third step of processing, FDP–LCT was conducted along the ^13^C dimension for each slice to enhance the local resolution of ^13^C signals. In the fourth step, instead of directly plotting the new 𝑍 values generated in Step 3, these values were transformed according to the operation Z_max_ − 𝑍 and then plotted to generate Fig. 8. According to Equation 5, background values equal to zero prior to transformation would otherwise become Z_max_after transformation, but subtracting from Z_max_ ensures that the converted background remains zero. This operation also turns all newly generated nonzero 𝑍 values corresponding to the broad, spike-free concave regions in the dark forest image into positive values, which appear as major peaks characterized at their centers by an inverted wedge-shaped feature perpendicular to the ^13^C axis, as exemplified in the zoom-in image (Fig. 9). It should be noted that the wedge feature originates from the local resolution enhancement of ^13^C signals generated by the FDP–LCT along the ^13^C dimension in Step 3, whereas the reversed orientation of the wedge arises directly from subtracting the newly generated 𝑍 values in Step 3 from Z_max_in Step 4.

The fifth step of processing allows zooming into the region of each major peak and averaging the longest columns of continuous nonzero values near the peak center, thereby minimizing artifacts from potential peak overlap at the contour edges. The resulting 1D ^13^C projections effectively captured the peak characteristics along the ^13^C dimension, represented by a broad peak containing a reversed sharp inverse spike produced by local resolution enhancement near the peak midpoint (Figs. 10–12, S13, S14, and S23). The corresponding ^13^C chemical shift was then determined from this sharp inverse spike (Tables 1 and 2). It is important to note that each original cross-peak yields a pair of symmetric spike-free surfaces, from which the 1D ^13^C projection is obtained by averaging the central columns of nonzero values across both surfaces. However, in cases where one of the two surfaces is overlapped by signals from other peaks, the less overlapped surface can be used to obtain the 1D projection and thereby determine the ^13^C chemical shift.

**Table 1.** Chemical shifts of ^1^H and ^13^C nuclei in the sugar rings of per-*O*-ethylated carrageenans.

| Per-*O*-ethylated carrageenan | Residues | <sup>1</sup>H and <sup>13</sup>C chemical shifts (ppm) |  |  |  |  |  |  |  |
| --- | --- | --- | --- | --- | --- | --- | --- | --- | --- |
|  |  |  | 1 | 2 | 3 | 4 | 5 | 6 | 6' |
| Kappa | Residue **A** | δ<sub>H</sub> | 5.07 | 3.86 | 4.19 | 4.31 | 3.61 | 3.71 | 3.56 |
|  |  | δ<sub>C</sub> | 94.8 | 77.5 | 80.0 | 80.0 | 74.8 | 72.0 | – |
|  | Residue **B** | δ<sub>H</sub> | 4.38 | 3.11 | 3.74 | 4.40 | 4.54 | 3.71 | 3.56 |
|  |  | δ<sub>C</sub> | 103.6 | 78.9 | 78.0 | 77.2 | 71.7 | 72.0 | – |
| Iota | Residue **C** | δ<sub>H</sub> | 5.22 | 4.37 | 4.59 | 4.41 | 3.56 | 3.68 | 3.56 |
|  |  | δ<sub>C</sub> | 93.3 | 74.3 | 79.5 | 80.0 | 74.6 | 71.6 | – |
|  | Residue **D** | δ<sub>H</sub> | 4.41 | 3.22 | 3.80 | 4.45 | 4.63 | 3.68 | 3.56 |
|  |  | δ<sub>C</sub> | 103.6 | 78.4 | 77.4 | 77.4 | 72.4 | 71.6 | – |

Note: “–” means “not applicable”. Residues A and B represent 4-linked 2-*O*-ethyl-3,6-anhydro-α-D-galactopyranose and 3-linked 2,6-di-*O*- ethyl-4-*O*-sulfo-β-D-galactopyranose in per-*O*-ethylated kappa-carrageenan, respectively. Residues C and D represent 4-linked 2-*O*-sulfo-3,6- anhydro-α-D-galactopyranose and 3-linked 2,6-di-*O*-ethyl-4-*O*-sulfo-β-D-galactopyranose in per-*O*-ethylated iota-carrageenan, respectively.

**Table 2.** Chemical shifts of ^1^H and ^13^C nuclei in the methylene groups and methyl groups of the ethyl groups of per-*O*-ethylated carrageenans.

| Per-*O*-ethylated carrageenans | Residues | Ethylation position |  | <sup>1</sup>H and <sup>13</sup>C chemical shifts (ppm) |  |
| --- | --- | --- | --- | --- | --- |
|  |  |  |  | Methylene | Methyl |
| Kappa | Residue **A** | *O*-2 | δ<sub>H</sub> | 3.58, 3.87 | 1.10, 1.10, 1.10 |
|  |  |  | δ<sub>C</sub> | 67.8 | 17.1 |
|  | Residue **B** | *O*-2 | δ<sub>H</sub> | 3.63, 3.76 | 1.12, 1.12, 1.12 |
|  |  |  | δ<sub>C</sub> | 68.6 | 17.1 |
|  |  | *O*-6 | δ<sub>H</sub> | 3.49, 3.49 | 1.12, 1.12, 1.12 |
|  |  |  | δ<sub>C</sub> | 67.0 | 16.9 |
| Iota | Residue **C** | — | δ<sub>H</sub> | — | — |
|  |  |  | δ<sub>C</sub> | — | — |
|  | Residue **D** | *O*-2 | δ<sub>H</sub> | 3.61, 3.74 | 1.09, 1.09, 1.09 |
|  |  |  | δ<sub>C</sub> | 68.6 | 17.1 |
|  |  | *O*-6 | δ<sub>H</sub> | 3.49, 3.49 | 1.14, 1.14, 1.14 |
|  |  |  | δ<sub>C</sub> | 67.0 | 16.8 |

Note: “—” means “not applicable”. Residues A and B represent 4-linked 2-*O*-ethyl-3,6-anhydro-α-D-galactopyranose and 3-linked 2,6-di-*O*- ethyl-4-*O*-sulfo-β-D-galactopyranose in per-*O*-ethylated kappa-carrageenan, respectively. Residues C and D represent 4-linked 2-*O*-sulfo-3,6- anhydro-α-D-galactopyranose and 3-linked 2,6-di-*O*-ethyl-4-*O*-sulfo-β-D-galactopyranose in per-*O*-ethylated iota-carrageenan, respectively.

Step 6 closes the loop of the workflow by returning to the dark forest image generated in Step 1, using the ^13^C chemical shift obtained in Step 5 as a locator to guide the extraction of the corresponding 1D ^1^H slice where the cross-peak apex emerges as a sharp central spike flanked by broad concave curves (Figs.13–15, S15, S16, S24, and S25), a feature that stands out clearly from the background spikes and enables accurate determination of the ^1^H chemical shift (Tables 1 and 2). The extracted chemical shifts, without rounding, are summarized in Tables S1 and S2.

The F60 fraction was isolated from *Mazzaella japonica*, a red seaweed in Gigartinaceae family known to synthesize both iota- and kappa-carrageenans [57, 60–62]. The F60 sample was collected as the final fraction soluble in 60% ethanol (w/w), which represented the highest ethanol concentration used during gradient ethanol precipitation for the purification of sulfated galactans, during which ethanol was gradually added dropwise to the vigorously stirred hot-water extract until reaching a final concentration of 60% (w/w), followed by centrifugation, dialysis, and lyophilization of the resulting supernatant to yield F60. Although F60 remained soluble in 60% ethanol (w/w) prior to lyophilization, its water solubility markedly decreased afterward, with visible insoluble material observed upon attempted dissolution in deionized water, even after heating at 70 °C with magnetic stirring overnight. The poor water solubility was further confirmed by the absence of prominent high-molecular-weight polysaccharide signals in a preliminary high-performance size-exclusion chromatography (HPSEC) analysis (data not shown). Consequently, it was not feasible to prepare a D2O solution of F60 for solution-state NMR analysis; however, the per-*O*-ethylated F60 derivative dissolved readily in DMSO-d6, allowing acquisition of high-quality solution-state NMR spectra. The results of the 2D NMR, monosaccharide, and linkage analyses of per-*O*-ethylated F60 are presented in the Supplementary Figure and Tables (Figs. S26–S37; Tables S3 and S4). The ^1^H and ^13^C chemical shifts (Tables 1 and 2) extracted from the 2D NMR spectra of the per-*O*-ethylated commercial iota- and kappa-carrageenan standards were used to identify cross-peaks corresponding to iota- and kappa-carrageenan components in F60. These assignments were validated by consistent correlations across the HSQC, COSY, TOCSY, and HMBC spectra of per-*O*-ethylated F60 (Figs. S26–S31). In addition to these characteristic signals from the iota- and kappa-carrageenans, some minor cross-peaks were observed whose chemical shifts did not match those of the iota-or kappa-type disaccharide units, indicating the presence of other carrageenan components in F60, as further supported by the results of GC–MS-based linkage analysis (Figs. S34–S37; Table S4). The results of monosaccharides and linkage analyses of F60 (Figs. S26–S37; Tables S3 and S4) were in good agreement with methylation and desulfation–methylation analyses of the same F60 sample described in our recent report [62].

## 5. Discussion

### 5.1. FDP–LCT workflow for post-processing 2D HSQC spectra

First-derivative analysis of spectroscopic data has long been employed in carbohydrate research, particularly in vibrational spectroscopy for quantitative and qualitative purposes [64–67], but has been only rarely used in carbohydrate NMR analysis [68] and, to our knowledge, has never been applied to the post-processing of 2D ^1^H–^13^C HSQC NMR data. Based on the novel FDP–LCT algorithm developed in this study, we present a new spectral analytical framework in which local extrema and inflection points in the original spectrum are transformed respectively into zero crossings and extrema in the first derivative spectrum, and subsequently into sharp spikes and flanking troughs under LCT processing, as detailed in Section 2 (Theory). When applied to the 2D ^1^H–^13^C HSQC spectra of per-*O*-ethylated carrageenans, FDP–LCT along the ^1^H dimension generated the characteristic dark forest image, in which cross-peak apices along each ^1^H slice were converted into spikes that enhanced local resolution, while overall clarity was reduced because background fluctuations also produced spikes. Importantly, only genuine cross-peaks gave rise to smooth valleys flanking the central spikes, which, through threshold screening and FDP–LCT along the ^13^C dimension, were converted into surfaces with a central inverse-wedge feature from which ^13^C chemical shifts were extracted. The loop was completed by returning to the dark-forest image, using the extracted ^13^C values as locators to guide the extraction of ^1^H slices carrying resolution-enhanced signals characterized by sharp central spikes flanked by broad concave valleys, thereby enabling accurate determination of ^1^H chemical shifts of the target cross-peaks.

The procedure involved two steps of FDP–LCT, with the first applied along the ^1^H dimension to enhance the resolution of the ^1^H slices, and the second along the ^13^C dimension to enhance that of the ^13^C slices. Notably, direct application of FDP–LCT along the ^13^C dimension to the 2D HSQC data, rather than to the threshold-filtered output of the ^1^H-processed data, was preliminarily tested and yielded poor results (data not shown). This may be because the relatively broad peaks along the ^13^C dimension generate valleys of insufficient depth, resulting in spectra with only weak, shallow, and indistinct features that make it difficult to define contour levels capable of distinguishing spike-free valley features arising from cross-peaks against the highly spiked background. By contrast, peaks along the ^1^H dimension are narrower and sharper, producing deeper and smoother valleys that provide more effective inputs for subsequent ^13^C-directed FDP–LCT, thereby enabling reliable extraction of ^13^C chemical shifts.

This post-processing method shows clear potential for practical application and implementation. By capturing both peak-apex information as resolution-boosted spikes and peak-steepness information as smooth valleys, the FDP–LCT algorithm showed its ability to extract and process features of HSQC cross-peaks and revealed potential as a feature-extraction tool for 2D NMR. This is central importance for any 2D NMR analysis where the indirect dimension is broadened due to either poor digital resolution or sample properties such as viscosity. FDP-LCT can also be applied in vibrational spectroscopy data of carbohydrates and beyond, and shows potential in situations where the extracted spectral features could be directly applied in chemometric analyses such as machine learning-based classification and pattern recognition [69]. Another potential lies in automation, as the workflow (Steps 1–6) implemented in this study utilized four basic data-processing operations: smoothing, derivative transformation, LCT conversion, and spike picking. These can be readily executed in programming environments, thereby allowing for full automation of the pipeline for spectroscopic data processing in a rapid high-throughout manner.

### 5.2. Advantages and potential of per-O-ethylation–2D NMR analysis of carrageenans

This study demonstrates the feasibility of per-*O*-ethylation for preparing samples suitable for solution-state ^1^H and ^13^C NMR investigation of carrageenan structures. Bearing negatively charged sulfate groups, carrageenans readily form complexes with cations and proteins in aqueous systems, thereby limiting water solubility [29–31]. Consequently, complete dissolution of native carrageenans in D2O for solution-state NMR generally requires highly purified samples largely free of proteins and interfering cations, but is not achievable for crude, unfractionated extracts such as the *Mazzaella japonica* F60 fraction. Once dissolved in D2O, the high charge density of carrageenans generates strong electrostatic repulsion, which expands the chain conformation and produces a viscous solution that restricts molecular motion [70], thereby accelerating transverse relaxation (shorter T2) and resulting in broad NMR signals [16, 17]. Sample solution viscosity can be reduced by lowering concentration, but this simultaneously weakens signal intensity and limits sensitivity, especially in ^13^C experiments, which typically require more scans and thus longer acquisition time to compensate [16, 17]. Per-*O*-ethylation circumvents these obstacles: the dialysis steps in derivatization provide additional purification to remove ions [57]; more importantly, the derivatives dissolve readily in DMSO-d6 due to favorable alkyl–solvent interactions [10, 15, 38], which prevent complexation with ions or proteins and allow for solution-state NMR spectral acquisition.

Per-*O*-ethylated kappa- and iota-carrageenans exhibit distinct 2D ^1^H–^13^C HSQC and other 2D NMR spectra, as the difference in the *O*-2 substituent of the 3,6-anhydrogalactose residue (*O*-sulfate in iota and *O*-ethyl in kappa) alters local electronic environments sufficiently to distinguish both the anhydrogalactose units and their neighboring galactose residues. Using the complete ^1^H and ^13^C chemical shifts obtained from per-*O*-ethylated commercial kappa- and iota-carrageenan standards, the 2D NMR spectra of the per-*O*-ethylated F60 fraction isolated from *Mazzaella japonica* were assigned, revealing kappa- and iota-specific signals together with minor cross-peaks indicative of other carrageenan components. These results highlight the need to interpret 2D NMR spectra of per-*O*-ethylated reference carrageenans beyond the kappa and iota types to establish a broader chemical shift library for reliable structural assignment in complex mixtures. For example, a recent per-*O*-methylation-based 2D HSQC study of higher-plant cell wall polysaccharides used ten per-*O*-methylated polysaccharide standards to build a chemical shift database that enabled identification of not only major wall polysaccharides but also low-abundance components (*e.g.*, glucomannan) whose signals were otherwise difficult to assign by NMR correlation data alone [10]. It is worth noting that in previous studies, both deuterated chloroform (CDCl3) [10] and deuterated dichloromethane (CD2Cl2) [71] were used to dissolve per-*O*-methylated polysaccharides, as these low-viscosity solvents are well suited for 2D NMR acquisition at room temperature. However, chloroform and dichloromethane are not well suited for dissolving per-*O*-alkylated sulfated polysaccharides, which dissolve better in DMSO [42, 57], where heating effectively reduces solution viscosity and enables the acquisition of high-quality NMR spectra. We also observed that per-*O*-ethylated carrageenans dissolved more readily in DMSO than their per-*O*-methylated counterparts [62], likely due to the greater DMSO affinity of ethyl groups, although their solubility was not quantitatively determined. Furthermore, per-*O*-ethylation offers two major analytical advantages for qualitative and quantitative 2D NMR analysis. For qualitative purposes, the ethyl substituent provides additional structural information, as the methylene and methyl groups within the ethyl moiety generate strong cross-signals that, while partially overlapping for substitutions at different positions in HSQC spectra, are well resolved in COSY, TOCSY, and HMBC spectra. It also enables direct confirmation of substitution sites through long-range correlations between the methylene protons and carbons of the ethyl group and the adjacent *O*-linked carbons and protons of the sugar ring, making both the location and completeness of ethylation self-explanatory from 2D NMR data. For quantitative purposes, per-*O*-ethylation enhances the detection sensitivity of free hydroxyl groups, those not involved in glycosidic linkages, natural sulfation, or other substitutions, by labeling each site with an ethyl group containing two carbons and five protons, thereby increasing ^13^C and ^1^H abundance and thus signal intensity, which is particularly advantageous for ^13^C nuclei with low natural abundance [16, 17].

Finally, it is worth noting that even in the nominally pure iota-carrageenan sample, the 2D NMR analysis was sensitive enough to detect very weak yet discernible signals attributable to kappa-type disaccharide structures. Their consistent occurrence across multiple 2D spectra and the excellent agreement of their chemical shifts with those extracted from per-*O*-ethylated kappa-carrageenan strongly suggest that these represent genuine structural features rather than random noise. These minor kappa-type signals were easily detectable in the homonuclear COSY (Fig. S5) and TOCSY (Fig. S6) spectra. They also remained in the heteronuclear HSQC spectrum, although very weak and requiring a low contour threshold to visualize, and were observable in the FDP*–*LCT-processed HSQC spectra (Figs. 3*–*5, 7, and 8). Nevertheless, it remains uncertain whether these weak signals originate from trace levels of kappa-type disaccharide units within the iota-carrageenan chain (structural heterogeneity) or from trace levels of separate kappa-carrageenan chains (sample impurity).

## 6. Conclusions

This study established a new method for post-processing 2D ^1^H–^13^C HSQC NMR spectra by novel FDP–LCT approach. The algorithm converted 2D HSQC spectra into dark forest images with enhanced local resolution, enabling the extraction of sharpened spectral features and reliable ^1^H and ^13^C chemical shifts, as demonstrated with per-*O*-ethylated kappa- and iota-carrageenans. Importantly, the FDP–LCT workflow provides an effective post-acquisition means of improving digital resolution in the indirect dimension without increasing experimental time, a benefit broadly applicable to 2D NMR experiments that require high resolution in both dimensions. It was also shown that dissolution of these per-*O*-ethylated carrageenans in DMSO-d6 enabled the collection of high-quality HSQC and other 2D spectra, permitting complete signal assignment and allowing clear distinction between two sulfated galactans whose structures differ by only a single *O*-2 substitution. Together, these advances broadened the analytical toolbox for resolving closely related complex polysaccharide structures by solution-state 2D ^1^H–^13^C NMR.

## CRediT authorship contribution statement

Conceptualization, X.X. and D.W.A.; methodology, X.X., T.M., and D.W.A.; software, X.X.; validation, X.X., T.M., S.W.C., and D.W.A.; formal analysis, X.X.; investigation, X.X., J.P.T., B.B., and V.W.; resources, D.W.A. and T.M.; data curation, X.X.; writing – original draft preparation, X.X.; writing – review and editing, X.X., T.M., S.W.C., and D.W.A.; visualization, X.X.; supervision, D.W.A.; project administration, D.W.A.; funding acquisition, D.W.A. All authors have read and agreed to the published version of the manuscript.

## Declaration of competing interest

The authors declare that they have no known competing financial interests or personal relationships that could have appeared to influence the work reported in this paper.

## Data availability

Data will be made available on request.

## Acknowledgments

We thank senior research technician Ms. Yolanda Brummer of the Guelph Research and Development Centre, Agriculture and Agri-Food Canada, for her technical support and for assisting with HPSEC data collection and analysis of *Mazzaella japonica* carrageenan. We also thank Dr. Spencer Serin and Mr. Edgar Smith (Beaver Meadow Farms Ltd.) for their assistance in sourcing *Mazzaella japonica* from Deep Bay in the Salish Sea.

## Appendix A. Supplementary data

Supplementary data include “Supplementary Materials and Methods” and “Supplementary Figures and Tables”.

## Notes

### Competing Interest Statement

The authors have declared no competing interest.

