## Supplementary Materials and Methods for "2D HSQC-derived “dark forest” image with enhanced local resolution via first derivative processing–logarithmic cosine transformation (FDP–LCT): Demonstration on per-*O*-ethylated kappa- and iota-carrageenans"

### Supplementary Material and Methods

#### *Chemicals, reagents, and consumables*

Dimethyl sulfoxide (Cat. No. 276855), iodoethane (Cat. No. I7780), sodium hydroxide (Cat. No. S5881), methanol (Cat. No. 322415), trifluoroacetic acid (Cat. No. 8.08260), glacial acetic acid (Cat. No. 69509), acetic anhydride (Cat. No. 242845), triethylamine hydrochloride (Cat. No. 8.21135), 4-methylmorpholine borane (Cat. No. 262323), dichloromethane (Cat. No. 1.00668), sodium bicarbonate (Cat. No. S8875), anhydrous sodium sulfate (Cat. No. 8.22286), hexane (Cat. No. 139386), ethyl acetate (Cat. No. 1.10972), and Wilmad 5 mm precision NMR tubes (Cat. No. Z412007) were purchased from Sigma-Aldrich Co., LLC (Burlington, MA, USA). Spectrum Spectra/Por 1 regenerated cellulose dialysis tubing with molecular weight cut-off (MWCO) of 6,000–8,000 Da (Cat. No. 08-670D), sodium borodeuteride with an isotopic purity of 99% (Cat. No. 035102.06), and 1-Cyclohexyl-3-(2-morpholinoethyl) carbodiimide metho-p-toluenesulfonate (Cat. No. AC111360050) were purchased from Thermo Fisher Scientific Inc. (Waltham, MA, USA). Ethanol (Cat. No., P016EAAN) was purchased from Commercial Alcohols Inc. (Chatham, ON, Canada). Thermostable α-amylase from *Bacillus licheniformis* (Cat. No. E-BLAAM) was purchased from Megazyme Ltd. (Bray, County Wicklow, Ireland).

Deionized water was generated from a Barnstead D0809 Nanopure II system (Thermo Fisher Scientific Inc., MA, USA). All chemical reactions were conducted in Pyrex® disposable glass tubes (thread 15-415, Cat. No. CLS99447161) sealed with Corning® phenolic caps with PTFE liners (Cat. No. CLS999815) and stirred using BRAND® magnetic stirring bars coated with PTFE (8 mm in length and 3 mm in diameter, Cat. No. Z328936), all purchased from Sigma-Aldrich Co., LLC (MA, USA). $N_2$ from a cylinder was used to fill the headspace of the tube, and a Stuart evaporator (Cole-Parmer, Vernon Hills, IL, USA) supplied with $N_2$ from a generator was used for the evaporation of samples to dryness.

*Mazzaella japonica* was collected along the beachfront near Maple Guard Drive, British Columbia (49.441695° N, 124.676979° W), as described in our recent report [1]. The samples were immediately flash-frozen and stored in liquid nitrogen, then freeze-dried and ball-milled.

***Extraction of crude polysaccharide from Mazzaella japonica***

Ball-milled dry powder of *Mazzaella japonica* (137.6 g) was soaked in 1.2 L of hexane under magnetic stirring for 2 h in a 2 L glass beaker covered with aluminum foil to reduce evaporation. The mixture was left undisturbed overnight to allow the powder to settle. The supernatant was carefully decanted using a glass pipette, leaving approximately 1 cm of liquid above the interface to avoid disturbing the precipitate. An additional 1.2 L of hexane was added, and the extraction process was repeated. The precipitate was then resuspended in 1.2 L of 95% (v/v) ethanol, magnetically stirred for 8 h, and centrifuged at 3,000 × *g* for 30 min at room temperature. The resulting pellet was resuspended in 1.2 L of 80% (v/v) ethanol, with the same stirring and centrifugation conditions applied, and the extraction process was repeated. The resulting pellet was evaporated to dryness in 50 mL centrifuge tubes using SpeedVac (Thermo Fisher Scientific, MA, USA). The dried sample was then transferred to a glass beaker and underwent two rounds of water extraction at room temperature with constant magnetic stirring, followed by three rounds of hot water extraction at 70 °C in an incubator. Each extraction used 2 L of deionized water and lasted 8 h, with the beaker covered with aluminum foil. After each extraction, centrifugation (3,000 × *g*, 30 min, room temperature) was conducted, and the supernatant was collected while the pellet was carried forward to the next extraction. Supernatants from all the water extractions were pooled, poured into 40 L of absolute ethanol, and left at room temperature overnight, followed by centrifugation (3,000 × *g*, 30 min, room temperature). The precipitate was evaporated to dryness in 50 mL centrifuge tubes using the SpeedVac, redissolved in 2 L of deionized water by incubating at 70 °C overnight, and freeze-dried (56.7 g).

***Amylase treatment of crude polysaccharide extracted from Mazzaella japonica***

Dry crude polysaccharide (14.4 g) was dissolved in 1 L of deionized water by incubating at 70 °C overnight. After that, 2 mL of thermostable α-amylase (3,000 units/mL, Megazyme, Ireland) was added, and the mixture was incubated at 70 °C for 8 h [2]. The solution was then poured into 4 L of absolute ethanol, left standing at 4 °C overnight, and centrifuged (3,000 × *g*, 30 min, room temperature). The residue was evaporated to dryness using the SpeedVac, redissolved in 500 mL of deionized water by incubating at 70 °C overnight, extensively dialyzed with MWCO of 6,000–8,000 Da against deionized water at 4 °C, and freeze-dried (13.8 g).

***Purification of Mazzaella japonica sulfated galactan by gradient ethanol precipitation***

This gradient ethanol precipitation procedure was adapted from a previously reported method [3]. Amylase-treated polysaccharide (915 mg) was dissolved in 400 mL of deionized water by incubating at 70 °C overnight. The resulting solution was cooled to room temperature, left standing at 4 °C overnight, and then centrifuged (3,000 × *g*, 30 min, room temperature). The supernatant was vigorously stirred magnetically to create a water tunnel in a 2 L glass beaker. Absolute ethanol was slowly added dropwise to achieve a 15% (w/w) ethanol concentration. The mixture was kept at 4 °C for 8 h, followed by centrifugation (3,000 × *g*, 30 min, room temperature). Absolute ethanol was then added dropwise to the vigorously stirred supernatant until the ethanol concentration reached 30% (w/w), followed by standing at 4 °C and centrifugation as described above. The resulting supernatant underwent two additional cycles of ethanol precipitation, with ethanol concentrations gradually increased to 45% and 60% (w/w), respectively. The final supernatant was evaporated to dryness in 50 mL centrifuge tubes using the SpeedVac. The dry sample was redissolved in 100 mL of deionized water by incubating at 70 °C overnight, and the resulting solution was freeze-dried (0.9 g, designated as fraction F60).

***1D and 2D NMR analysis of per-O-ethylated derivatives of commercial iota-carrageenan, commercial kappa-carrageenan, and Mazzaella japonica F60 fraction***

Detailed parameters for each 1D and 2D NMR experiment performed on the per-*O*-ethylated derivatives of the commercial iota- and kappa-carrageenan standards, as well as the F60 fraction isolated from *Mazzaella japonica*, are described as follows:

1D $^{1}$H NMR spectra of the per-*O*-ethylated derivatives of iota-carrageenan, kappa-carrageenan, and F60 were recorded at 700.44 MHz using the standard zg30 pulse sequence with a 1 s relaxation delay, and automatic phase and baseline corrections applied to all datasets. For per-*O*-ethylated F60, spectra were acquired with a 3.1195 s acquisition time, a 15 ppm (10,504.2 Hz) spectral width, and 3,500 scans, yielding an acquired size of 32,768 points, which was zero filled to a final spectral size of 131,072 points. The per-*O*-ethylated iota-carrageenan sample was measured using 512 scans over a 3.1999 s acquisition period, with a spectral width of 20 ppm (14,097.7 Hz) and an acquired size of 45,112 points, also zero filled to 131,072 points. Similarly, the per-*O*-ethylated kappa-carrageenan sample was analyzed with 3,400 scans under the same acquisition time and spectral width conditions as the per-*O*-ethylated iota-carrageenan, with an acquired size of 45,112 points zero filled to 131,072 points.

1D $^{13}$C NMR spectra of the per-*O*-ethylated iota-carrageenan, kappa-carrageenan, and F60 samples were recorded using the zgpg30 pulse sequence with a 2 s relaxation delay, 1.5729 s acquisition time, and automatic phase and baseline corrections applied. For per-*O*-ethylated F60, spectra were acquired with 15,000 scans at a spectrometer frequency of 176.14 MHz, with a spectral width of 236 ppm (41,666.7 Hz); data were collected as 65,536 points and subsequently zero filled to a final spectral size of 2,097,152 points. The per-*O*-ethylated iota-carrageenan sample was measured under the same acquisition conditions as the per-*O*-ethylated F60 sample, except that 2,000 scans were collected; the acquired data were zero filled to a final size of 131,072 points. The per-*O*-ethylated kappa-carrageenan sample was likewise recorded under the same acquisition parameters, using 12,000 scans, and the acquired data were zero filled to 131,072 points.

2D $^{1}$H–$^{13}$C HSQC spectra were acquired using the hsqcedetgpsp.3 pulse sequence on a 700.44 MHz spectrometer (with a 176.13 MHz channel) for the per-*O*-ethylated iota-carrageenan, kappa-carrageenan, and F60 samples. All experiments employed automatic phase and baseline correction and applied a sine-square apodization at 90° along both $F_1$ and $F_2$ (the first $F_1$ point fixed at 0.50 and no adjustment in $F_2$). For per-*O*-ethylated F60, 16 scans were acquired with a 2.0000 s relaxation delay and a 0.4503 s acquisition time, using spectral widths of 9.0 ppm (6,329.1 Hz, $F_2$) and 113.5 ppm (20,000.0 Hz, $F_1$). Data were acquired as 2,850 ($t_2$) and 4,096 ($t_1$) points and processed to 4,096 ($F_2$) and 8,192 ($F_1$) points, with $F_2$ zero filled from 2,944 to 4,096 points (no linear prediction). For per-*O*-ethylated iota-carrageenan, 64 scans were recorded with a 1.5 s relaxation delay and a 0.2925 s acquisition time, using spectral widths of 10.0 ppm (7,002.8 Hz, $F_2$) and 125 ppm (22,026.4 Hz, $F_1$). Data were acquired as 2,048 ($t_2$) and 256 ($t_1$) points and processed to 2,048 ($F_2$) and 4,096 ($F_1$) points; $F_1$ was zero filled from 256 to 4,096 (without linear prediction) while $F_2$ remained unchanged. Denoise by volume of interest (VOI) compression [4], a function built into Mnova, was applied with a threshold of 7.00 and a minimum VOI of 40.was applied with a threshold of 7.00 and a minimum VOI of 40. For per-*O*-ethylated kappa-carrageenan, under identical acquisition parameters (64 scans, 1.5 s delay, 0.2925 s acquisition), data were acquired as 2,048 ($t_2$) and 256 ($t_1$) points and processed to 2,048 ($F_2$) and 1,024 ($F_1$) points, with $F_1$ zero filled from 256 to 1,024 (without linear prediction); VOI compression was applied with a threshold of 9.00 and a minimum VOI of 25. However, it is important to note that VOI was omitted for the 2D HSQC data of per-*O*-ethylated carrageenan standards used in FDP–LCT post-processing, as less-compressed data are preferable for accurate derivative analysis.

2D $^{1}$H–$^{1}$H COSY spectra were recorded using the cosygpmfqf pulse sequence on a 700.44 MHz spectrometer for the per-*O*-ethylated iota-carrageenan, kappa-carrageenan, and F60 samples. All experiments employed a 1.5 s relaxation delay and automatic baseline correction, and VOI denoising was omitted. For per-*O*-ethylated F60, 16 scans were acquired with a 1.2943 s acquisition time and spectral widths of 9.0 ppm (6,329.1 Hz, $F_2$) and 9.0 ppm (6,321.1 Hz, $F_1$). Data were acquired as 8129 points in $t_2$ and 800 in $t_1$, and processed to final sizes of 8,192 ($F_2$) and 2,048 ($F_1$) points. No apodization was applied in $F_2$ (first point fixed at 0.50), whereas a bell-shaped apodization was applied in $F_1$. The $F_1$ data were zero filled from 800 to 2,048 points without linear prediction. For per-*O*-ethylated iota-carrageenan, 64 scans were recorded with a 1.1698 s acquisition time and a spectral width of 10.0 ppm (7,002.8 Hz) in both dimensions. Data were acquired as 8192 ($t_2$) and 256 ($t_1$) points and processed to 8,192 ($F_2$) and 2,048 ($F_1$) points. A sine-square apodization at 90° was applied in both dimensions (with the first $F_2$ point fixed at 0.50). The $F_1$ data were zero filled from 256 to 1,024 points, extended by MIST linear prediction to 2,048 points, and COSY-like symmetrization was additionally applied. For per-*O*-ethylated kappa-carrageenan, 84 scans were acquired under identical acquisition parameters as iota, yielding 8,192 ($t_2$) and 256 ($t_1$) points processed to 8,192 ($F_2$) and 2,048 ($F_1$). A sine-square apodization was applied in $F_2$ at 0° (first point fixed at 0.50) and in $F_1$ at 90°; $F_1$ was zero filled directly to 2,048 points without linear prediction, and no symmetry enforcement was performed.

2D $^{1}$H–$^{1}$H TOCSY spectra were recorded using the dipsi2gpphzs pulse sequence on a 700.44 MHz spectrometer for the per-*O*-ethylated iota-carrageenan, kappa-carrageenan, and F60 samples. All experiments employed automatic phase and baseline correction and omitted VOI denoising. A square apodization at 90° was applied along both $F_1$ and $F_2$ dimensions, with the first point fixed at 0.50 in $F_1$ and unadjusted in $F_2$. For per-*O*-ethylated F60, 16 scans were recorded with a 2.0000 s relaxation delay and a 1.2943 s acquisition time, using spectral widths of 9.0 ppm (6,329.1 Hz, $F_2$) and 9.0 ppm (6,321.1 Hz, $F_1$). Data were acquired as 8,192 ($t_2$) and 650 ($t_1$) points and processed to final sizes of 8,192 ($F_2$) and 2,048 ($F_1$), with no additional zero filling. For per-*O*-ethylated iota-carrageenan, 8 scans were recorded under identical relaxation and acquisition conditions, and the same spectral widths as per-*O*-ethylated F60. Data were acquired as 8,192 ($t_2$) and 650 ($t_1$) points and processed to 8,192 ($F_2$) and 2,048 ($F_1$) points; the $F_1$ data were further zero filled from 2,048 to 8,192 points (without linear prediction), and COSY-like symmetrization was additionally applied. For per-*O*-ethylated kappa-carrageenan, 58 scans were acquired with a 1.5000 s relaxation delay and a 0.2925 s acquisition time, employing spectral widths of 10.0 ppm (7,002.8 Hz, $F_1$ and $F_2$). Data were acquired as 2,048 ($t_2$) and 512 ($t_1$) points, processed to 2,048 ($F_2$) and 2,048 ($F_1$) points, with $F_1$ zero filled from 512 to 2,048 points (without linear prediction); no symmetrization was performed.

2D $^{1}$H–$^{13}$C HMBC spectra were recorded using the hmbcetgpl3nd pulse sequence on a 700.44 MHz spectrometer (176.13 MHz channel) for the per-*O*-ethylated iota-carrageenan, kappa-carrageenan, and F60 samples. Automatic phase and baseline corrections were applied in all cases, with no symmetry enforcement. For per-*O*-ethylated F60, 48 scans were acquired with a 2.0000 s relaxation delay and a 2.5887 s acquisition time, using spectral widths of 9.0 ppm (6,329.1 Hz, $F_2$) and 113.5 ppm (20,000.0 Hz, $F_1$). Data were acquired as 16,384 ($t_2$) and 1,024 ($t_1$) points and processed to 65,536 ($F_2$) and 2,048 ($F_1$) points; no apodization or zero filling was applied. For per-*O*-ethylated iota-carrageenan, 32 scans were recorded with a 1.5 s relaxation delay and a 2.5887 s acquisition time under identical spectral widths. Apodization was applied along $F_1$ using a sine-square function at 90° (first point fixed at 0.50) in combination with a sine-bell function at 45°, and $F_1$ data were zero filled from 1,024 to 4,096 points (without linear prediction), yielding final spectral sizes of 16,384 ($F_2$) and 4,096 ($F_1$). For per-*O*-ethylated kappa-carrageenan, 128 scans were acquired with a 1.5 s relaxation delay and a 0.2925 s acquisition time, using spectral widths of 10.0 ppm (7,002.8 Hz, $F_2$) and 125 ppm (22,026.4 Hz, $F_1$). Data were acquired as 2,048 ($t_2$) and 512 ($t_1$) points and processed to 2,048 ($F_2$) and 2,048 ($F_1$) points; $F_1$ was zero filled from 512 to 2,048 points (without linear prediction) after apodization identical to that used for per-*O*-ethylated iota-carrageenan. VOI compression was applied only for the kappa-carrageenan sample (threshold = 2.85; minimum VOI = 10) and omitted for per-*O*-ethylated F60 and iota-carrageenan.

***Monosaccharide composition analysis of F60 isolated from Mazzaella japonica***

Dry F60 (5 mg) was subjected to carbodiimide activation followed by sodium borodeuteride reduction to convert any uronic acids present in the sample to their 6,6'-dideuterated neutral sugars, following the published protocol [5]. An aliquot (1 mg) of the carboxyl-reduced sample was converted to C-1 deuterium-labeled alditol acetates (AAs) by 2 M trifluoracetic acid hydrolysis (120 °C, 2 h), sodium borodeuteride reduction, and acetylation by heating in a mixture of acetic anhydride and trifluoracetic acid (5:1, v/v) at 60 °C for 60 min [6, 7]. Another aliquot (1 mg) of the sample was subjected to reductive hydrolysis followed by acetylation to generate AAs including those from anhydrogalactose [8]. The derivatives were tested on an Agilent 7890A-

5977B GC-MS system (Agilent Technologies, CA, USA) equipped with a Supelco SP-2380 column (100 m × 0.25 mm × 0.2 µm; Sigma-Aldrich, MA, USA), with the oven temperature programmed to start at 100 °C (hold 1 min), followed by increases of 15 °C/min to 200 °C, then at 1 °C/min to 250 °C (hold 20 min). The derivatives were also tested on an Agilent 7890A GC-FID system (Agilent Technologies, CA, USA) equipped with a Supelco SP-2380 column (30 m × 0.25 mm × 0.2 µm; Sigma-Aldrich, MA, USA), with the oven temperature programmed to start at 60 °C (hold 1 min), followed by increases of 15 °C/min to 140 °C, 4 °C/min to 210 °C, then 8 °C/min to 250 °C (hold 15 min). An inlet temperature of 250 °C was used for both GC-MS and GC-FID analyses, with constant column helium flows of 0.8 mL/min for GC-FID and 1.2 mL/min for GC-MS. Derivatives were identified based on their EI-MS fragmentation patterns and by comparing retention times to those of standards. Anhydrogalactose and galactose were quantified using FID response factors obtained from iota-carrageenan standard (catalogue No. YC30038; Biosynth Carbosynth, USA ). Two separate experiments were conducted.

***Glycosidic linkage analysis of F60 fraction isolated from Mazzaella japonica***

Dry F60 (10 mg) was per-*O*-ethylated as described in Section 3.2, followed by redissolving the ethylation product in methanol, aliquoting, and then evaporating the aliquots to dryness under nitrogen [9]. One aliquot (2 mg) was subjected to 2 M trifluoracetic acid hydrolysis, sodium borodeuteride reduction, and acetylation to generate C-1 deuterium-labeled partially ethylated alditol acetates (PEAAs), while another aliquot (2 mg) underwent reductive hydrolysis and acetylation to generate PEAAs without deuterium labeling, including those from anhydrogalactose linkages, following the procedure detailed in our previous report [9]. The PEAAs were tested on an Agilent 7890A-5977B GC-MS system (Agilent Technologies, CA, USA) equipped with a Supelco SP-2380 column (60 m × 0.25 mm × 0.2 µm; Sigma-Aldrich, MA, USA), with oven temperature programmed to start at 120 °C (hold 1 min), followed by increases of 3 °C/min to 195 °C, 0.1 °C/min to 200 °C, and then 3 °C/min to 250 °C (hold 20 min). The same samples were also tested on an Agilent 7890A GC-FID system (Agilent Technologies, CA, USA), equipped with another identical Supelco SP-2380 column, using an oven program starting at 55 °C (hold 1 min), followed by increases of 20 °C/min to 120 °C, 3 °C/min to 195 °C, 0.1 °C/min to 200 °C, and then 3 °C/min to 250 °C (hold 20 min). Both GC-FID and GC-MS runs had identical settings for inlet temperature (250 °C) and constant column helium flow of 0.8 mL/min. Data acquisition and analysis were performed using Agilent OpenLab CDS software version 2.5 (Agilent Technologies, CA, USA). Identification of the PEAAs was based on their EI-MS ion fragmentation patterns, while quantitation was based on the FID response, using relative response factors calculated from the effective carbon number concept [10, 11], except for anhydrogalactose linkages. A relative response factor of 0.54 was used for the derivative from the 2,4-AnGal*p* linkage, according to our previous report [9], from which the response factor for the PEAA from the 4-AnGal*p* linkage was calculated to be 0.6, based on the principle that substitution of an ethyl group at the *O*-2 position, in place of an acetyl group, results in a 0.06 increase in the response factor according to the effective carbon number concept [10, 11]. Two separate experiments were conducted.
