## Supplementary Figures and Tables for "2D HSQC-derived “dark forest” image with enhanced local resolution via first derivative processing–logarithmic cosine transformation (FDP–LCT): Demonstration on per-*O*-ethylated kappa- and iota-carrageenans"

Table of contents


| Page No. | Figure/Table |
|---|---|
| 2 | **Fig. S1.** 2D COSY NMR spectrum of per-*O*-ethylated kappa-carrageenan |
| 3 | **Fig. S2.** 2D TOCSY NMR spectrum of per-*O*-ethylated kappa-carrageenan |
| 4 | **Fig. S3.** 2D HMBC NMR spectrum of per-*O*-ethylated kappa-carrageenan |
| 5, 6 | **Fig. S4.** 2D NMR region showing the ethyl group signals of per-*O*-ethylated kappa-carrageenan |
| 7 | **Fig. S5.** 2D COSY NMR spectrum of per-*O*-ethylated iota-carrageenan |
| 8 | **Fig. S6.** 2D TOCSY NMR spectrum of per-*O*-ethylated iota-carrageenan |
| 9, 10 | **Fig. S7.** 2D NMR region showing the ethyl group signals of per-*O*-ethylated iota-carrageenan |
| 11 | **Fig. S8.** Side-view dark forest image of HSQC data of per-*O*-ethylated kappa-carrageenan |
| 12 | **Fig. S9.** Top-view dark forest image of HSQC data of per-*O*-ethylated kappa-carrageenan |
| 13 | **Fig. S10.** Inverted side-view dark forest image of HSQC of per-*O*-ethylated kappa-carrageenan |
| 14, 15 | **Figs. S11, S12.** Further-processed HSQC data of per-*O*-ethylated kappa-carrageenan |
| 16 | **Fig. S13.** Local resolution-boosted spectra of sugar ring carbons from residue A |
| 17 | **Fig. S14.** Local resolution-boosted spectra of sugar ring carbons from residue B |
| 18 | **Fig. S15.** Local resolution-boosted spectra of sugar ring protons from residue A |
| 19 | **Fig. S16.** Local resolution-boosted spectra of sugar ring protons from residue B |
| 20 | **Fig. S17.** Dark forest image of HSQC (methyl region) of per-*O*-ethylated iota-carrageenan |
| 21, 22 | **Figs. S18, S19.** Further-processed HSQC (methyl region) of per-*O*-ethylated iota-carrageenan |
| 23 | **Fig. S20.** Dark forest image of HSQC (methyl region) of per-*O*-ethylated kappa-carrageenan |
| 24, 25 | **Figs. S21, S22.** Further-processed HSQC (methyl region) of per-*O*-ethylated kappa-carrageenan |
| 26 | **Fig. S23.** Local resolution-boosted spectra of ethyl group carbons from residues A and B |
| 27 | **Fig. S24.** Local resolution-boosted spectra of ethyl group protons from residue A |
| 28 | **Fig. S25.** Local resolution-boosted spectra of ethyl group protons from residue B |
| 29 | **Table S1.** Chemical shifts of $^{1}$H and $^{13}$C atoms in sugar rings of per-*O*-ethylated carrageenans |
| 30 | **Table S2.** Chemical shifts of $^{1}$H and $^{13}$C atoms in ethyl groups of per-*O*-ethylated carrageenans |
| 31 | **Fig. S26.** 2D HSQC NMR spectrum of per-*O*-ethylated F60 isolated from *Mazzaella japonica* |
| 32 | **Fig. S27.** 2D COSY NMR spectrum of per-*O*-ethylated F60 isolated from *Mazzaella japonica* |
| 33 | **Fig. S28.** 2D TOCSY NMR spectrum of per-*O*-ethylated F60 isolated from *Mazzaella japonica* |
| 34, 35 | **Fig. S29.** 2D HMBC NMR spectrum of per-*O*-ethylated F60 isolated from *Mazzaella japonica* |
| 36, 37 | **Fig. S30.** 2D NMR region (ethyl group) of per-*O*-ethylated F60 isolated from *Mazzaella japonica* |
| 38 | **Fig. S31.** 2D HSQC spectrum (low contour threshold) of per-*O*-ethylated F60 |
| 39–41 | **Figs. S32, S33; Table S3.** GC-based monosaccharide analysis of F60 from *Mazzaella japonica* |
| 42–46 | **Figs. S34–S37; Table S4.** GC–MS-based linkage analysis of F60 from *Mazzaella japonica* |

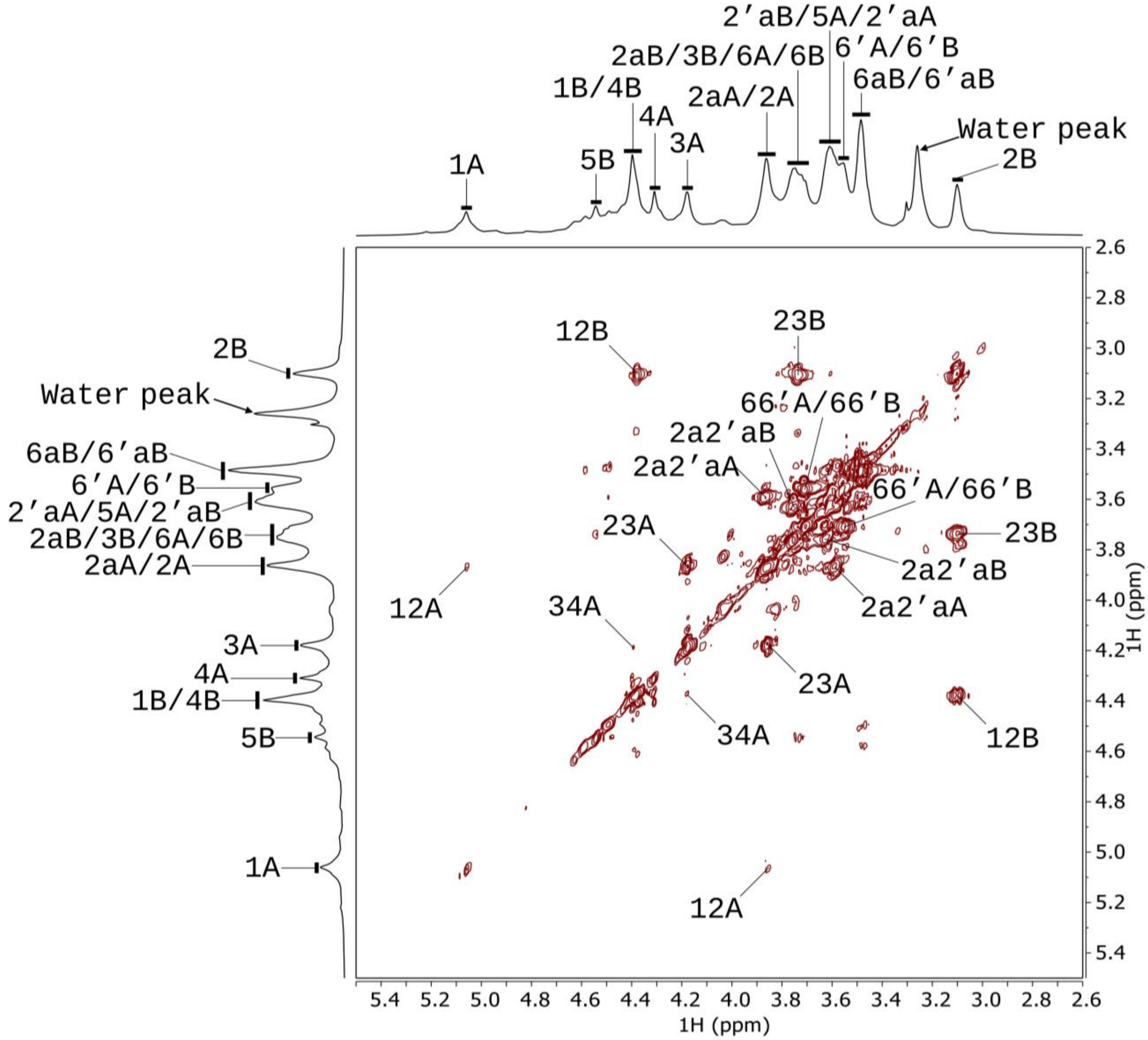


**Fig. S1.** 2D $^{1}$H–$^{1}$H COSY NMR spectrum (700 MHz, 323 K) of DMSO-$d_6$ solution of per-*O*-ethylated commercial kappa-carrageenan. Residues A and B represent 4-linked 2-*O*-ethyl-3,6-anhydro-α-D-galactopyranose and 3-linked 2,6-di-*O*-ethyl-4-*O*-sulfo-β-D-galactopyranose in per-*O*-ethylated kappa-carrageenan, respectively. Cross peaks corresponding to correlations between neighboring protons are marked. For example, the cross-peak "12A" refers to the correlation between H-1 and H-2 of residue A. The lowercase "a" indicates a signal from the methylene group of the ethyl group, and the number before it indicates the location of the ethyl group on the sugar ring. Signals separated by the solidus (/) symbol indicate overlapping. For example, "66'A/66'B" refers to the overlapping of signals 66'A and 66'B. $^{1}$H chemical shifts were internally referenced to TSP-$d_4$ as 0 ppm.

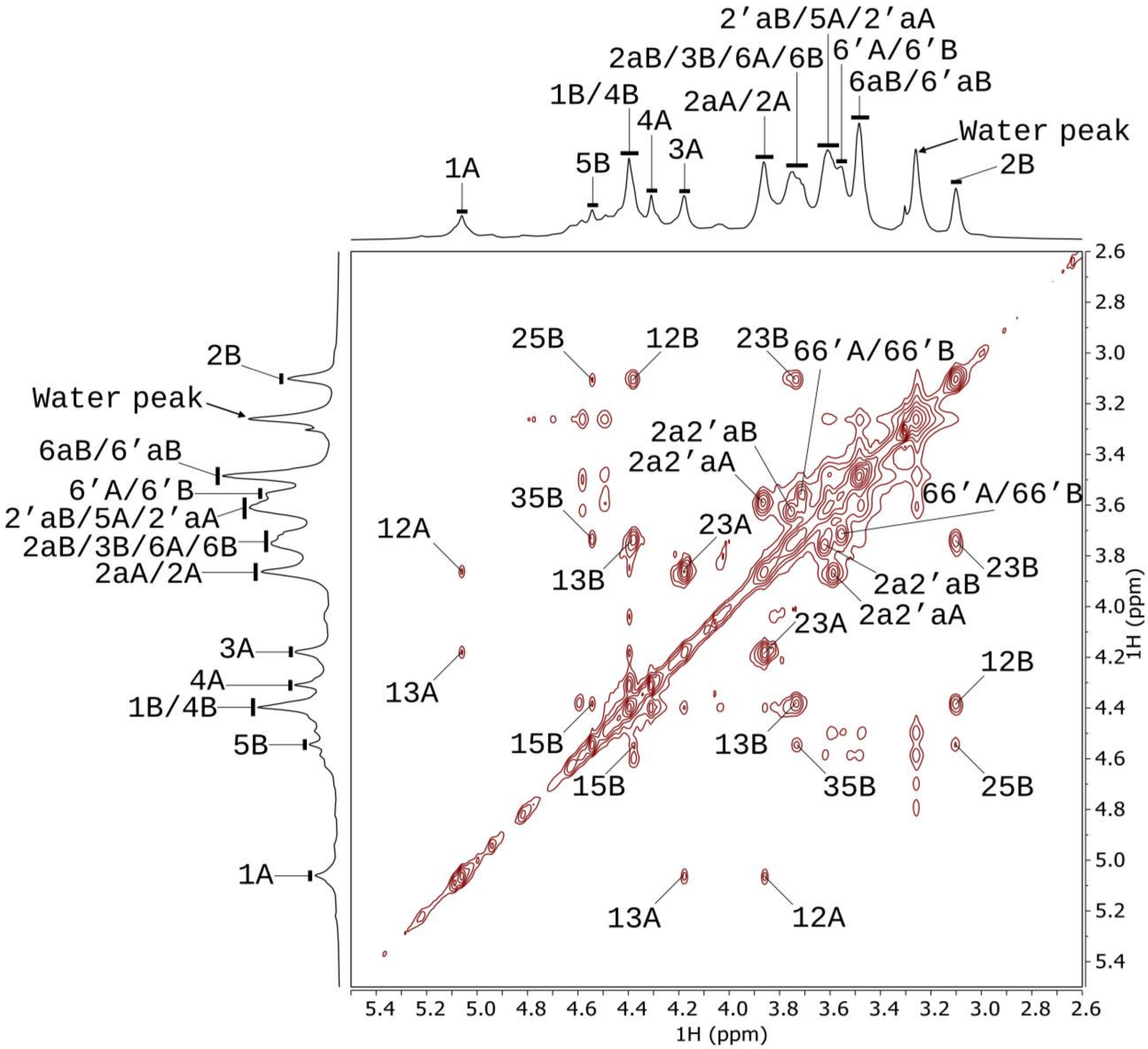


**Fig. S2**. 2D $^{1}$H–$^{1}$H TOCSY NMR spectrum (700 MHz, 323 K) of DMSO-$d_6$ solution of per-*O*-ethylated commercial kappa-carrageenan. Residues A and B represent 4-linked 2-*O*-ethyl-3,6-anhydro-α-D-galactopyranose and 3-linked 2,6-di-*O*-ethyl-4-*O*-sulfo-β-D-galactopyranose in per-*O*-ethylated kappa-carrageenan, respectively. Cross peaks corresponding to correlations between neighboring protons are marked. For example, the cross-peak "13A" refers to the correlation between H-1 and H-3 of residue A. The lowercase "a" indicates a signal from the methylene group of the ethyl group, and the number before it indicates the location of the ethyl group on the sugar ring. Signals separated by the solidus (/) symbol indicate overlapping. For example, "66'A/66'B" refers to the overlapping of signals 66'A and 66'B. $^{1}$H chemical shifts were internally referenced to TSP-$d_4$ as 0 ppm.

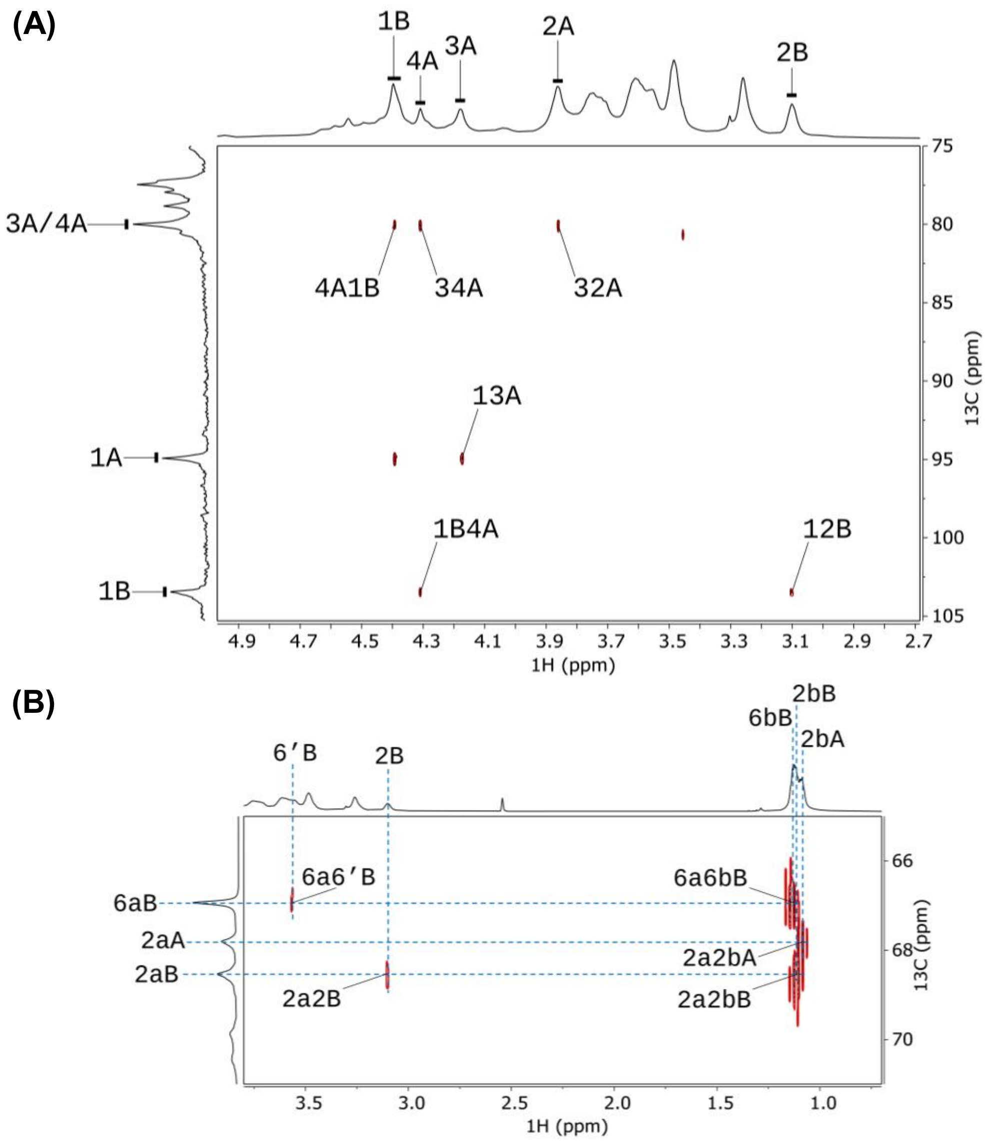


**Fig. S3**. Two regions of 2D $^{1}$H–$^{13}$C HMBC NMR spectrum (700 MHz, 323 K) of DMSO-$d_6$ solution of per-*O*-ethylated commercial kappa-carrageenan. Residues A and B represent 4-linked 2-*O*-ethyl-3,6-anhydro-α-D-galactopyranose and 3-linked 2,6-di-*O*-ethyl-4-*O*-sulfo-β-D-galactopyranose in per-*O*-ethylated kappa-carrageenan, respectively. $^{1}$H–$^{13}$C cross-signals within the same sugar ring (intra-ring) and across glycosidic linkages (inter-ring) are marked. For example, in region A, "13A" refers to the intra-ring correlation between C-1 and H-3 of residue A, while "4A1B" indicates the inter-ring correlation between C-4 of residue A and H-1 of residue B. The lowercase "a" and "b" indicate relevant $^{1}$H or $^{13}$C signals from the methylene group and the methyl group of the ethyl group, and the number before "a" and "b" indicates the location of the ethyl group on the sugar ring. For example, the cross signal "2a2bA" in the region B indicates the correlation between the methylene $^{13}$C and the methyl proton in the ethyl group attached to the *O*-2 position of residue A. Signals separated by the solidus (/) symbol indicate overlapping. $^{1}$H and $^{13}$C chemical shifts were internally referenced to TSP-$d_4$ as 0 ppm.

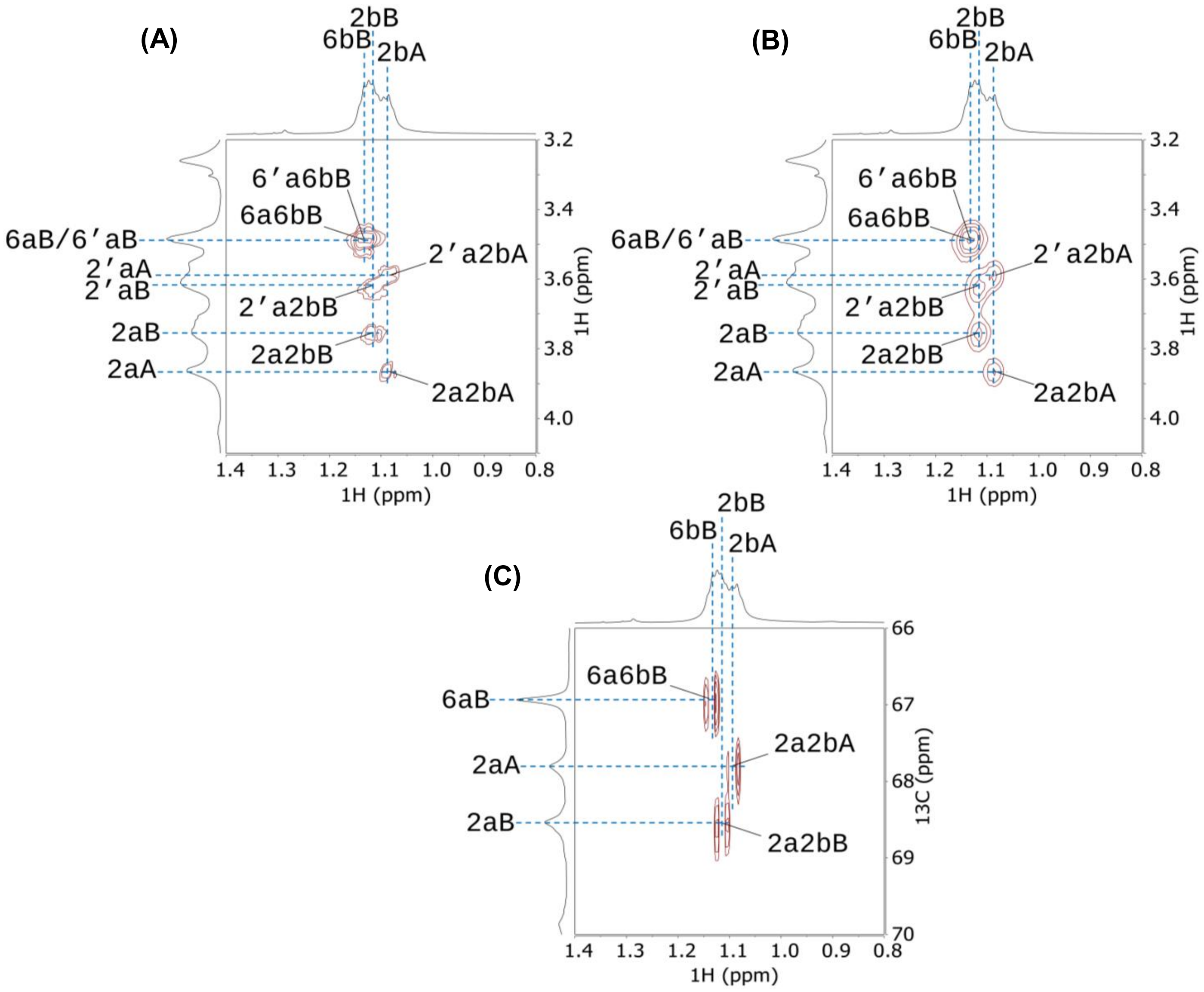


**Fig. S4.** Regions of (A) 2D $^{1}H$–$^{1}H$ COSY, (B) 2D $^{1}H$–$^{1}H$ TOCSY, and (C) 2D $^{1}H$–$^{13}C$ HMBC spectra (700 MHz, 323 K) of DMSO-$d_6$ solution of per-*O*-ethylated commercial kappa-carrageenan, showing correlations between the atoms of the methylene group and the methyl group in the ethyl group attached to different positions on the sugar ring. Residues A and B represent 4-linked 2-*O*-ethyl-3,6-anhydro-α-D-galactopyranose and 3-linked 2,6-di-*O*-ethyl-4-*O*-sulfo-β-D-galactopyranose in per-*O*-ethylated kappa-carrageenan, respectively. The lowercase "a" and "b" indicate relevant $^{1}H$ or $^{13}C$ signals from the methylene group and the methyl group of the ethyl group, and the number before "a"

and “b” indicates the location of the ethyl group on the sugar ring. For example, the cross signals “2a2bA” in the COSY and TOCSY spectra indicate the correlation between the methylene proton and the methyl proton in the ethyl group attached to the *O*-2 position of residue A, while in the HMBC spectrum, it indicates the correlation between the methylene $^{13}C$ and the methyl proton in the ethyl group attached to the *O*-2 position of residue A. $^{1}H$ and $^{13}C$ chemical shifts were internally referenced to TSP-$d_4$ as 0 ppm.

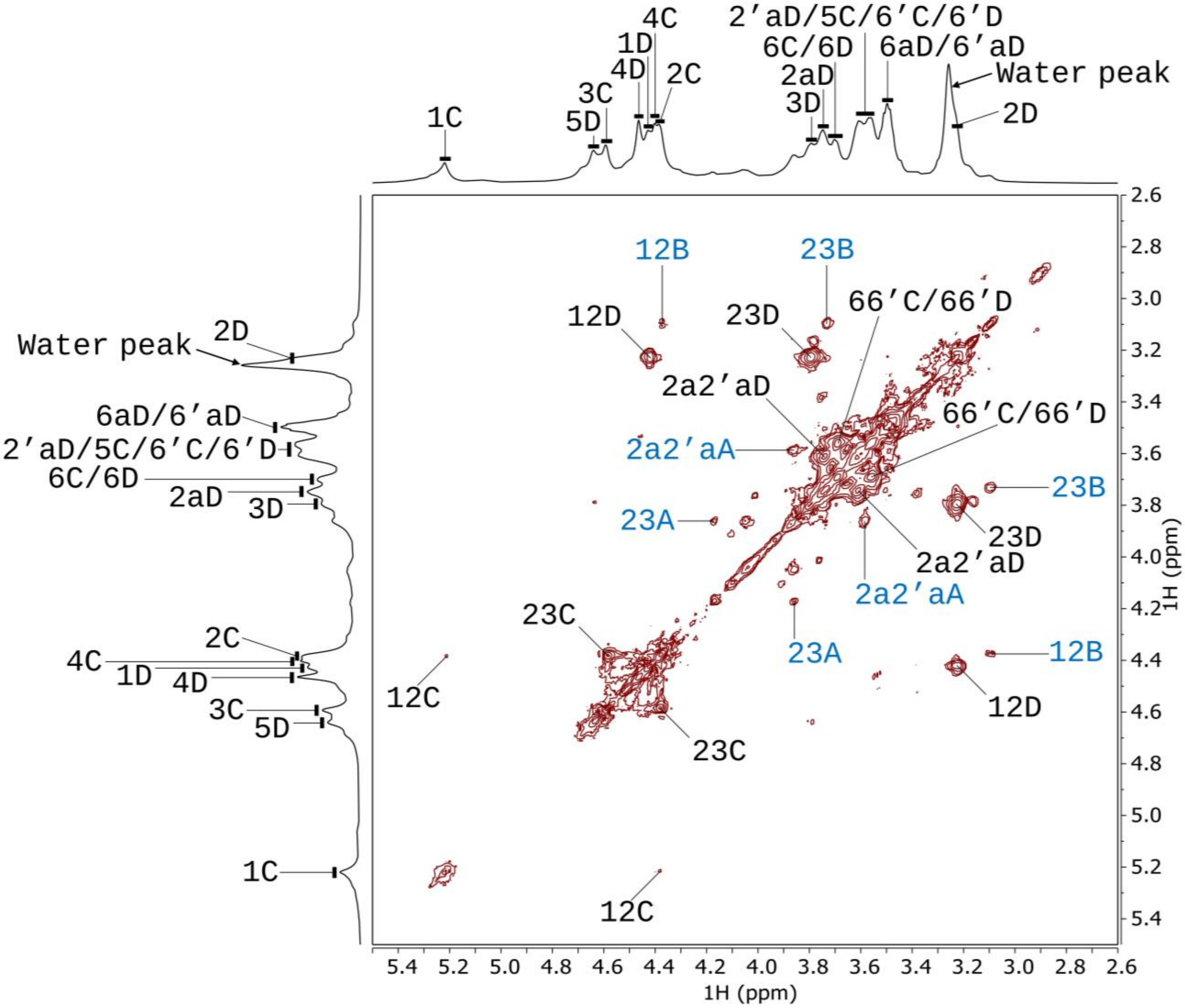


**Fig. S5**. 2D $^{1}$H–$^{1}$H COSY NMR spectrum (700 MHz, 323 K) of DMSO-$d_6$ solution of per-*O*-ethylated commercial iota-carrageenan. Residues C and D represent 4-linked 2-*O*-sulfo-3,6-anhydro-α-D-galactopyranose and 3-linked 2,6-di-*O*-ethyl-4-*O*-sulfo-β-D-galactopyranose in per-*O*-ethylated iota-carrageenan, respectively. Residues A and B represent 4-linked 2-*O*-ethyl-3,6-anhydro-α-D-galactopyranose and 3-linked 2,6-di-*O*-ethyl-4-*O*-sulfo-β-D-galactopyranose in per-*O*-ethylated product of minor level of kappa-carrageenan in the commercial sample, respectively. Labels corresponding to the minor kappa-carrageenan structures are shown in blue. Cross peaks corresponding to correlations between neighboring protons are marked. For example, the cross-peak “12C” refers to the correlation between H-1 and H-2 of residue C. The lowercase “a” indicates a signal from the methylene group of the ethyl group, and the number before it indicates the location of the ethyl group on the sugar ring. Signals separated by the solidus (/) symbol indicate overlapping. For example, “66’C/66’D” refers to the overlapping of signals 66’C and 66’D. $^{1}$H chemical shifts were internally referenced to TSP-$d_4$ as 0 ppm.

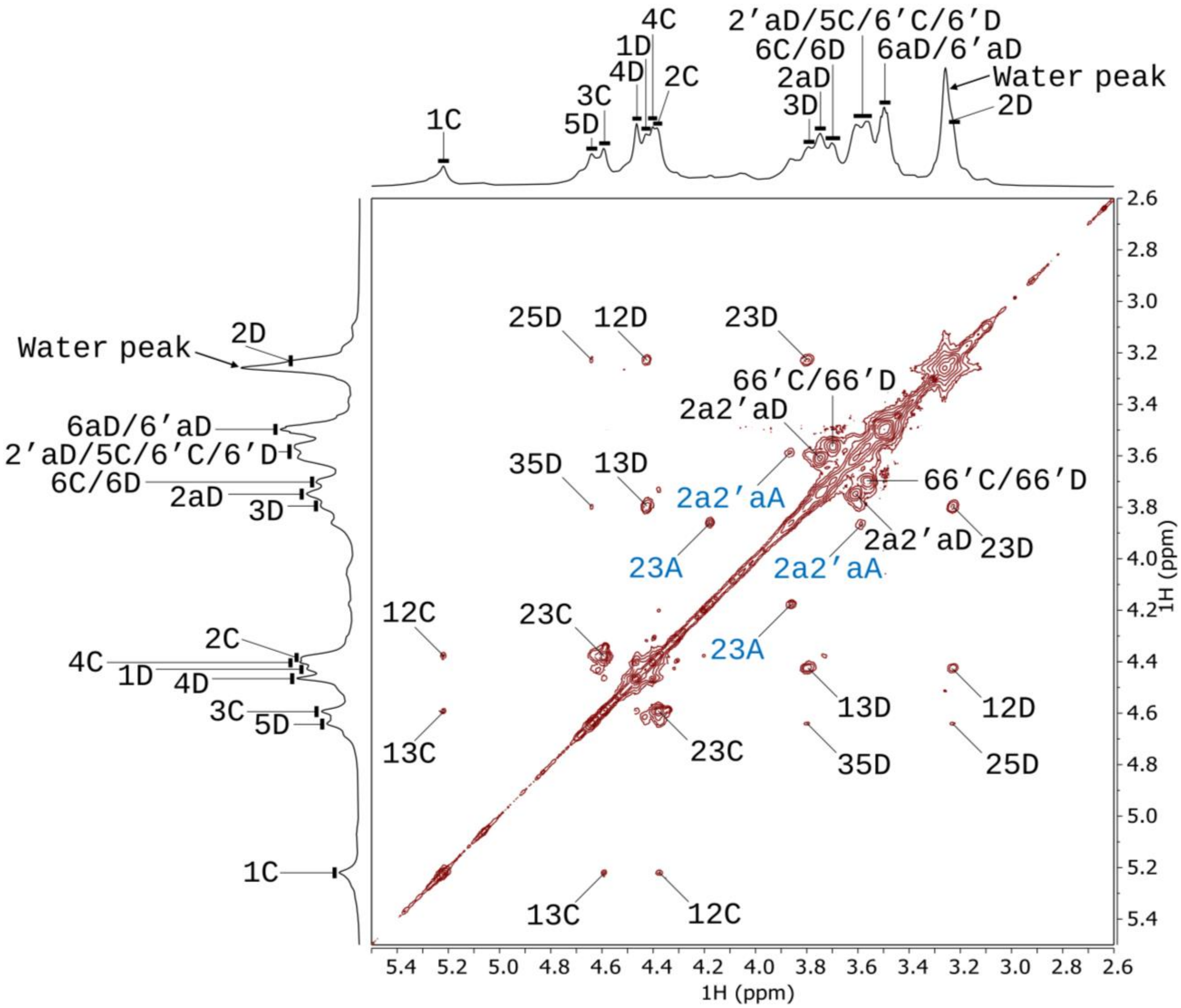


**Fig. S6**. 2D $^{1}$H–$^{1}$H TOCSY NMR spectrum (700 MHz, 323 K) of DMSO-$d_6$ solution of per-*O*-ethylated commercial iota-carrageenan. Residues C and D represent 4-linked 2-*O*-sulfo-3,6-anhydro-α-D-galactopyranose and 3-linked 2,6-di-*O*-ethyl-4-*O*-sulfo-β-D-galactopyranose in per-*O*-ethylated iota-carrageenan, respectively. Residues A and B represent 4-linked 2-*O*-ethyl-3,6-anhydro-α-D-galactopyranose and 3-linked 2,6-di-*O*-ethyl-4-*O*-sulfo-β-D-galactopyranose in per-O-ethylated product of minor level of kappa-carrageenan in the commercial sample, respectively. Labels corresponding to the minor kappa-carrageenan structures are shown in blue. Cross peaks corresponding to correlations between neighboring protons are marked. For example, the cross-peak “13C” refers to the correlation between H-1 and H-3 of residue C. The lowercase “a” indicates a signal from the methylene group of the ethyl group, and the number before it indicates the location of the ethyl group on the sugar ring. Signals separated by the solidus (/) symbol indicate overlapping. For example, “66’C/66’D” refers to the overlapping of signals 66’C and 66’D. $^{1}$H chemical shifts were internally referenced to TSP-$d_4$ as 0 ppm.

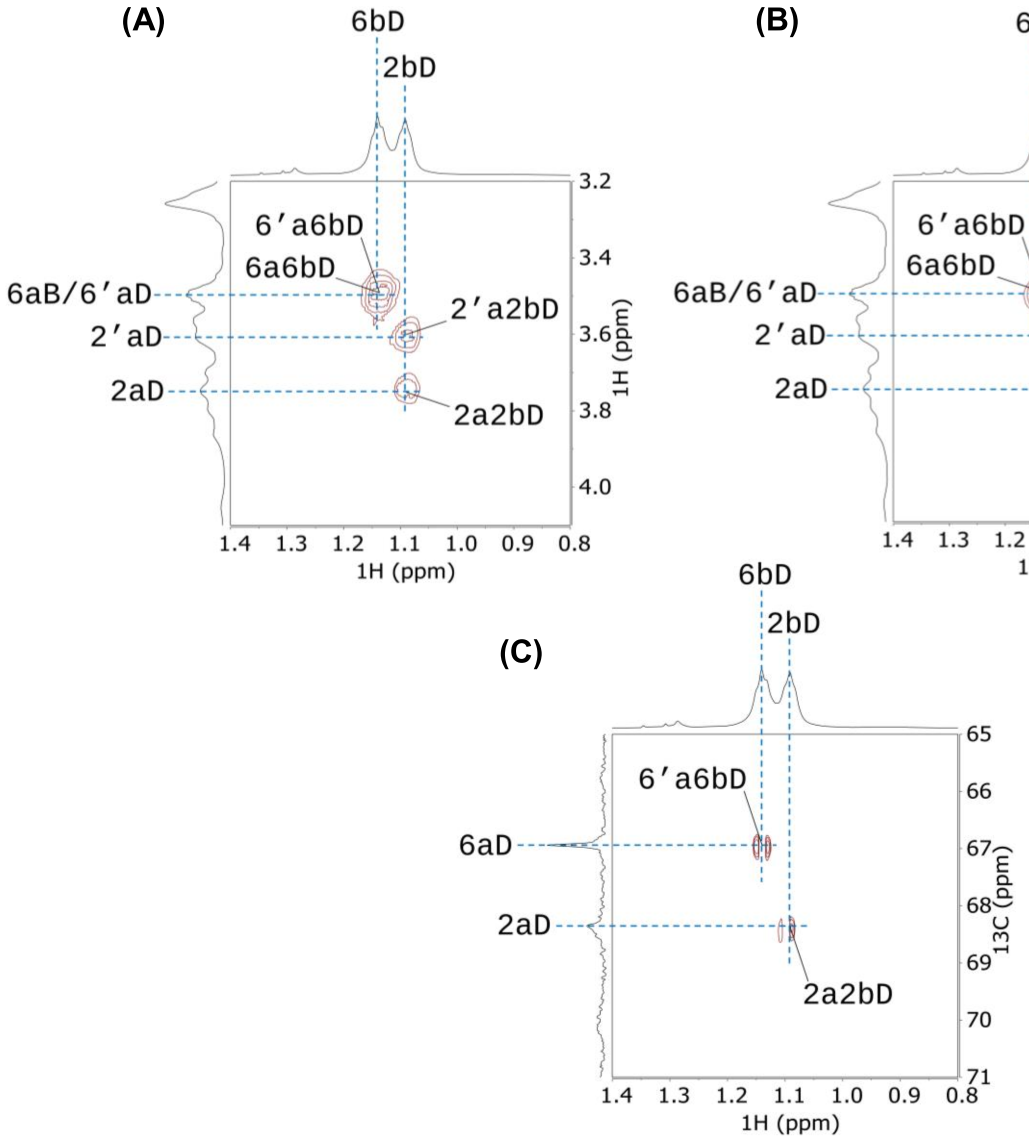


**Fig. S7.** Regions of (A) 2D $^{1}H$–$^{1}H$ COSY, (B) 2D $^{1}H$–$^{1}H$ TOCSY, and (C) 2D $^{1}H$–$^{13}C$ HMBC spectra (700 MHz, 323 K) of DMSO-$d_6$ solution of per-*O*-ethylated commercial iota-carrageenan. Residues C and D represent 4-linked 2-*O*-sulfo-3,6-anhydro-α-D-galactopyranose and 3-linked 2,6-di-*O*-ethyl-4-*O*-sulfo-β-D-galactopyranose in per-*O*-ethylated iota-carrageenan, respectively. Residues A and B represent 4-linked 2-*O*-ethyl-3,6-anhydro-α-D-galactopyranose and 3-linked 2,6-di-*O*-ethyl-4-*O*-sulfo-β-D-galactopyranose in per-*O*-ethylated product of minor level of kappa-carrageenan in the commercial sample, respectively. The lowercase “a” and “b” indicate relevant $^{1}H$ or $^{13}C$ signals from the methylene group and the methyl group of the ethyl group, and the number before “a” and “b” indicates the location of the ethyl group on the sugar ring. For example, the cross signals “2a2bA” in the COSY and TOCSY spectra indicate the correlation between the methylene proton and the methyl proton in the ethyl group attached to the *O*-2 position of residue A, while in the HMBC spectrum, it indicates the correlation between the methylene $^{13}C$ and the methyl proton in the ethyl group attached to the *O*-2 position of residue A. $^{1}H$ and $^{13}C$ chemical shifts were internally referenced to TSP-$d_4$ as 0 ppm.

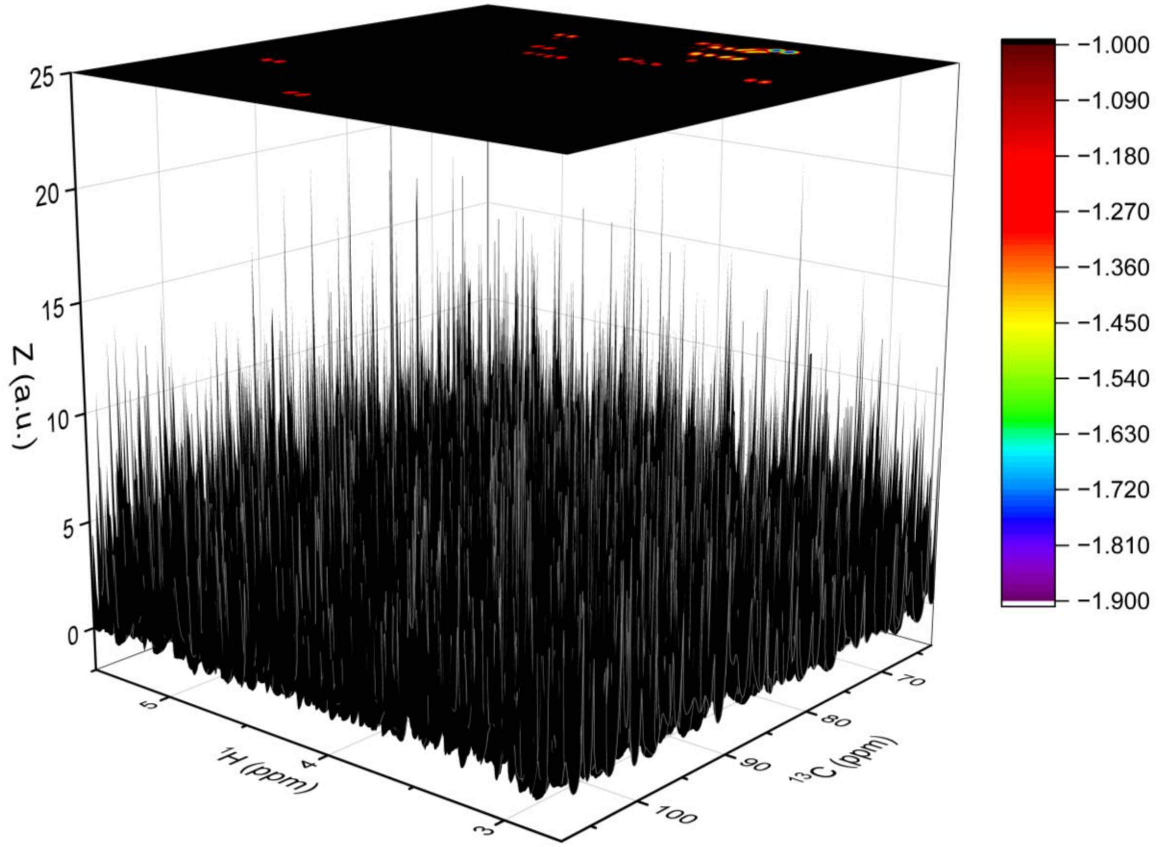


**Fig. S8.** Side-view dark forest image (3D colormap surface plot) of 2D $^{1}$H–$^{13}$C HSQC data of per-*O*-ethylated commercial kappa-carrageenan after first derivative processing and logarithmic cosine transformation (FDP–LCT) along the $^{1}$H dimension, with the surface generated from *Z* values corresponding to $^{1}$H and $^{13}$C chemical shifts, the color scale displayed on the right, and the corresponding 2D contour map projection shown above.

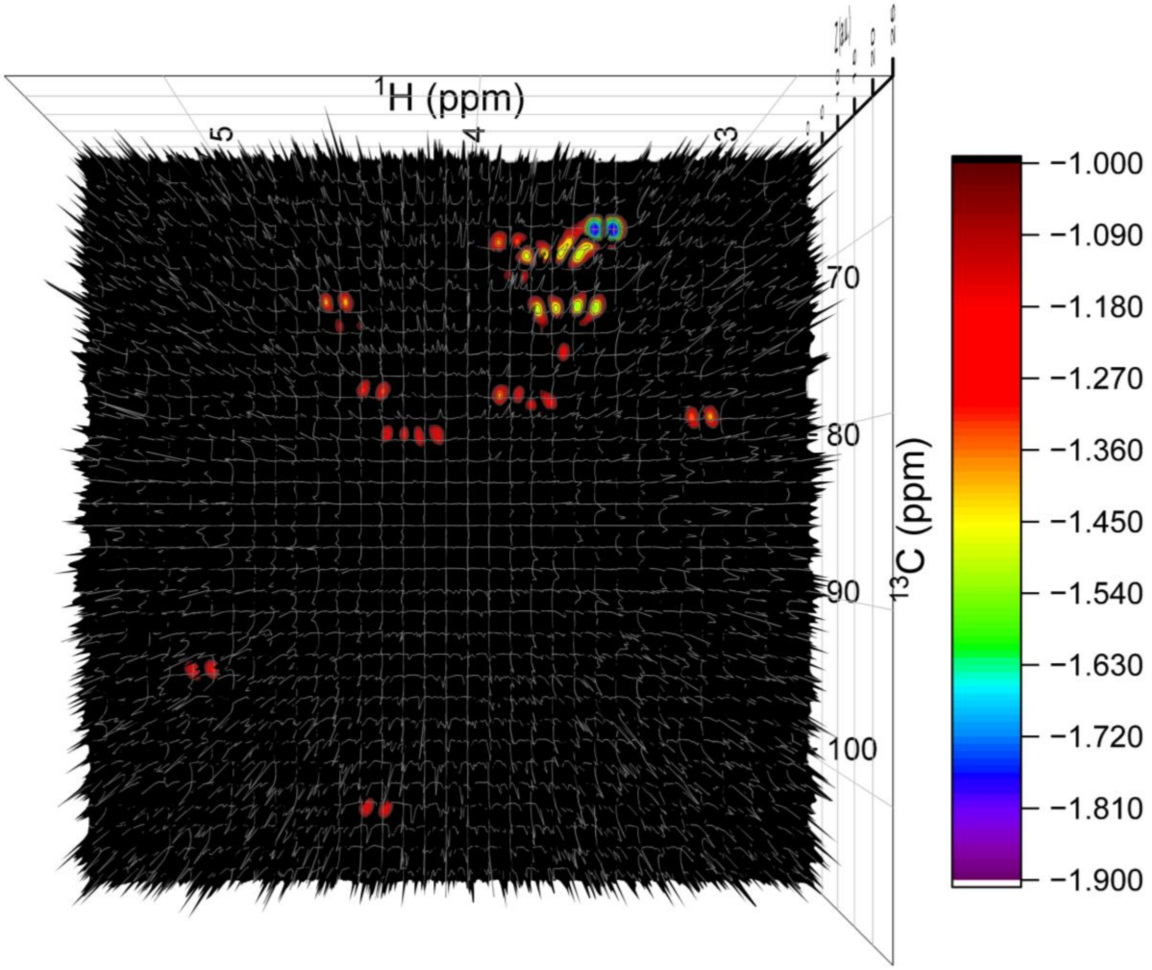


**Fig. S9**. Top-view dark forest image (3D colormap surface plot) of 2D $^{1}$H–$^{13}$C HSQC data of per-*O*-ethylated commercial kappa-carrageenan after FDP–LCT along the $^{1}$H dimension, with the surface generated from *Z* values corresponding to $^{1}$H and $^{13}$C chemical shifts and the color scale displayed on the right.

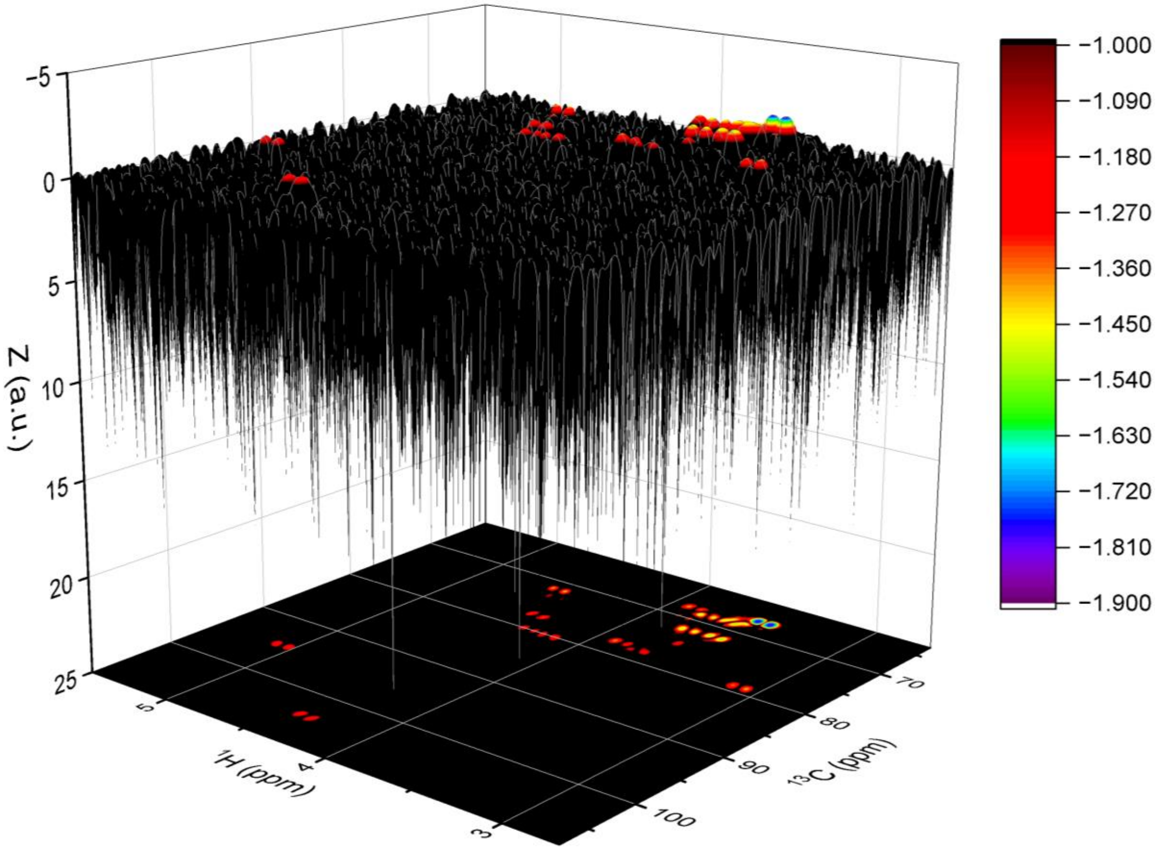


**Fig. S10.** Inverted side-view dark forest image (3D colormap surface plot) of 2D $^{1}H$–$^{13}C$ HSQC data of per-*O*-ethylated commercial kappa-carrageenan after FDP–LCT along the $^{1}H$ dimension. The surface is generated from *Z* values corresponding to $^{1}H$ and $^{13}C$ chemical shifts, with the color scale displayed on the right and the corresponding 2D contour map projection shown below.

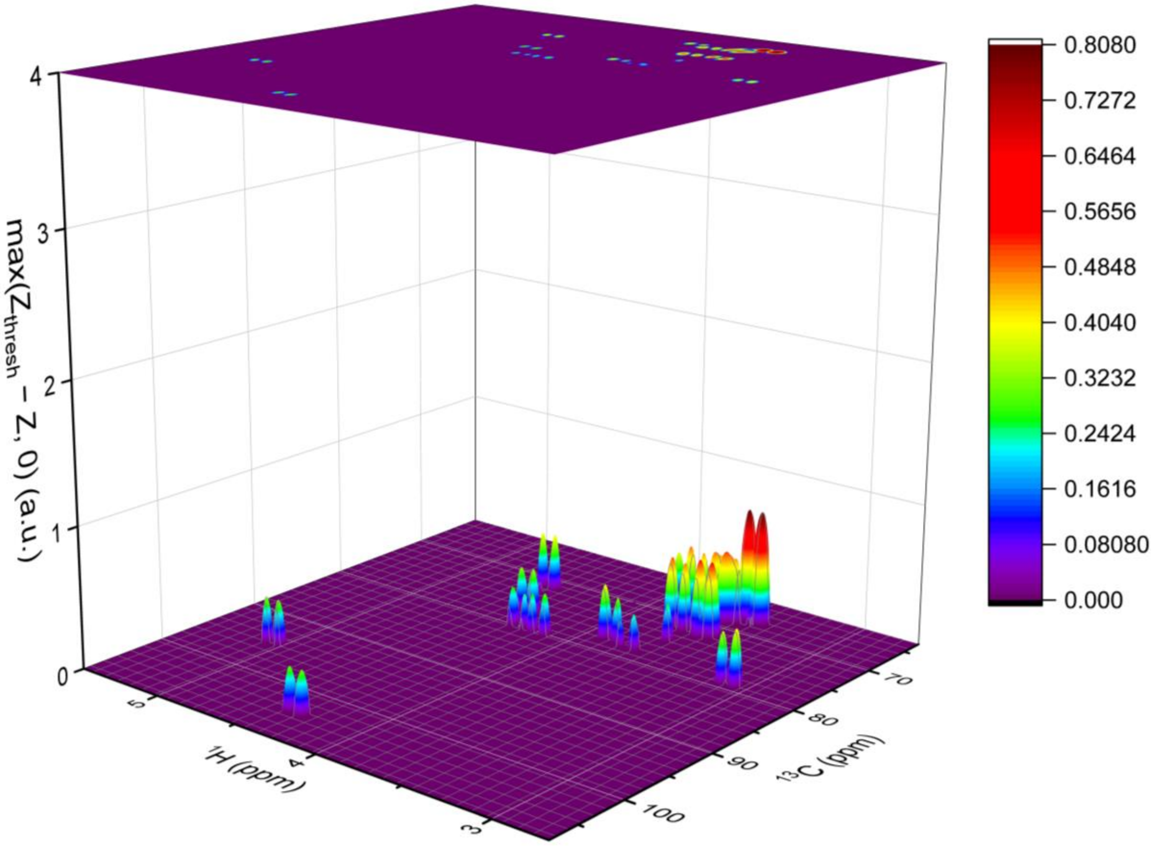


**Fig. S11.** 3D colormap surface plot of 2D [1]H–[13]C HSQC data of per-*O*-ethylated commercial kappa-carrageenan after FDP–LCT along the [1]H dimension, with the surface generated from $\max(\mathrm{Z_{thresh}} - Z, 0)$ corresponding to [1]H and [13]C chemical shifts, the color scale displayed on the right, and the corresponding 2D contour map projection shown above. $\mathrm{Z_{thresh}}$ is an empirically optimized threshold value, and a $\mathrm{Z_{thresh}}$ of 1.0 was used for plotting this figure.

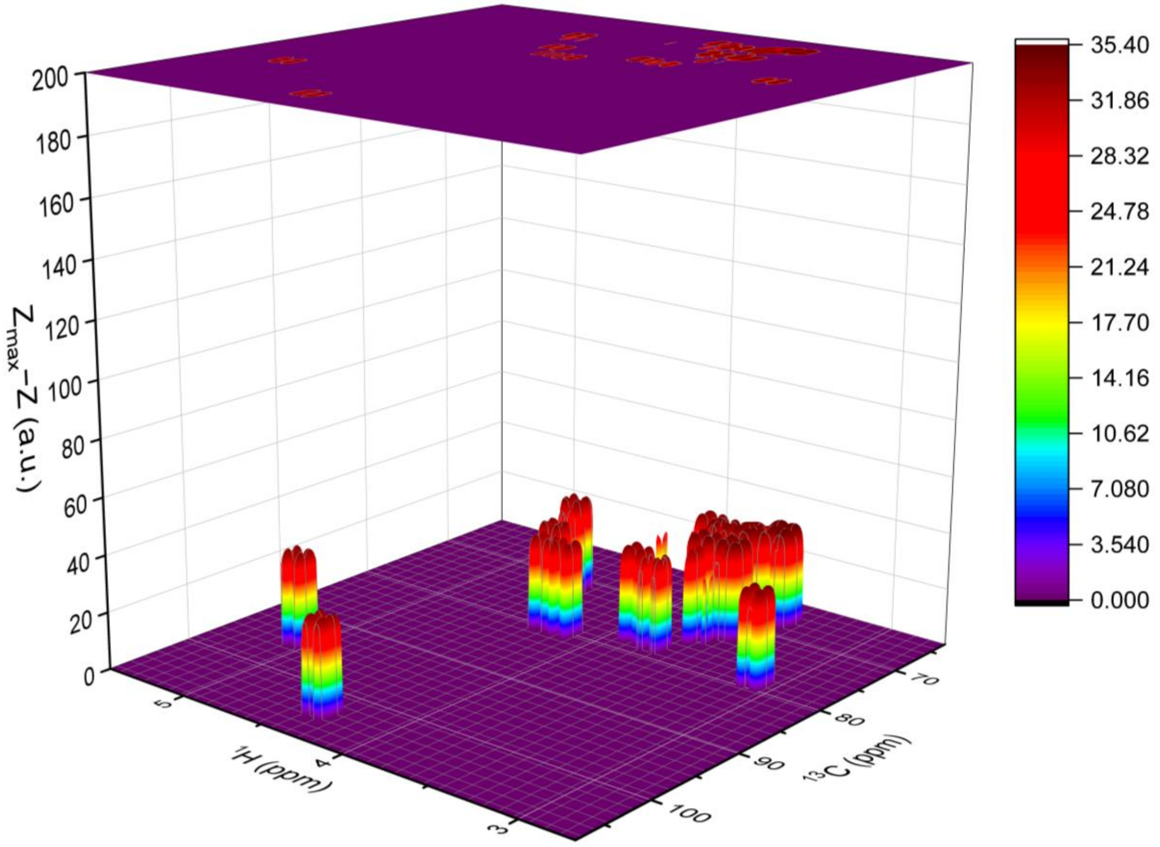


**Fig. S12.** 3D colormap surface plot of 2D $^{1}$H–$^{13}$C HSQC data of per-*O*-ethylated commercial kappa-carrageenan after FDP–LCT along the $^{13}$C dimension to the processed data shown in Fig. S11. The surface is generated from $Z_{max} - Z$ corresponding to $^{1}$H and $^{13}$C chemical shifts, with the color scale displayed on the right and the corresponding 2D contour map projection shown above. $Z_{max}$ is a constant set as 36.7368005696771, as detailed in Section 2 (Theory).

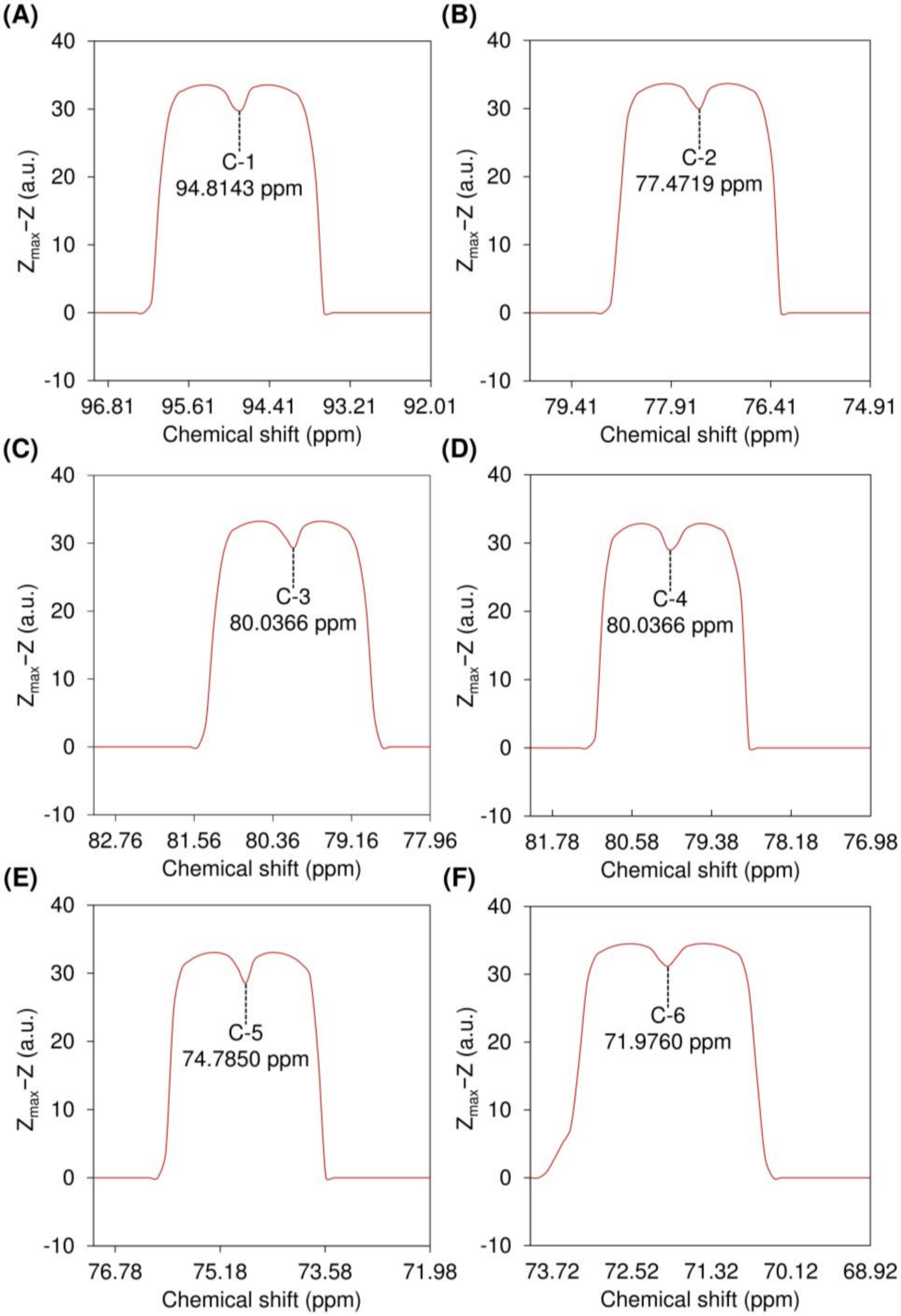


**Fig. S13.** Local resolution-boosted spectra of sugar ring carbons from 4-linked 2-*O*-ethyl-3,6-anhydro-α-D-galactopyranose (residue A), extracted from 2D $^{1}H$–$^{13}C$ HSQC data of per-*O*-ethylated commercial kappa-carrageenan after FDP–LCT along the $^{1}H$ dimension followed by the $^{13}C$ dimension. $Z_{max}$ is a constant set as 36.7368005696771, as detailed in Section 2 (Theory).

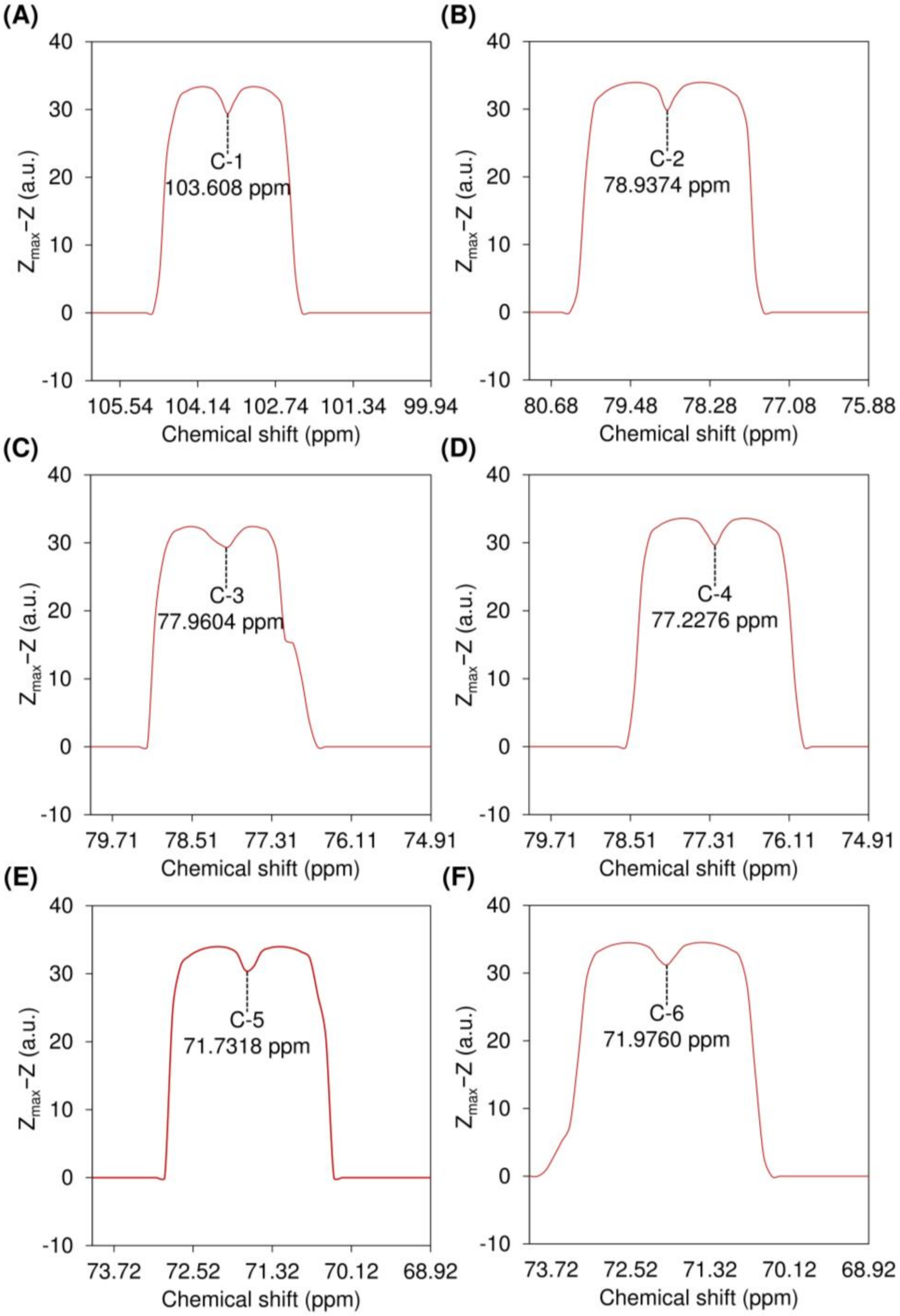


**Fig. S14.** Local resolution-boosted spectra of sugar ring carbons from 3-linked 2,6-di-*O*-ethyl-4-*O*-sulfo-β-D-galactopyranose (residue B), extracted from 2D $^{1}H$–$^{13}C$ HSQC data of per-*O*-ethylated commercial kappa-carrageenan after FDP–LCT along the $^{1}H$ dimension followed by the $^{13}C$ dimension. $Z_{max}$ is a constant set as 36.7368005696771, as detailed in Section 2 (Theory).

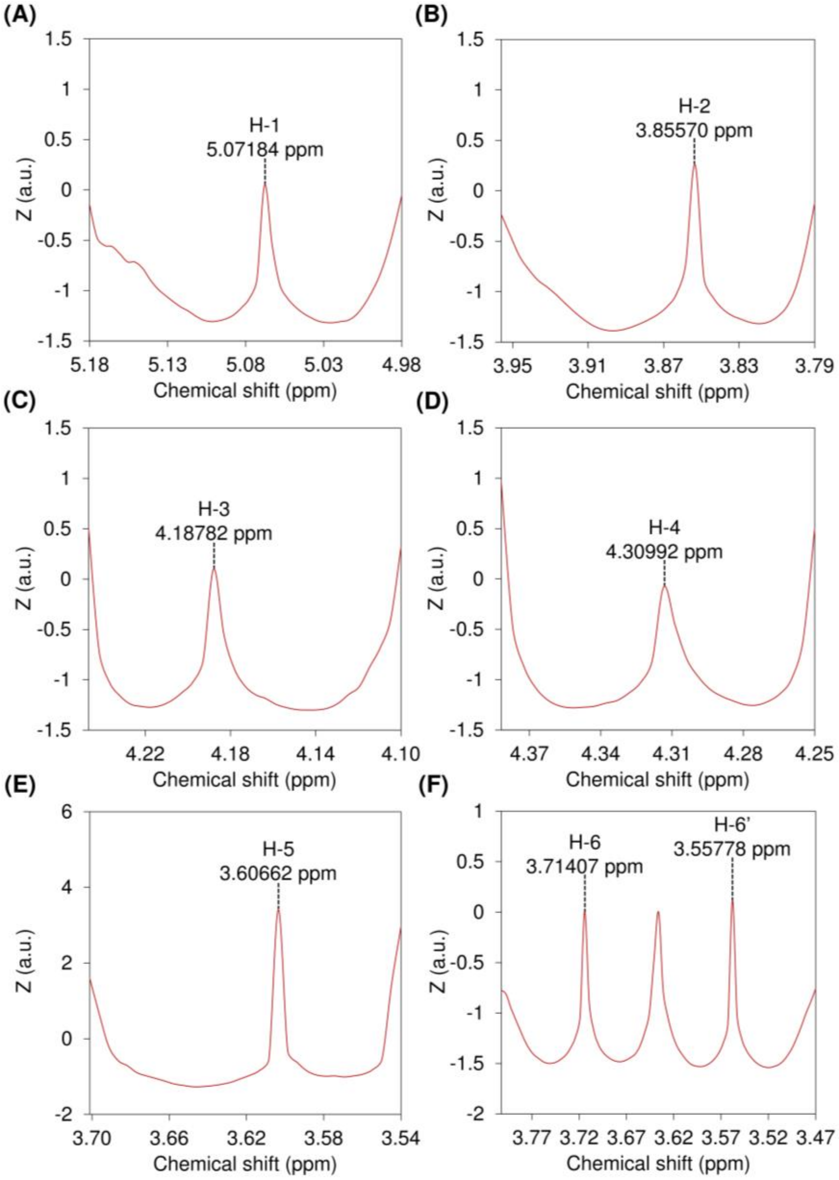


**Fig. S15.** Local resolution-boosted spectra of sugar ring protons from 4-linked 2-*O*-ethyl-3,6-anhydro-α-D-galactopyranose (residue A), extracted from 2D $^{1}H$–$^{13}C$ HSQC data of per-*O*-ethylated commercial kappa-carrageenan after FDP–LCT along the $^{1}H$ dimension.

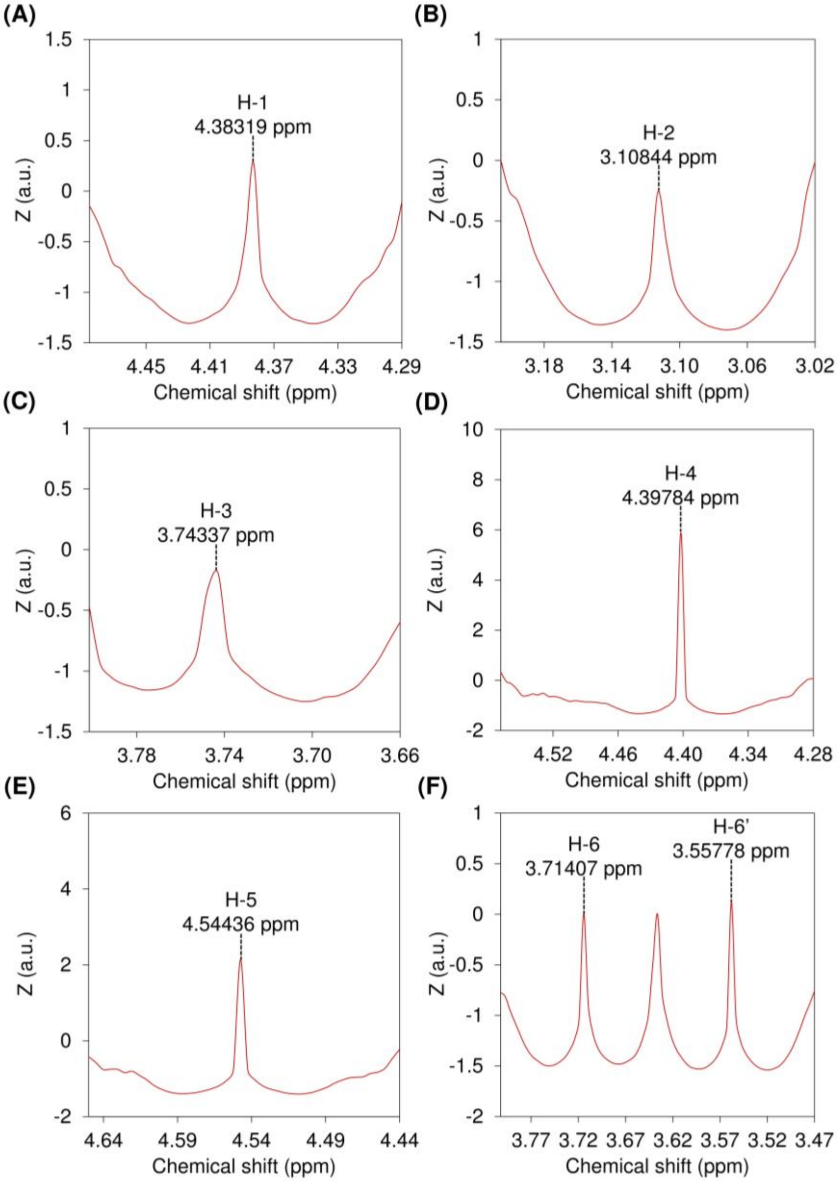


**Fig. S16.** Local resolution-boosted spectra of sugar ring protons from 3-linked 2,6-di-*O*-ethyl-4-*O*-sulfo-β-D-galactopyranose (residue B), extracted from 2D $^{1}$H–$^{13}$C HSQC data of per-*O*-ethylated commercial kappa-carrageenan after FDP–LCT along the $^{1}$H dimension.

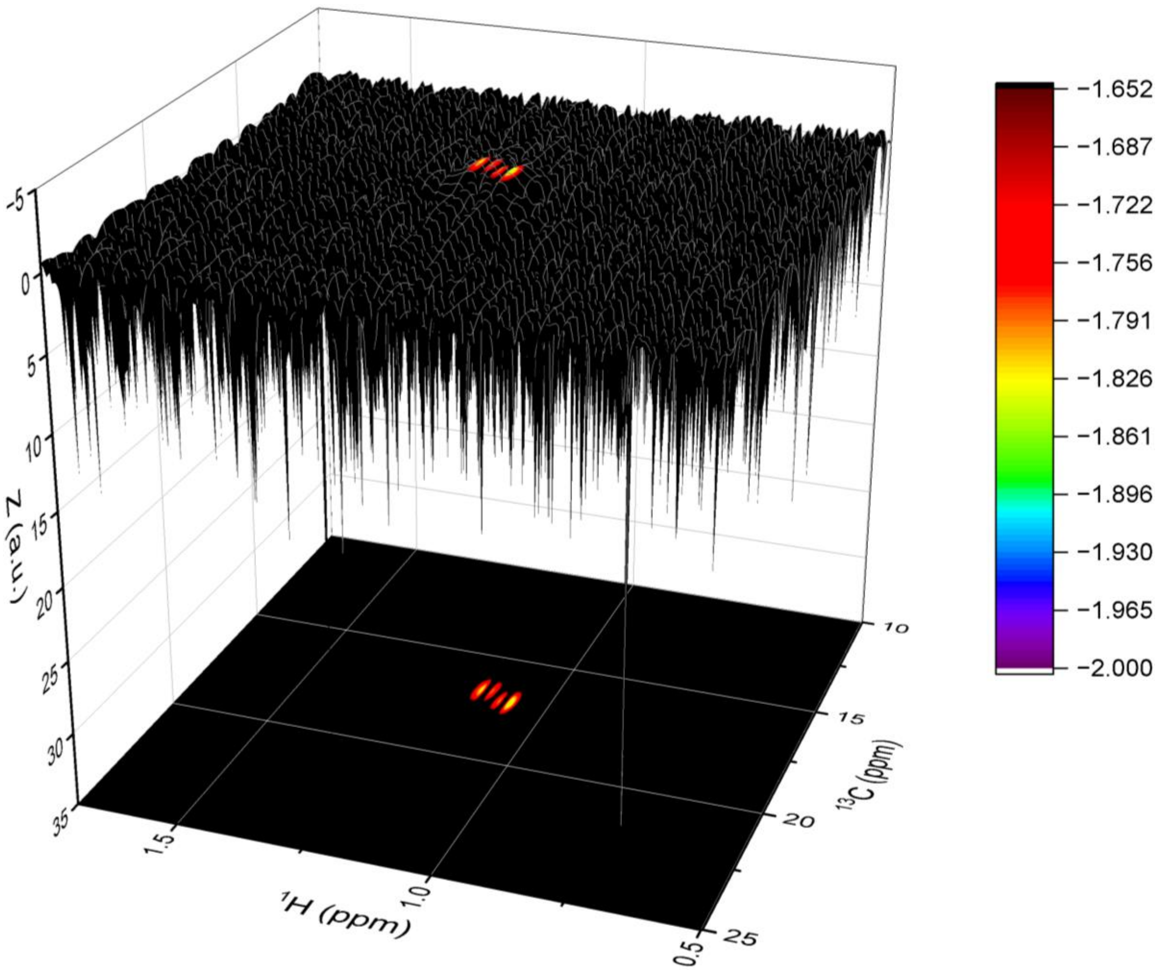


**Fig. S17.** Inverted side-view dark forest image (3D colormap surface plot) of 2D HSQC data showing regions of $^{1}$H–$^{13}$C correlations for the methyl group in the ethyl group of per-*O*-ethylated commercial iota-carrageenan after FDP–LCT along the $^{1}$H dimension. The surface was generated from *Z* values corresponding to $^{1}$H and $^{13}$C chemical shifts, with the color scale displayed on the right and the corresponding 2D contour map projection shown below.

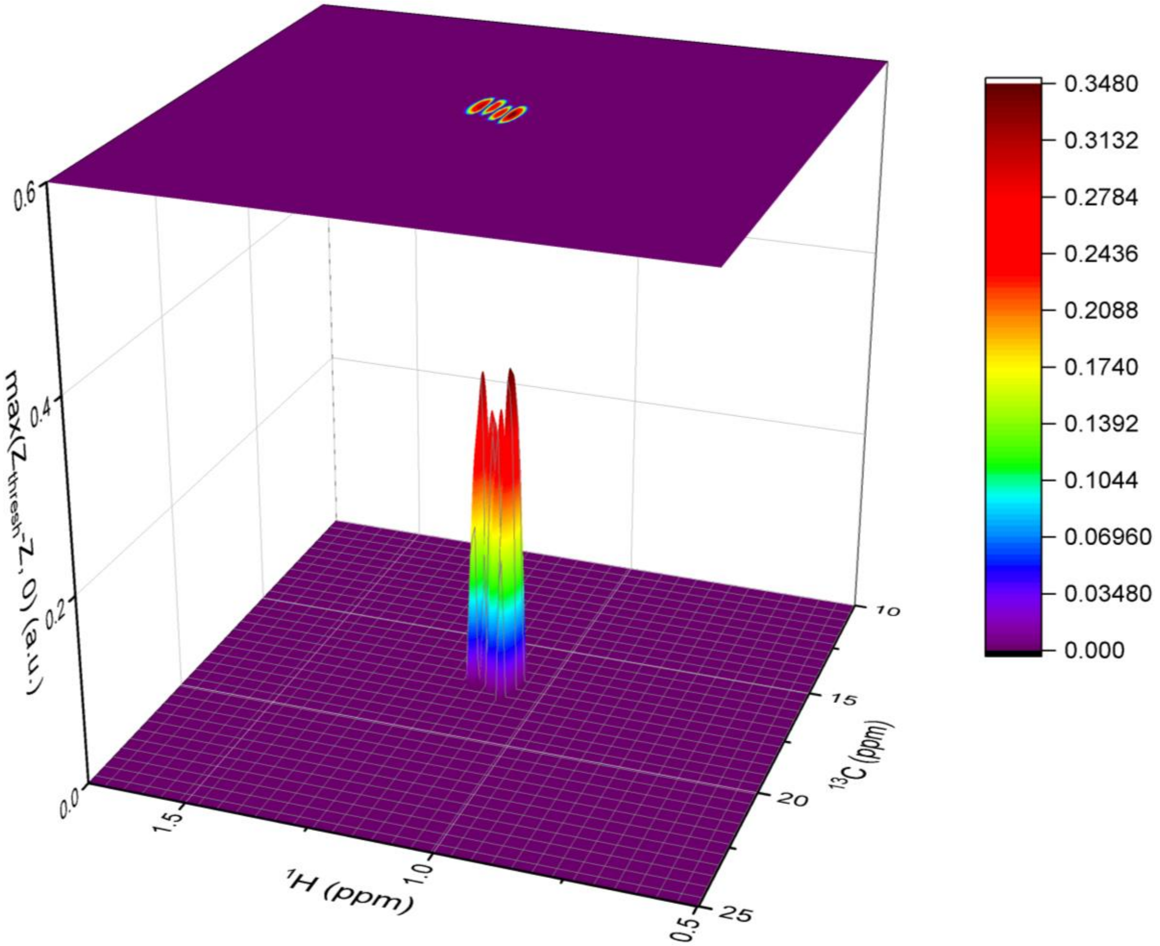


**Fig. S18.** 3D colormap surface plot of 2D HSQC data showing regions of $^{1}$H–$^{13}$C correlations for the methyl group in the ethyl group of per-*O*-ethylated commercial iota-carrageenan after FDP–LCT along the $^{1}$H dimension, with the surface generated from $\max(Z_{\mathrm{thresh}} - Z, 0)$ corresponding to $^{1}$H and $^{13}$C chemical shifts, the color scale displayed on the right, and the corresponding 2D contour map projection shown above. $Z_{\mathrm{thresh}}$ is an empirically optimized threshold value, and a $Z_{\mathrm{thresh}}$ of 0.148 was used for plotting this figure.

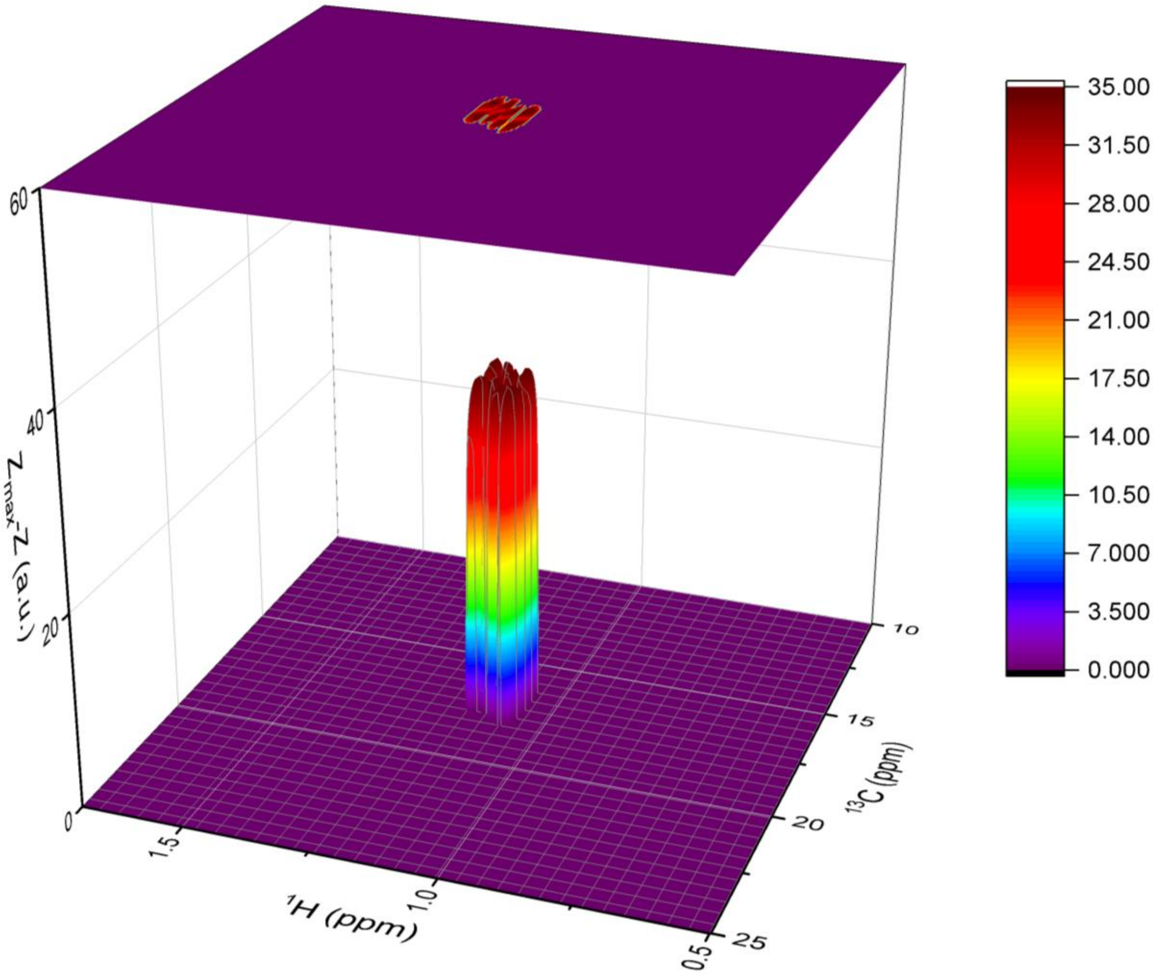


**Fig. S19.** 3D colormap surface plot of 2D HSQC data showing regions of $^1H$–$^{13}C$ correlations for the methyl group in the ethyl group of per-*O*-ethylated commercial iota-carrageenan after FDP–LCT along the $^{13}C$ dimension to the processed data shown in Fig. S18. The surface is generated from $Z_{max} - Z$, corresponding to $^1H$ and $^{13}C$ chemical shifts, with the color scale displayed on the right and the corresponding 2D contour map projection shown above. $Z_{max}$ is a constant set as 36.7368005696771, as detailed in Section 2 (Theory).

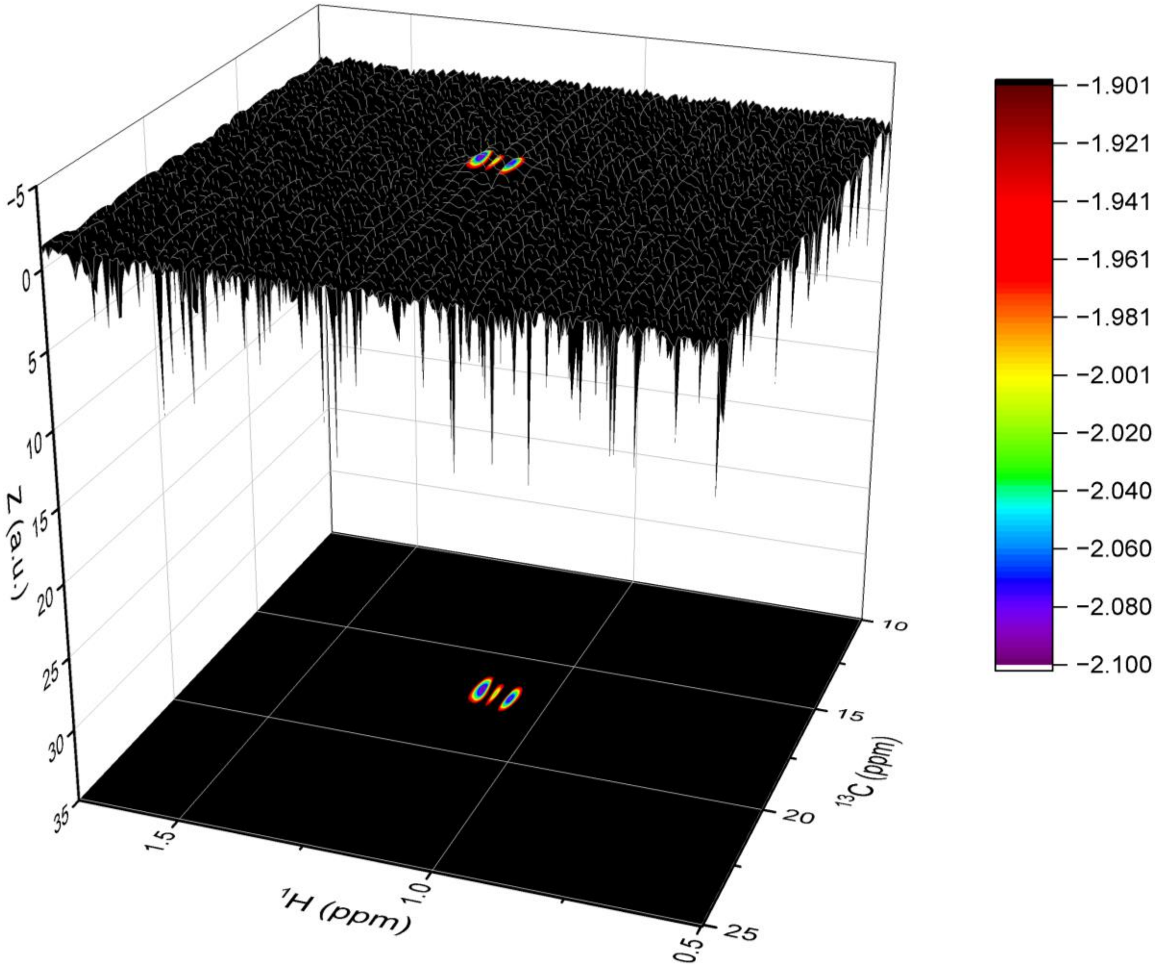


**Fig. S20.** Inverted side-view dark forest image (3D colormap surface plot) of 2D HSQC data showing regions of $^{1}$H–$^{13}$C correlations for the methyl group in the ethyl group of per-*O*-ethylated commercial kappa-carrageenan after FDP–LCT along the $^{1}$H dimension. The surface was generated from *Z* values corresponding to $^{1}$H and $^{13}$C chemical shifts, with the color scale displayed on the right and the corresponding 2D contour map projection shown below.

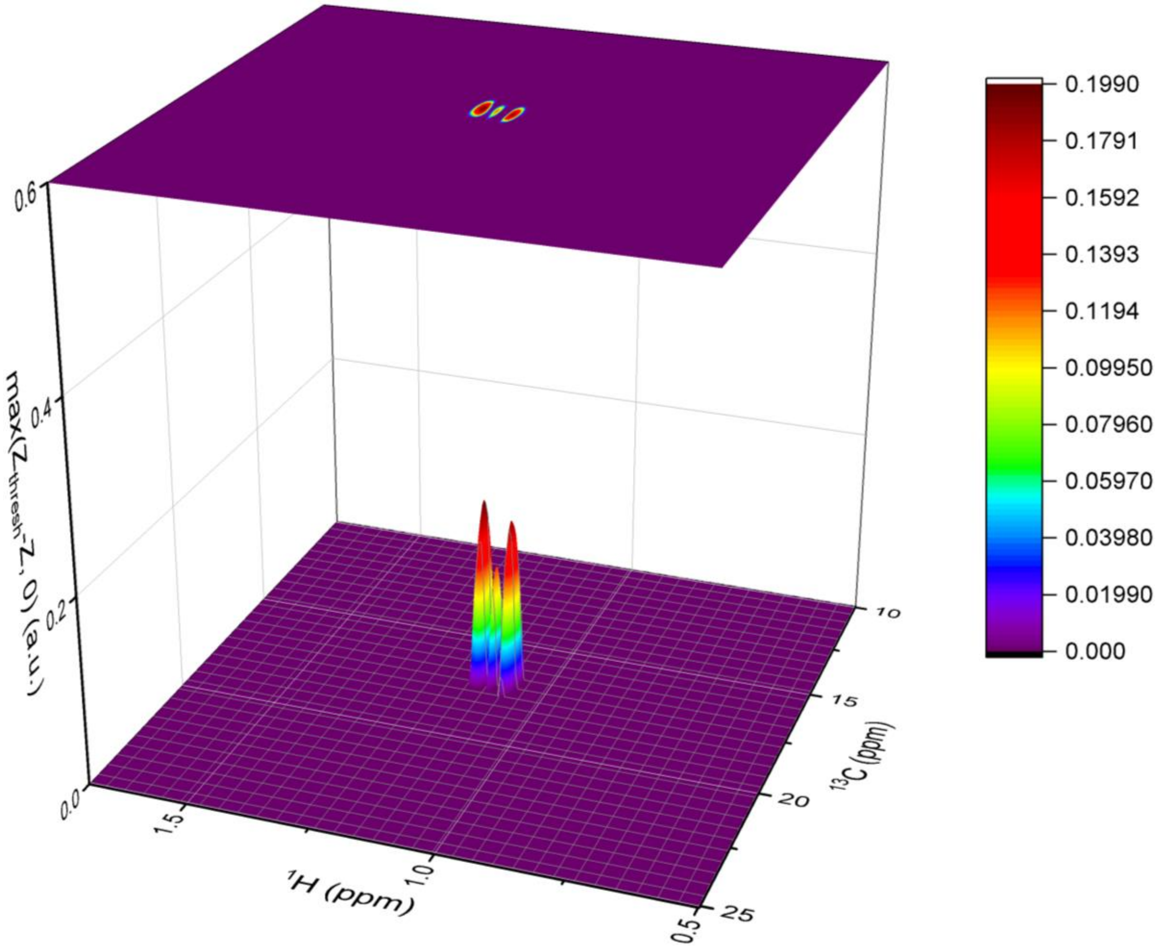


**Fig. S21.** 3D colormap surface plot of 2D HSQC data showing regions of $^{1}H$–$^{13}C$ correlations for the methyl group in the ethyl group of per-*O*-ethylated commercial kappa-carrageenan after FDP–LCT along the $^{1}H$ dimension, with the surface generated from $\max(Z_{\mathrm{thresh}} - Z, 0)$ corresponding to $^{1}H$ and $^{13}C$ chemical shifts, the color scale displayed on the right, and the corresponding 2D contour map projection shown above. $Z_{\mathrm{thresh}}$ is an empirically optimized threshold value, and a $Z_{\mathrm{thresh}}$ of $-1.901$ was used for plotting this figure.

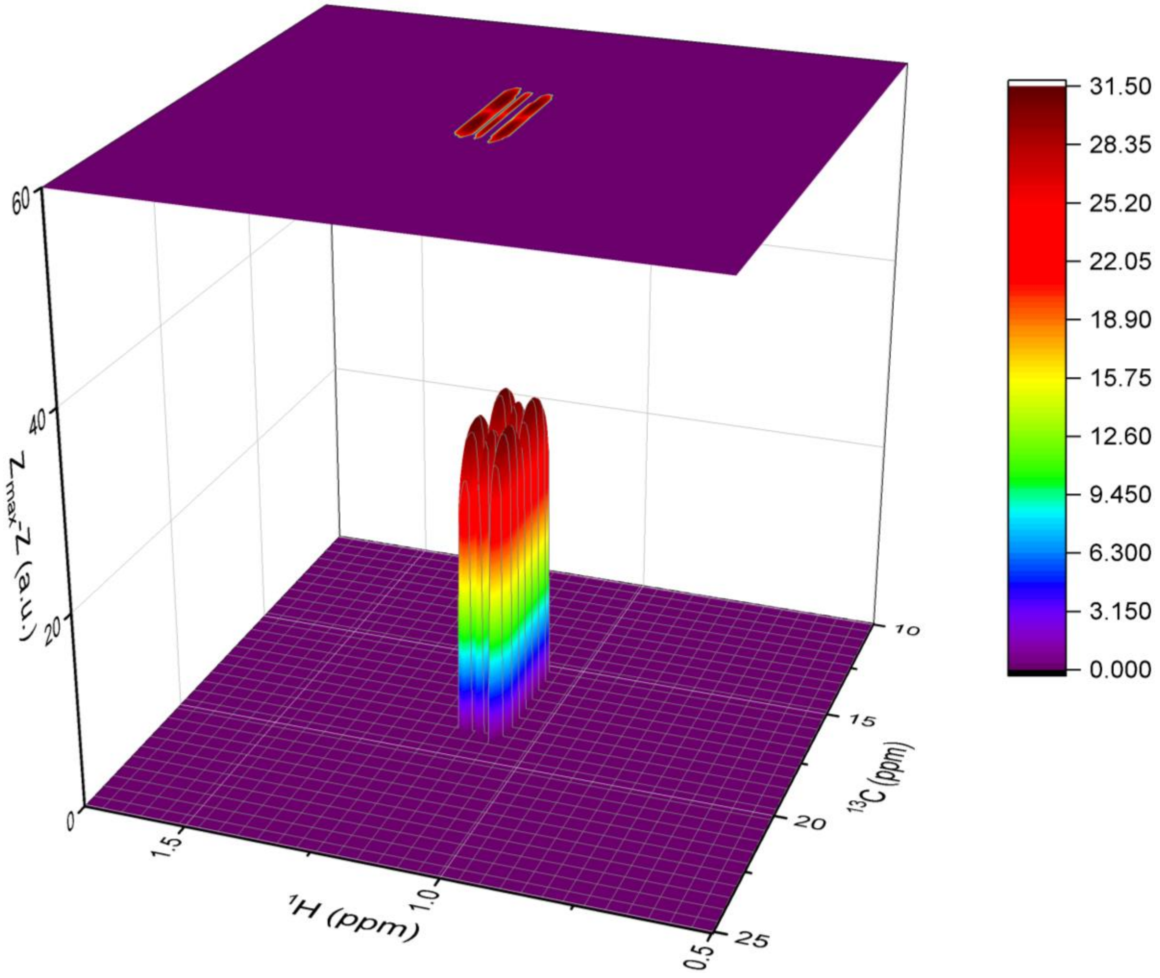


**Fig. S22.** 3D colormap surface plot of 2D HSQC data showing regions of $^{1}H$–$^{13}C$ correlations for the methyl group in the ethyl group of per-*O*-ethylated commercial iota-carrageenan after FDP–LCT along the $^{13}C$ dimension to the processed data shown in Fig. S21. The surface is generated from $Z_{max} - Z$, corresponding to $^{1}H$ and $^{13}C$ chemical shifts, with the color scale displayed on the right and the corresponding 2D contour map projection shown above. $Z_{max}$ is a constant set as 36.7368005696771, as detailed in Section 2 (Theory).

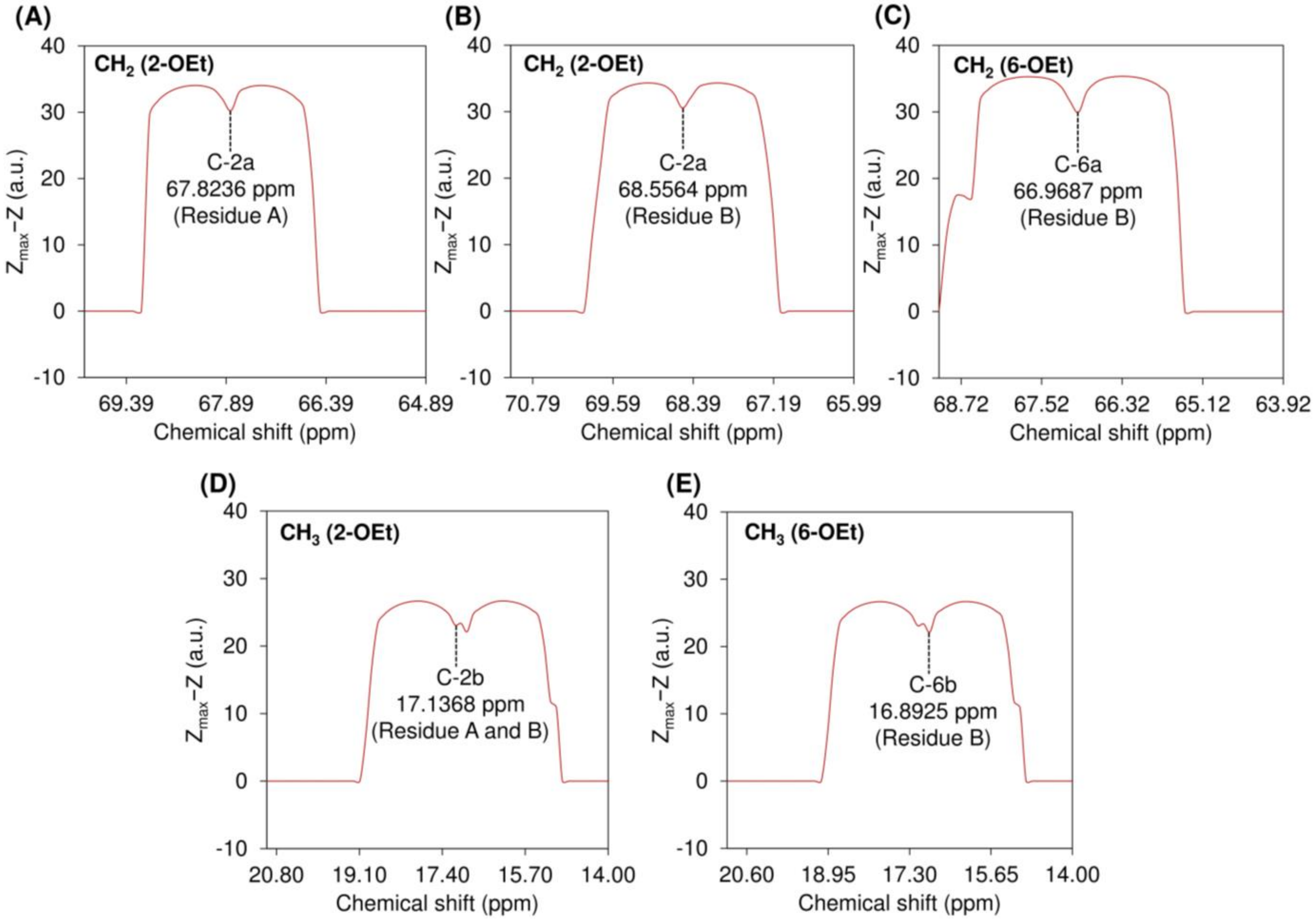


**Fig. S23.** Local resolution-boosted spectra of carbons on the methylene ($CH_2$) and methyl ($CH_3$) groups of the ethyl substituents at the *O*-2 and *O*-6 positions (designated 2-OEt and 6-OEt, respectively) of 4-linked 2-*O*-ethyl-3,6-anhydro-α-D-galactopyranose (residue A) and 3-linked 2,6-di-*O*-ethyl-4-*O*-sulfo-β-D-galactopyranose (residue B), extracted from 2D $^{1}H$–$^{13}C$ HSQC data of per-*O*-ethylated commercial kappa-carrageenan after FDP–LCT along the $^{1}H$ dimension followed by the $^{13}C$ dimension. $Z_{max}$ is a constant set as 36.7368005696771, as detailed in Section 2 (Theory).

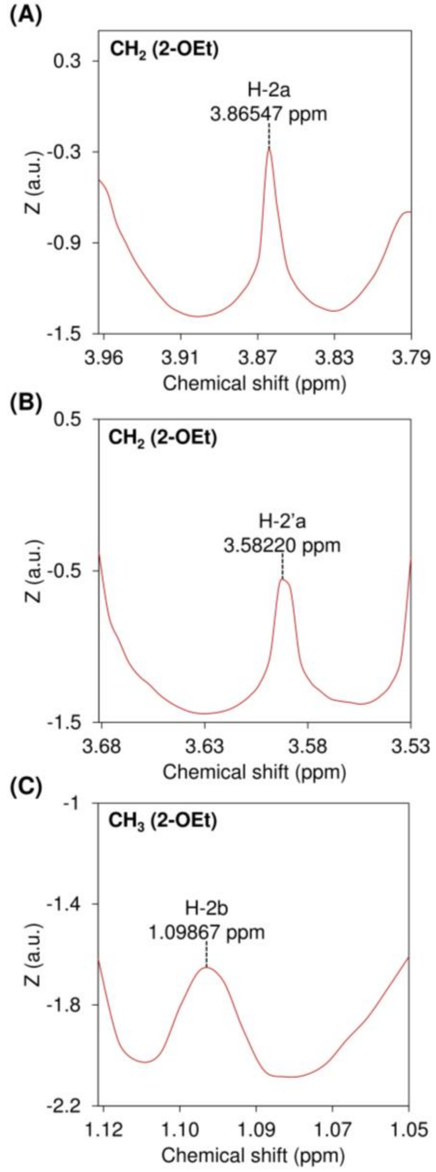


**Fig. S24.** Local resolution-boosted spectra of protons on the methylene ($CH_2$) and methyl ($CH_3$) groups of the ethyl substituents at the *O*-2 and *O*-6 positions (designated 2-OEt and 6-OEt, respectively) of 4-linked 2-*O*-ethyl-3,6-anhydro-α-D-galactopyranose (residue A), extracted from 2D $^{1}H$–$^{13}C$ HSQC data of per-*O*-ethylated commercial kappa-carrageenan after FDP–LCT along the $^{1}H$ dimension.

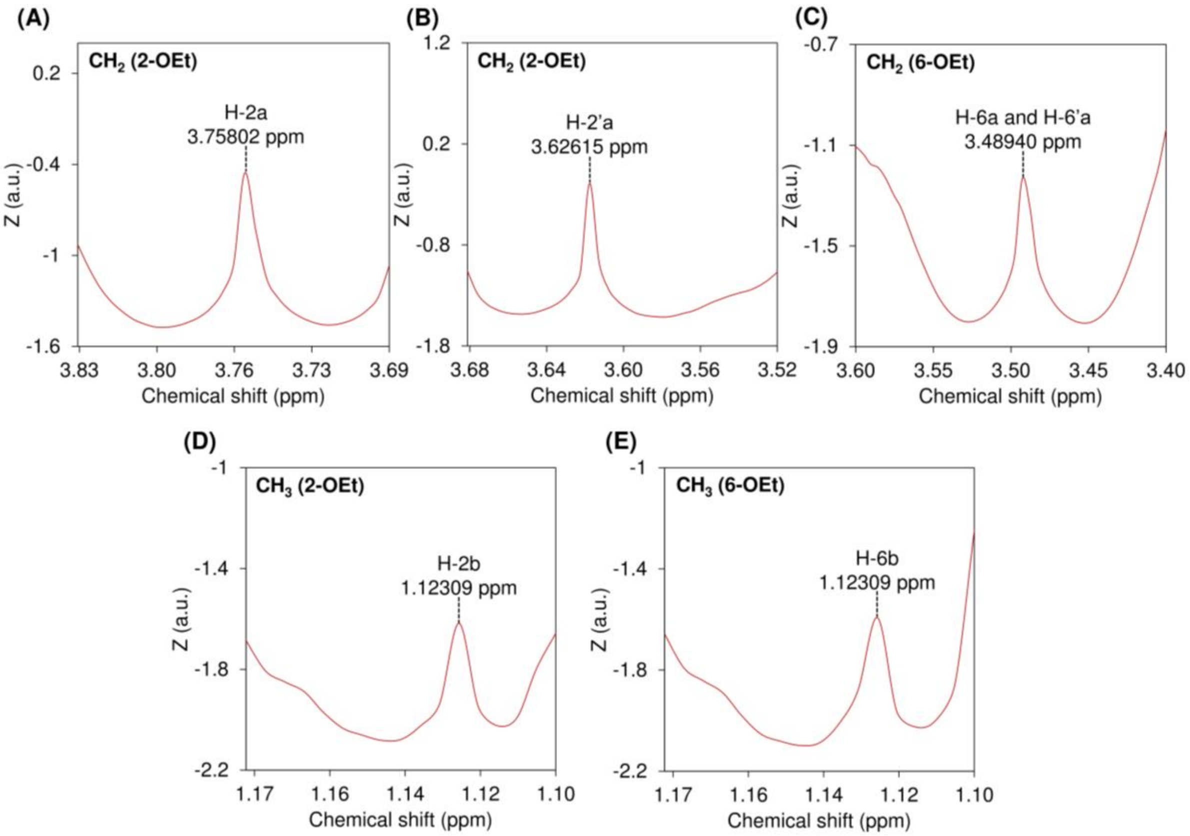


**Fig. S25.** Local resolution-boosted spectra of protons on the methylene ($CH_2$) and methyl ($CH_3$) groups of the ethyl substituents at the *O*-2 and *O*-6 positions (designated 2-OEt and 6-OEt, respectively) of 3-linked 2,6-di-*O*-ethyl-4-*O*-sulfo-β-D-galactopyranose (residue B), extracted from 2D $^{1}H$–$^{13}C$ HSQC data of per-*O*-ethylated commercial kappa-carrageenan after FDP–LCT along the $^{1}H$ dimension.

**Table S1.** Chemical shifts (unrounded values) of $^{1}H$ and $^{13}C$ nuclei in the sugar rings of per-*O*-ethylated carrageenans

| Per-*O*-ethylated carrageenan | Residues | | $^{1}H$ and $^{13}C$ chemical shifts (ppm) | | | | | | |
|---|---|---|---|---|---|---|---|---|---|
| | | | 1 | 2 | 3 | 4 | 5 | 6 | 6’ |
| Kappa | Residue **A** | $\delta_H$ | 5.07184 | 3.85570 | 4.18782 | 4.30992 | 3.60662 | 3.71407 | 3.55778 |
| | | $\delta_C$ | 94.8143 | 77.4719 | 80.0366 | 80.0366 | 74.7850 | 71.9760 | – |
| | Residue **B** | $\delta_H$ | 4.38319 | 3.10844 | 3.74337 | 4.39784 | 4.54436 | 3.71407 | 3.55778 |
| | | $\delta_C$ | 103.608 | 78.9374 | 77.9604 | 77.2276 | 71.7318 | 71.9760 | – |
| Iota | Residue **C** | $\delta_H$ | 5.22177 | 4.36747 | 4.59203 | 4.41141 | 3.56199 | 3.68404 | 3.55711 |
| | | $\delta_C$ | 93.3488 | 74.2660 | 79.5176 | 80.0366 | 74.6018 | 71.5791 | – |
| | Residue **D** | $\delta_H$ | 4.41141 | 3.21539 | 3.80120 | 4.44558 | 4.63109 | 3.68404 | 3.55711 |
| | | $\delta_C$ | 103.608 | 78.4184 | 77.3803 | 77.4108 | 72.3730 | 71.5791 | – |

Note: “–” means “not applicable”. Residues A and B represent 4-linked 2-*O*-ethyl-3,6-anhydro-α-D-galactopyranose and 3-linked 2,6-di-*O*-ethyl-4-*O*-sulfo-β-D-galactopyranose in per-*O*-ethylated kappa-carrageenan, respectively. Residues C and D represent 4-linked 2-*O*-sulfo-3,6-anhydro-α-D-galactopyranose and 3-linked 2,6-di-*O*-ethyl-4-*O*-sulfo-β-D-galactopyranose in per-*O*-ethylated commercial iota-carrageenan, respectively.

**Table S2.** Chemical shifts (unrounded values) of [1]H and [13]C nuclei in the methylene and methyl groups of the ethyl substituents in per-*O*-ethylated carrageenans

| Per-*O*-ethylated carrageenans | Residues | Ethylation position | | [1]H and [13]C chemical shifts (ppm) | |
|---|---|---|---|---|---|
| | | | | Methylene | Methyl |
| Kappa | Residue **A** | *O*-2 | $\delta_H$ | 3.58220, 3.86547 | 1.09867, 1.09867, 1.09867 |
| | | | $\delta_C$ | 67.8236 | 17.1368 |
| | Residue **B** | *O*-2 | $\delta_H$ | 3.62615, 3.75802 | 1.12309, 1.12309, 1.12309 |
| | | | $\delta_C$ | 68.5564 | 17.1368 |
| | | *O*-6 | $\delta_H$ | 3.48940, 3.48940 | 1.12309, 1.12309, 1.12309 |
| | | | $\delta_C$ | 66.9687 | 16.8925 |
| Iota | Residue **C** | – | $\delta_H$ | – | – |
| | | | $\delta_C$ | – | – |
| | Residue **D** | *O*-2 | $\delta_H$ | 3.60593, 3.73773 | 1.08847, 1.08847, 1.08847 |
| | | | $\delta_C$ | 68.5869 | 17.1062 |
| | | *O*-6 | $\delta_H$ | 3.48877, 3.48877 | 1.13729, 1.13729, 1.13729 |
| | | | $\delta_C$ | 66.9993 | 16.8009 |

Note: "–" means "not applicable". Residues A and B represent 4-linked 2-*O*-ethyl-3,6-anhydro-α-D-galactopyranose and 3-linked 2,6-di-*O*-ethyl-4-*O*-sulfo-β-D-galactopyranose in per-*O*-ethylated kappa-carrageenan, respectively. Residues C and D represent 4-linked 2-*O*-sulfo-3,6-anhydro-α-D-galactopyranose and 3-linked 2,6-di-*O*-ethyl-4-*O*-sulfo-β-D-galactopyranose in per-*O*-ethylated commercial iota-carrageenan, respectively.

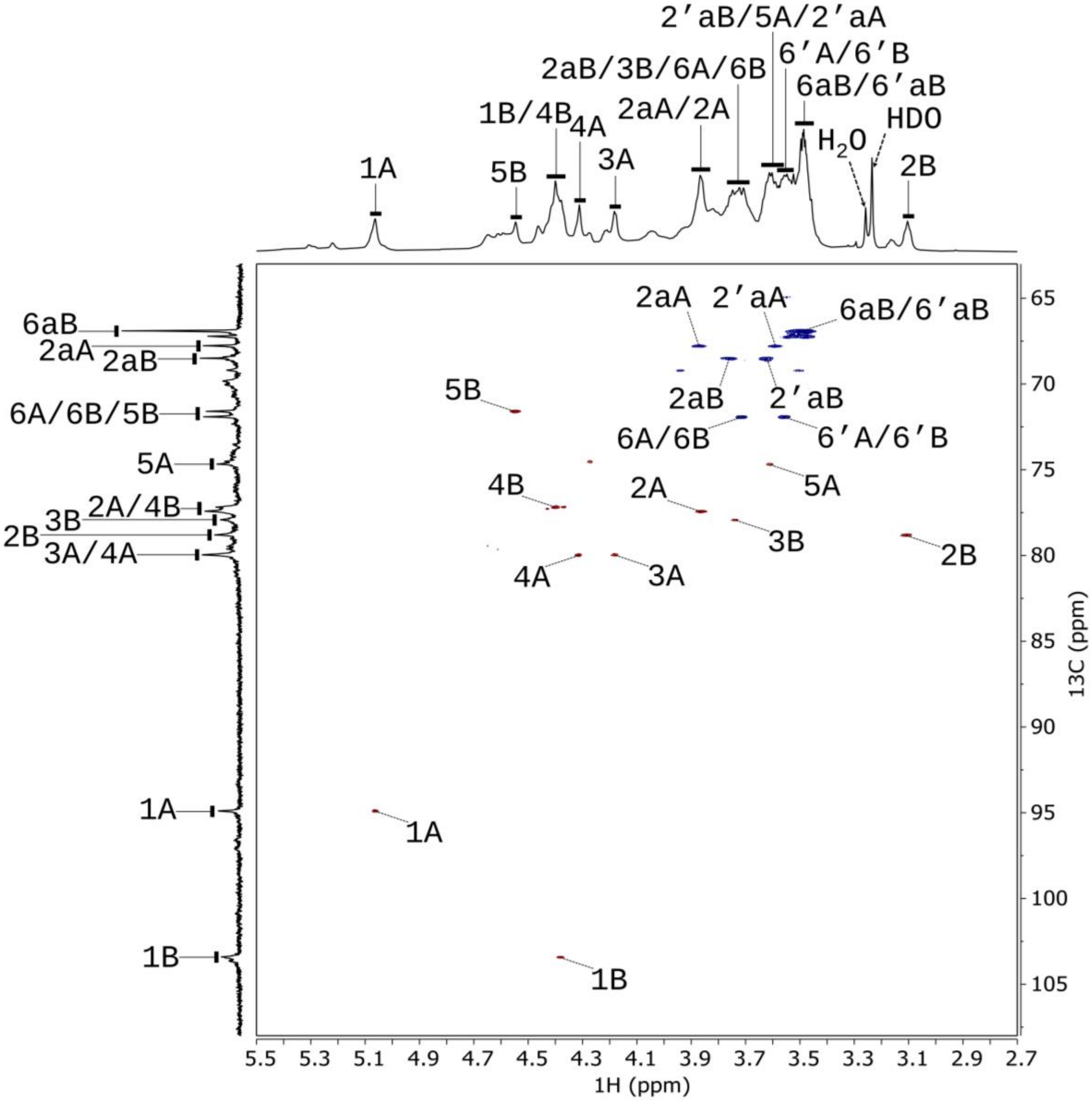


**Fig. S26.** 2D $^{1}H$–$^{13}C$ HSQC NMR spectrum (700 MHz, 323 K) of DMSO-$d_6$ solution of per-*O*-ethylated F60 isolated from *Mazzaella japonica*. Residues A and B represent 4-linked 2-*O*-ethyl-3,6-anhydro-α-D-galactopyranose and 3-linked 2,6-di-*O*-ethyl-4-*O*-sulfo-β-D-galactopyranose in per-*O*-ethylated kappa-carrageenan (major component), respectively. Cross peaks corresponding to correlations between directly bonded carbons and protons (one bond away) are marked. For example, the cross-peak "1A" refers to the correlation between C-1 and H-1 of residue A. The lowercase "a" indicates a signal from the methylene group of the ethyl group, and the number before it indicates the location of the ethyl group on the sugar ring. Signals separated by the solidus (/) symbol indicate overlapping. For example, "6A/6B" refers to the overlapping of signals 6A and 6B. $^{1}H$ and $^{13}C$ chemical shifts were internally referenced to TSP-$d_4$ as 0 ppm.

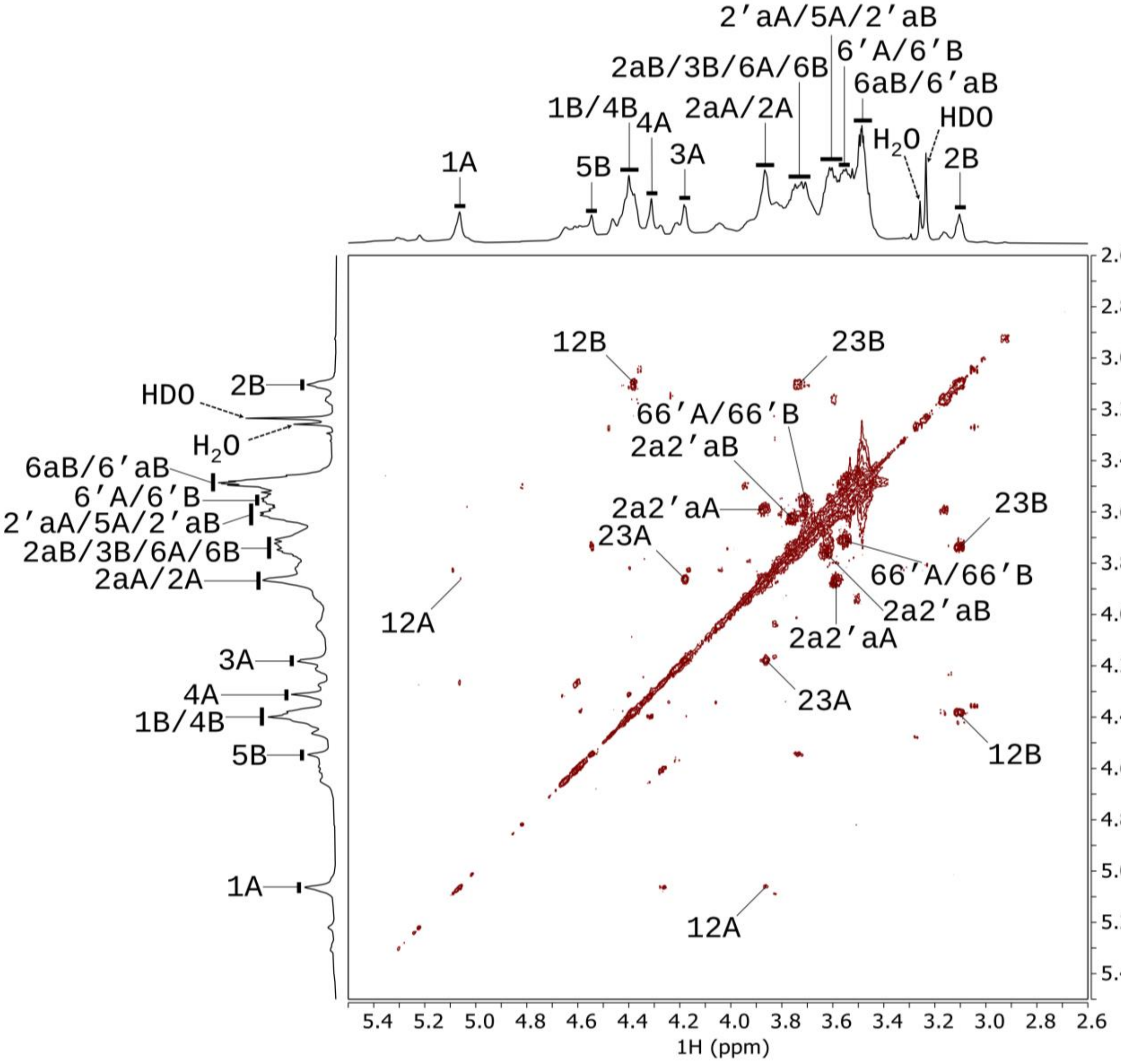


**Fig. S27**. 2D $^1$H–$^1$H COSY NMR spectrum (700 MHz, 323 K) of DMSO-$d_6$ solution of per-*O*-ethylated F60 isolated from *Mazzaella japonica*. Residues A and B represent 4-linked 2-*O*-ethyl-3,6-anhydro-α-D-galactopyranose and 3-linked 2,6-di-*O*-ethyl-4-*O*-sulfo-β-D-galactopyranose in per-*O*-ethylated kappa-carrageenan (major component), respectively. Cross peaks corresponding to correlations between neighboring protons are marked. For example, the cross-peak "12A" refers to the correlation between H-1 and H-2 of residue A. The lowercase "a" indicates a signal from the methylene group of the ethyl group, and the number before it indicates the location of the ethyl group on the sugar ring. Signals separated by the solidus (/) symbol indicate overlapping. For example, "66'A/66'B" refers to the overlapping of signals 66'A and 66'B. $^1$H chemical shifts were internally referenced to TSP-$d_4$ as 0 ppm.

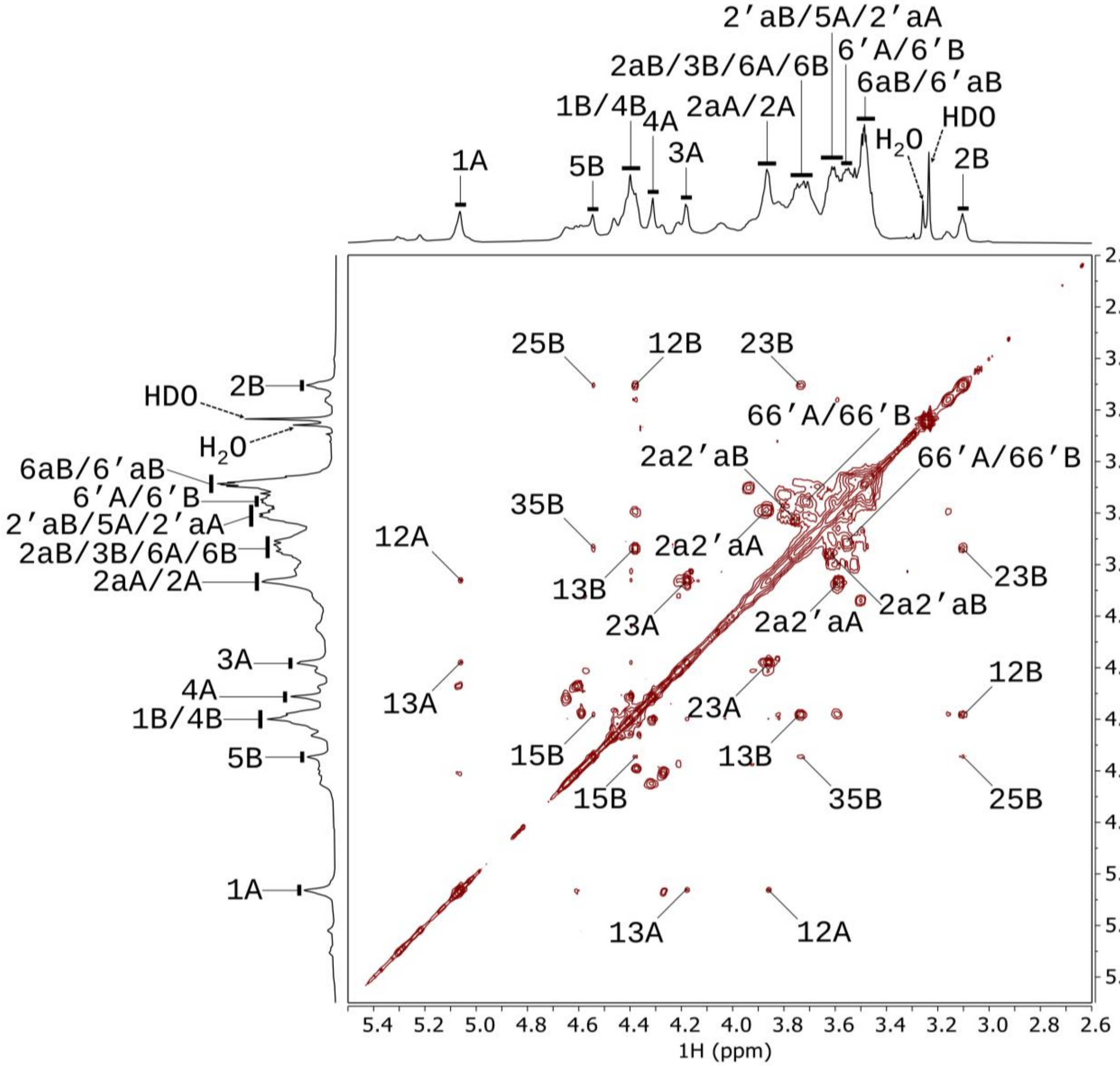


**Fig. S28**. 2D $^{1}$H–$^{1}$H TOCSY NMR spectrum (700 MHz, 323 K) of DMSO-$d_6$ solution of per-*O*-ethylated F60 isolated from *Mazzaella japonica*. Residues A and B represent 4-linked 2-*O*-ethyl-3,6-anhydro-α-D-galactopyranose and 3-linked 2,6-di-*O*-ethyl-4-*O*-sulfo-β-D-galactopyranose in per-*O*-ethylated kappa-carrageenan (major component), respectively. Cross peaks corresponding to correlations between neighboring protons are marked. For example, the cross-peak “13A” refers to the correlation between H-1 and H-3 of residue A. The lowercase “a” indicates a signal from the methylene group of the ethyl group, and the number before it indicates the location of the ethyl group on the sugar ring. Signals separated by the solidus (/) symbol indicate overlapping. For example, “66’A/66’B” refers to the overlapping of signals 66’A and 66’B. $^{1}$H chemical shifts were internally referenced to TSP-$d_4$ as 0 ppm.

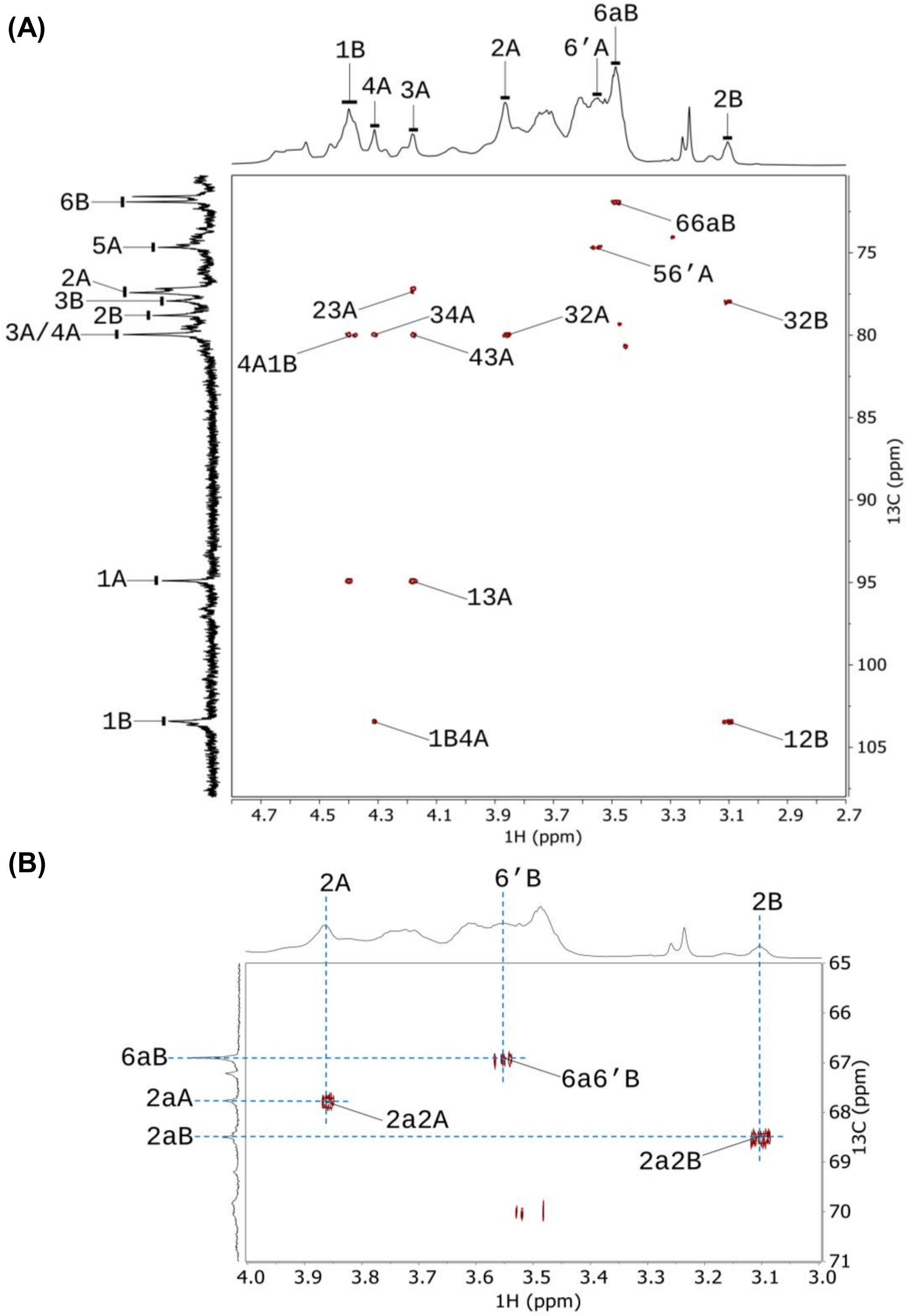


**Fig. S29**. Two regions of 2D $^{1}H$–$^{13}C$ HMBC NMR spectrum (700 MHz, 323 K) of DMSO-$d_6$ solution of per-*O*-ethylated F60 isolated from *Mazzaella japonica*. Residues A and B represent 4-linked 2-*O*-ethyl-3,6-anhydro-α-D-galactopyranose and 3-linked 2,6-di-*O*-ethyl-4-*O*-sulfo-β-D-galactopyranose in per-*O*-ethylated kappa-carrageenan (major component), respectively. $^{1}H$–$^{13}C$ cross-signals within the same sugar ring (intra-ring) and across glycosidic linkages (inter-ring) are marked. For example, in region A, "13A" refers to the intra-ring correlation between C-1 and H-3 of residue A, while “4A1B” indicates the inter-ring correlation between C-4 of residue A and H-1 of residue B. The lowercase “a” in both regions indicates a signal from the methylene group of the ethyl group, and the number before it indicates the location of the ethyl group on the sugar ring. $^{1}H$ and $^{13}C$ chemical shifts were internally referenced to TSP-$d_4$ as 0 ppm.

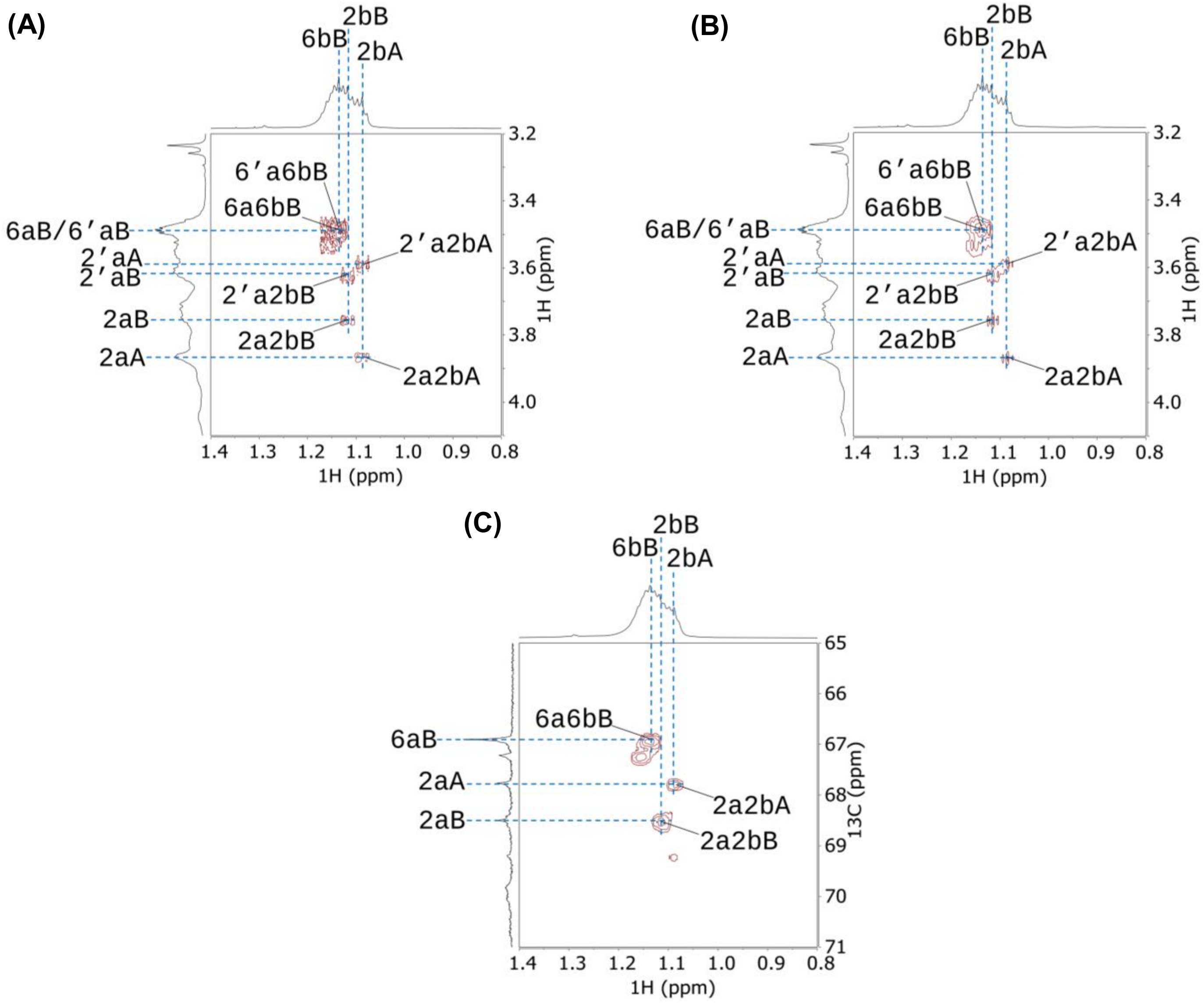


**Fig. S30**. Regions of (A) 2D $^1$H–$^1$H COSY, (B) 2D $^1$H–$^1$H TOCSY, and (C) 2D $^1$H–$^{13}$C HMBC spectra (700 MHz, 323 K) of DMSO-$d_6$ solution of per-*O*-ethylated F60 isolated from *Mazzaella japonica*, showing correlations between the atoms of the methylene group and the methyl group in the ethyl group attached to different positions on the sugar ring. Residues A and B represent 4-linked 2-*O*-ethyl-3,6-anhydro-α-D-galactopyranose and 3-linked 2,6-di-*O*-ethyl-4-*O*-sulfo-β-D-galactopyranose in per-*O*-ethylated kappa-carrageenan (major component), respectively. The lowercase "a" and "b" indicate relevant $^{1}H$ or $^{13}C$ signals from the methylene group and the methyl group of the ethyl group, and the number before "a" and "b" indicates the location of the ethyl group on the sugar ring. For example, the cross signals "2a2bA" in the COSY and TOCSY spectra indicate the correlation between the methylene proton and the methyl proton in the ethyl group attached to the *O*-2 position of residue A, while in the HMBC spectrum, it indicates the correlation between the methylene $^{13}C$ and the methyl proton in the ethyl group attached to the *O*-2 position of residue A. $^{1}H$ and $^{13}C$ chemical shifts were internally referenced to TSP-$d_4$ as 0 ppm.

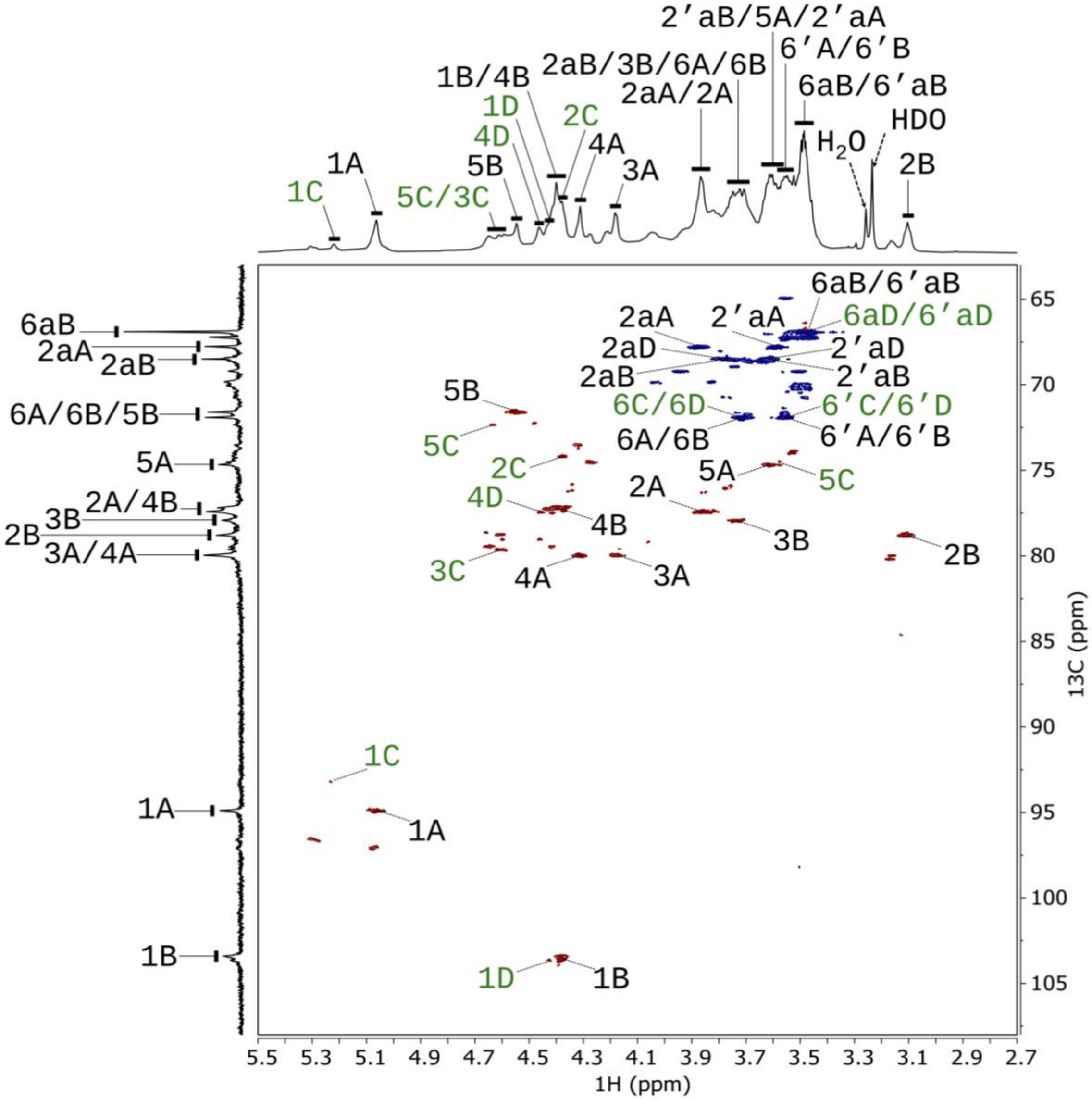


**Fig. S31**. 2D $^{1}$H–$^{13}$C HSQC NMR spectrum (700 MHz, 323 K) of DMSO-$d_6$ solution of per-*O*-ethylated F60 isolated from *Mazzaella japonica*, shown with a low contour-threshold display setting to reveal weaker cross-peaks. Residues A and B represent 4-linked 2-*O*-ethyl-3,6-anhydro-α-D-galactopyranose and 3-linked 2,6-di-*O*-ethyl-4-*O*-sulfo-β-D-galactopyranose in per-*O*-ethylated kappa-carrageenan (major component), respectively. Residues C and D represent 4-linked 2-*O*-sulfo-3,6-anhydro-α-D-galactopyranose and 3-linked 2,6-di-*O*-ethyl-4-*O*-sulfo-β-D-galactopyranose in per-*O*-ethylated iota-carrageenan (minor component). Labels corresponding to the minor iota-carrageenan structures are shown in green. Cross peaks corresponding to correlations between directly bonded carbons and protons (one bond away) are marked. For example, the cross-peak “1A” refers to the correlation between C-1 and H-1 of residue A. The lowercase “a” indicates a signal from the methylene group of the ethyl group, and the number before it indicates the location of the ethyl group on the sugar ring. Signals separated by the solidus (/) symbol indicate overlapping. For example, “6A/6B” refers to the overlapping of signals 6A and 6B. $^{1}$H and $^{13}$C chemical shifts were internally referenced to TSP-$d_4$ as 0 ppm.

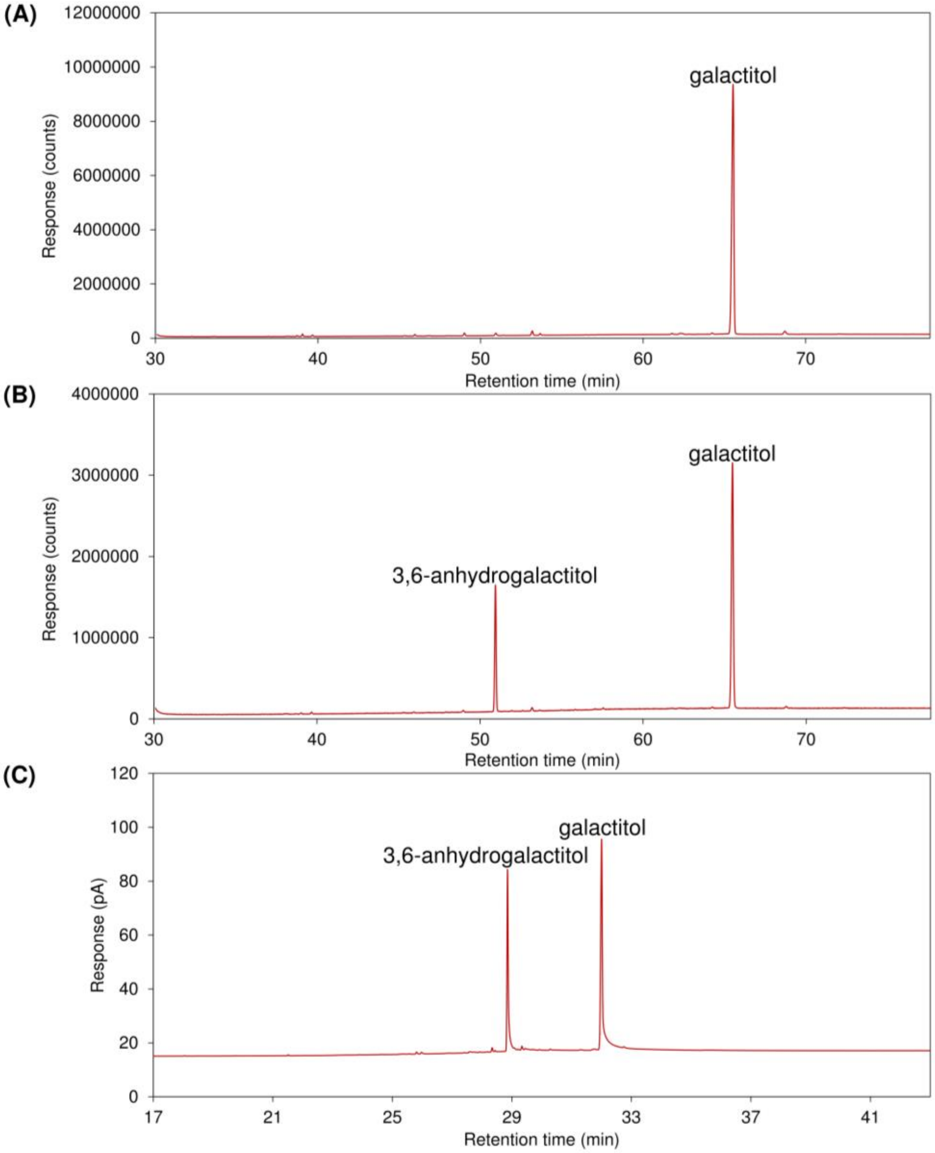


**Fig. S32**. GC–MS and GC–FID chromatograms of alditol acetates (AAs) prepared from the *Mazzaella japonica* F60 fraction, including (A) total ion current (TIC) chromatogram of AAs generated by depolymerization with 2 M TFA hydrolysis, showing a single dominant peak from acetates of galactitol; (B) TIC chromatogram and (C) FID chromatogram of AAs generated by depolymerization via reductive hydrolysis, showing peaks from galactitol and 3,6-anhydrogalactitol. Prior to depolymerization, the sulfated galactan was treated with carbodiimide activation at pH 4.75 followed by sodium borodeuteride ($NaBD_4$) to reduce and 6,6'-dideuterium-label any existing uronic acids, in order to confirm that the sample was not contaminated with uronic acids.

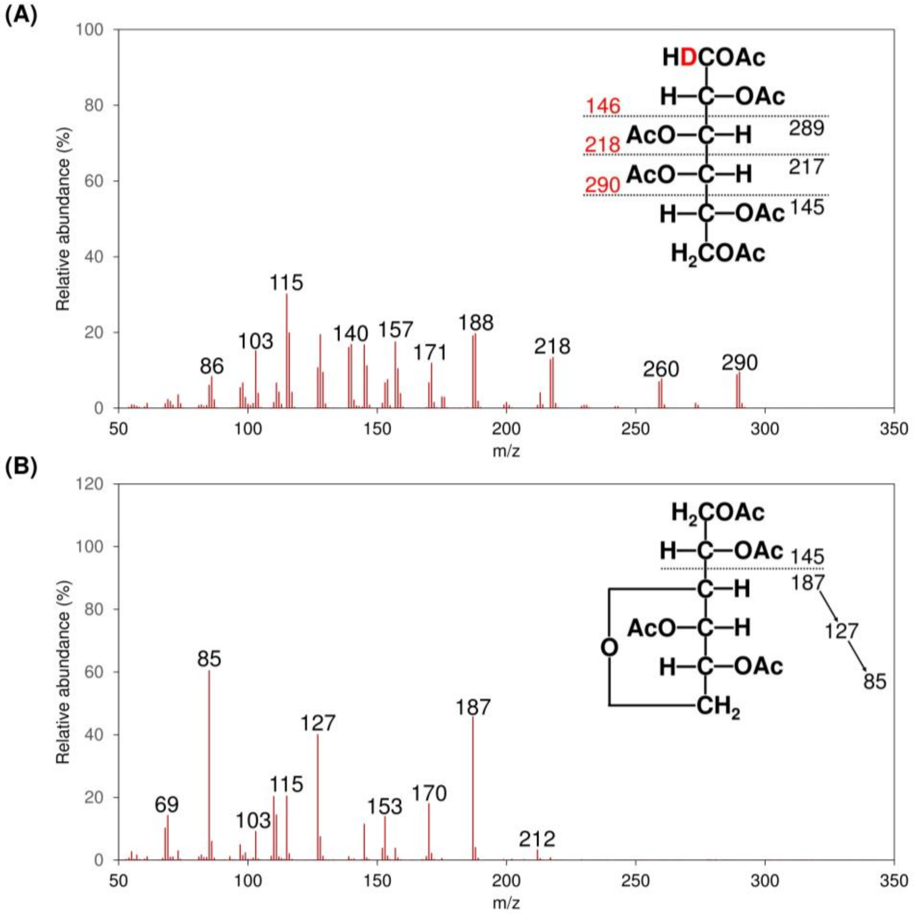


**Fig. S33.** Electron impact (EI)–MS spectra and fragmentation pattern assignments of AAs derived from (A) galactitol with deuterium labeling and (B) 3,6-anhydrogalactitol without deuterium labeling, extracted from the TIC chromatograms shown in Fig. S32A and S32B, respectively. In the annotated AA structure, *m/z* values of fragments carrying deuterium are highlighted in red, while those not carrying deuterium are shown in black.

**Table S3.** Monosaccharide composition (mol%) of the *Mazzaella japonica* F60 fraction determined by alditol acetate analysis.

| Monosaccharides | Experiment 1 | Experiment 2 | Mean |
|---|---|---|---|
| AnGal | 28.58 | 32.42 | 30.50 |
| Gal | 71.42 | 67.58 | 69.50 |
| 3-*O*-methyl Gal | t.r. | t.r. | t.r. |

Note: Two independent experiments were conducted on the sample, and the mean values were calculated from duplicate analyses. Gal and AnGal denote galactose and 3,6-anhydrogalactose residues, respectively. The prefix *O*-methyl in the monosaccharide name indicates a naturally occurring methyl substituent. "t.r." indicates trace amounts (mol% < 0.5).

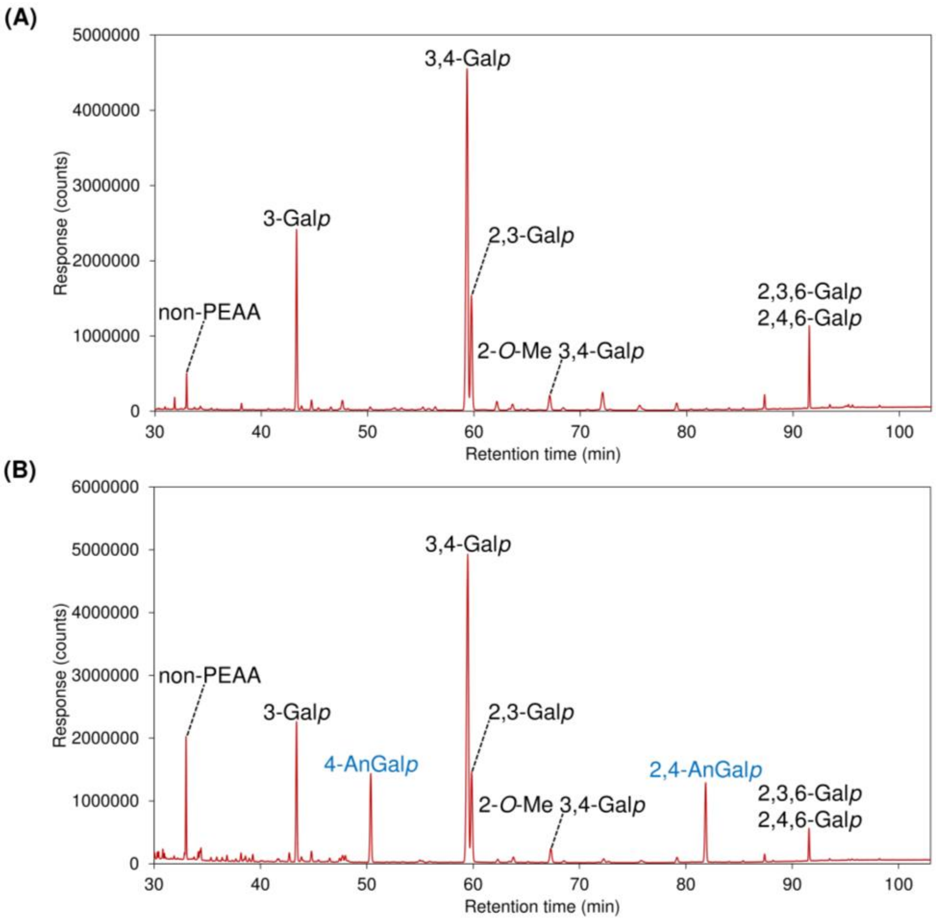


**Fig. S34.** GC–MS chromatograms of partially ethylated alditol acetates (PEAAs) prepared from the *Mazzaella japonica* F60 fraction, including (A) TIC chromatogram of PEAAs generated by depolymerization of the per-*O*-ethylated sample with 2 M TFA hydrolysis, showing signals from galactopyranose (Gal*p*) linkages but lacking signals from the acid-labile linkages of anhydrogalactopyranose (AnGal*p*); (B) TIC chromatogram of PEAAs generated by depolymerization via reductive hydrolysis, showing peaks from both Gal*p* and AnGal*p* linkages.

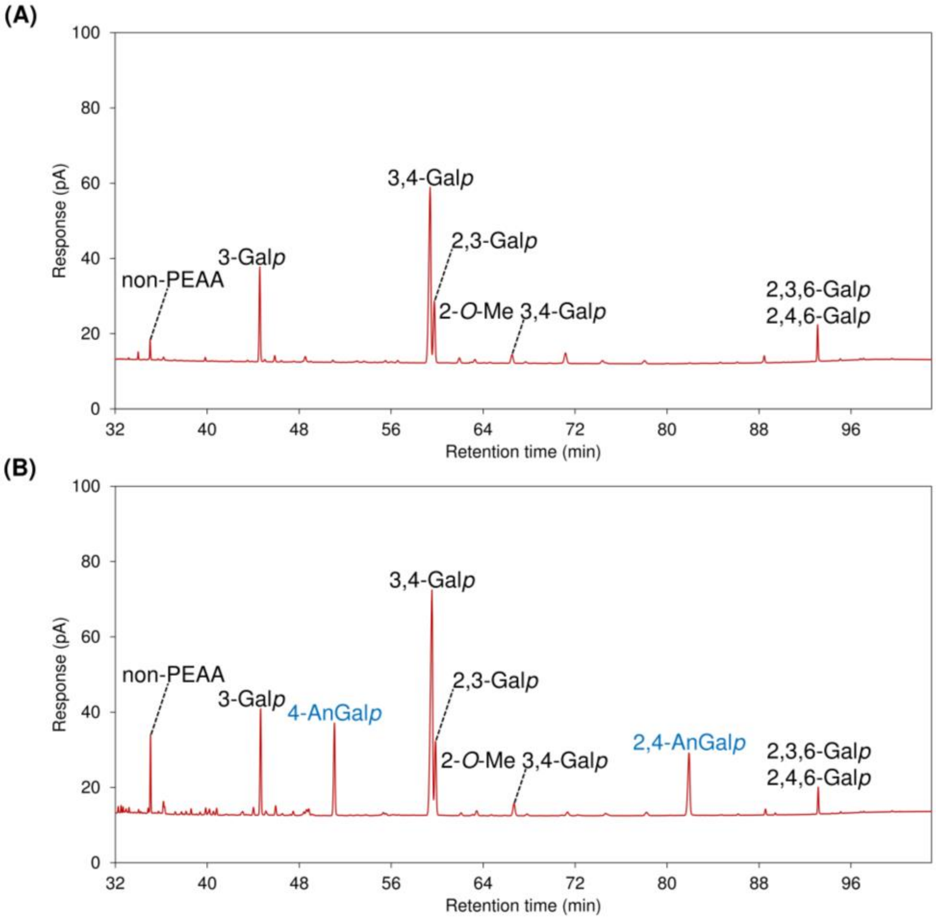


**Fig. S35.** GC–FID chromatograms of PEAAs prepared from the *Mazzaella japonica* F60 fraction by (A) depolymerization of the per-*O*-ethylated sample with 2 M TFA hydrolysis, showing signals from Gal*p* linkages but lacking those from the acid-labile AnGal*p* linkages, and (B) depolymerization via reductive hydrolysis, showing peaks from both Gal*p* and AnGal*p* linkages.

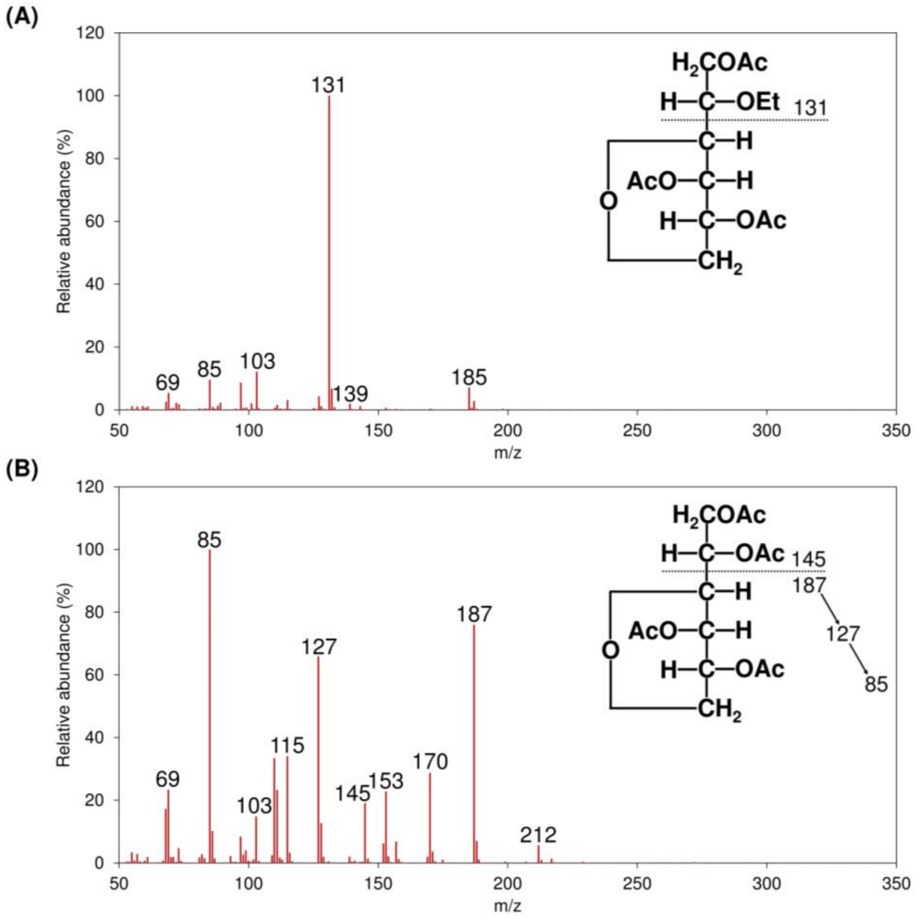


**Fig. S36.** EI–MS spectra and fragmentation pattern assignments of PEAAs from (A) 4-AnGal*p* and (B) 2,4-AnGla*p* without deuterium labeling, both extracted from the TIC chromatograms shown in Fig. S34B.

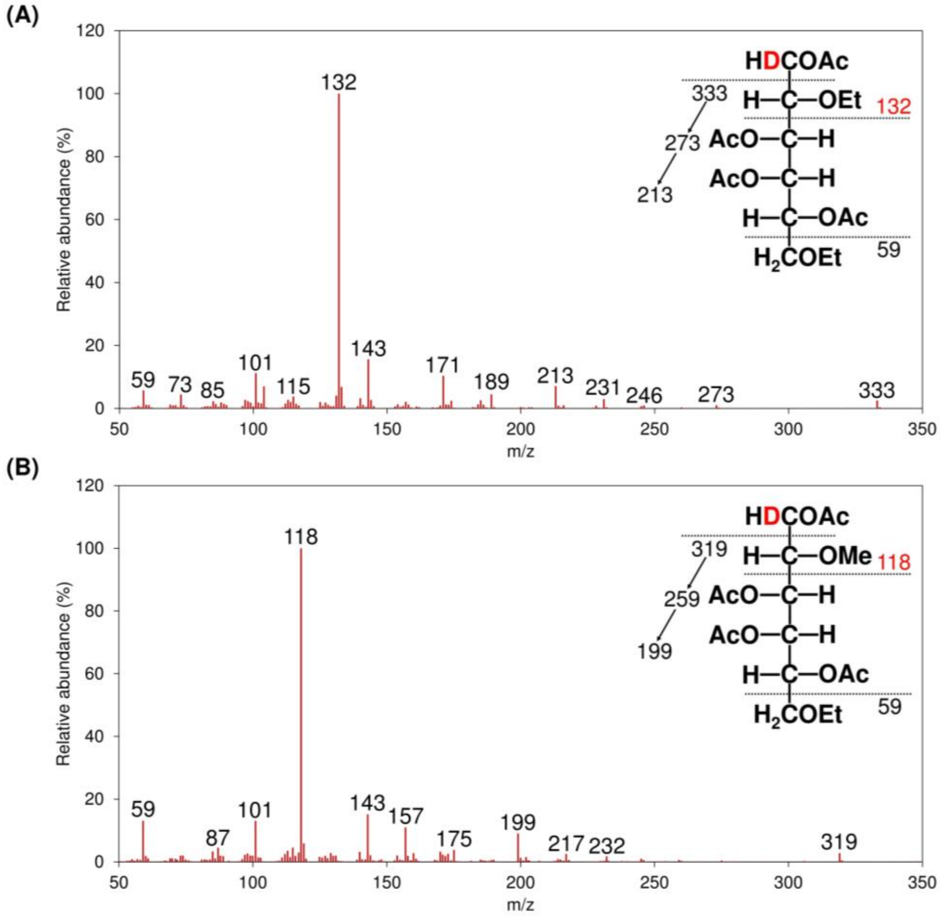


**Fig. S37.** EI–MS spectra and fragmentation pattern assignments of PEAAs derived from (A) 3,4-Gal*p* and (B) 2-*O*-methyl-3,4-Gla*p* with deuterium labeling, both extracted from the TIC chromatograms shown in Fig. S34A. In the annotated PEAA structure, *m/z* values of fragments carrying deuterium are highlighted in red, while those not carrying deuterium are shown in black.

**Table S4.** Linkage composition (mol%) of the *Mazzaella japonica* F60 fraction determined by per-*O*-ethylation–GC–MS analysis.

| Linkages | Experient 1 | Experiment 2 | Mean |
|---|---|---|---|
| t-Gal*p* | 0.44 | 0.42 | 0.43 |
| 2-Gal*p* | 0.25 | 0.23 | 0.24 |
| 3-Gal*p* | 9.87 | 9.38 | 9.63 |
| 2-*O*-methyl 3-Gal*p* | 0.31 | 0.30 | 0.31 |
| 4-*O*-methyl 3-Gal*p* | 0.66 | 0.68 | 0.67 |
| 6-*O*-methyl 3-Gal*p* | 0.16 | 0.16 | 0.16 |
| 4-Gal*p* | 0.69 | 0.70 | 0.69 |
| 2,3-Gal*p* | 10.04 | 9.77 | 9.90 |
| 4-*O*-methyl 2,3-Gal*p* | 0.38 | 0.35 | 0.37 |
| 6-*O*-methyl 2,3-Gal*p* | 0.21 | 0.19 | 0.20 |
| 2,4-Gal*p* | 0.18 | 0.17 | 0.18 |
| 3-*O*-methyl 2,6-Gal*p* | 0.18 | 0.17 | 0.17 |
| 3,4-Gal*p* | 34.15 | 34.40 | 34.28 |
| 2-*O*-methyl 3,4-Gal*p* | 2.01 | 1.99 | 2.00 |
| 6-*O*-methyl 3,4-Gal*p* | 0.80 | 0.76 | 0.78 |
| 3,6-Gal*p* | 0.78 | 0.78 | 0.78 |
| 4,6-Gal*p* | 2.41 | 2.43 | 2.42 |
| 2-*O*-methyl 4,6-Gal*p* | 0.11 | 0.09 | 0.10 |
| 3-*O*-methyl 4,6-Gal*p* | 0.13 | 0.13 | 0.13 |
| 2,3,6-Gal*p* | 1.11 | 1.09 | 1.10 |
| 2,4,6-Gal*p* | 2.61 | 2.63 | 2.62 |
| 3,4,6-Gal*p* | 0.93 | 0.80 | 0.87 |
| 2,3,4,6-Gal*p* | 0.19 | 0.19 | 0.19 |
| 4-AnGal*p* | 16.11 | 16.23 | 16.17 |
| 2,4-AnGal*p* | 15.29 | 15.97 | 15.63 |

Note: Two independent experiments were conducted on the sample, and the mean values were calculated from the duplicate analyses. Gal*p* and AnGal*p* denote galactopyranose and 3,6-anhydrogalactopyranose residues, respectively. The prefix *O*-methyl in the linkage name indicates a naturally occurring methyl substituent.
